# ROCK-mediated junctional remodelling preserves barrier function in a developing epithelium during hypoxia

**DOI:** 10.64898/2026.06.29.735176

**Authors:** Michelle Fernandes, Aman Kaushik, Mahendra Sonawane

## Abstract

Epithelia contribute to organismal homeostasis by functioning as selectively permeable barriers. Being avascular, epithelia are particularly vulnerable to hypoxia. Yet how developing epithelia adapt to preserve tissue integrity under hypoxic conditions remains elusive. Here, we show that in bilayered zebrafish epidermis hypoxia disrupts epithelial architecture reversibly, but preserves tight junction-mediated barrier function. ROCK maintains the tight apico-cortical localisation of active Non-muscle myosin-II (pNM-II) to mediate a squamous-to-cuboidal shape transition in epithelial cells under hypoxia. It further suppresses Crb3-dependent local force imbalance that otherwise results in cell delamination and cell-rosette formation. Importantly, we identify ROCK as a regulator that promotes barrier function maintenance specifically under hypoxia. Our genetic analyses demonstrate that ROCK primarily coordinates junctional remodelling to maintain barrier function during hypoxia while actomyosin contractility may foster the junctions secondarily. We propose that under an energy constrained hypoxic environment, dismantling E-cadherin based adhesion and augmenting ROCK-dependent barrier function in a developing epithelium ensures organism survival allowing recovery when normoxic conditions resume.

## Introduction

In metazoans, epithelia form the lining of the body and internal surfaces. These tissues are made up of epithelial cells -having defined morphologies such as squamous, columnar, cuboidal, etc.-that are arranged in mono- or multi-layered sheets held together by cell-cell and cell-matrix adhesions. The vertebrate epidermis is a stratified epithelium, which develops from a bi-layered epithelium consisting of an outer periderm and an underlying basal epidermis. The epidermis, whether during development or in adults, primarily acts as a selectively permeable barrier and thus, segregates the organism’s interior milieu from the exterior. In terrestrial adult vertebrates, along with the stratum corneum, the tight junctions contribute significantly to the maintenance of the epidermal barrier function thereby, preventing water loss and controlling pathogen entry <u>(Morita, K., et al 2011)</u>. Additionally, the epidermis exhibits other types of cell adhesion mechanisms such as desmosomes, hemidesmosomes, and adherens junctions, that in conjunction with the associated cytoskeleton, participate in providing mechanical support and regulating tissue morphogenesis <u>(Sumigray, K.D. and Lechler, T., 2015, McGrath, J.A. and Uitto, J., 2023)</u>.

Cellular junctions and associated cytoskeleton are essential for epithelial sheets to act as a continuum. The formation of these junctions, especially the tight junctions and adherens junctions, are spatiotemporally determined in concert by biochemical and biomechanical signalling <u>(Garcia, M.A., et al 2018, Mira-Osuna, M. and Borgne, R.L., 2024)</u>. Amongst the biochemical signals, regulators of apicobasal cell-polarity, such as the apically localized Crumbs complex, are essential in the formation and positioning of the cellular junctions <u>(Feigin, M.E.</u> <u>and Muthuswamy, S.K., 2009, Buckley, C.E. and St Johnston, D., 2022)</u>. Further, the formation and maintenance of cell-cell adhesion requires the presence of contractile actomyosin regulated by the activity of RhoGTPases at the cell cortex <u>(Bertet, C., et al 2004, Itoh, M. et al, 2012, Citi,</u> <u>S., et al 2014, Zihni, C. and Terry, S.J., 2015)</u>. Importantly, Crumbs is also known to determine RhoGTPase localisation to the apical domain to regulate cytoskeletal organisation associated with the apical junctions <u>(Roh, M.H., et al 2003, Lemmers, C., et al 2004, Flores-Benitez, D. and</u> <u>Knust, E., 2015, Silver, J.T., et al 2019)</u>. Ultimately, the balance between inward contractile forces generated by actomyosin and relaxing adhesive forces are essential to maintain the epithelial architecture. Under different physiological contexts, force imbalance generated by effective directional increase in actomyosin contractility at the cellular level results in tissue shape changes or even extrusion of cells from the sheet <u>(Bertet, C., et al 2004, Rosenblatt, J., et</u> <u>al 2001, Nishimura, T. and Takeichi, M., 2008, Martin, A.C., et al 2009, Van Itallie, C.M., et al</u> <u>2009, Martin, A.C., et al 2010, Krueger, D., et al 2018, Okuda, S. and Fujimoto, K., 2020, Chouhan, G., et al 2024)</u>.

Though our understanding of the regulation of junction formation and maintenance by polarity modules under different physiological contexts is primarily derived from simple epithelia, several of these regulators have conserved functions in a stratified epithelium like the epidermis <u>(Helfrich, I., et al 2007, Muroyama, A. and Lechler, T., 2012, Ali, N.J., et al 2016, Arora, P., et al</u> <u>2020)</u>. Similarly, the RhoGTPases participate in tight junctional remodelling (formation and maintenance) in the epidermis, however the mechanistic details remain underexplored <u>(Slattum,</u> <u>G., et al 2009, Zhang, M., et al 2019, Cho, Y., et al 2025)</u>.

Epithelia are avascular tissues. They rely on diffusion for their oxygen requirements and are, therefore, susceptible to fluctuating oxygen levels. This susceptibility is profound in epithelia that are responsible for oxygen uptake and distribution to the entire organism like the alveolar and the gill epithelium, with prolonged or severe hypoxia impacting their ability to regulate tissue integrity <u>(Jain, M. and Sznajder, J.I., 2005, Bouvry, D., et al 2006, Nilsson, G.E., et al</u> <u>2012, Jonz, M.G., et al 2016, Prochazkova, K. and Uhlík, J., 2024, Glover, L.E. and Colgan, S.P.,</u> <u>2017, Song, H.A., et al 2017, Wong, S.L., et al 2023. Huang, J., et al 2025)</u>. The epidermis is also known to acquire oxygen from the surrounding environment for its sustenance and is consequently, shown to persist under mild hypoxic conditions <u>(Stücker, M., et al 2002, Heise,</u> <u>H.M., et al 2003, Rezvani, H.R., et al 2011)</u>. Besides, during early development, when the lungs or gills are not formed, the epidermis is one of the major sources of oxygen uptake in several vertebrates <u>(Feder, M.E. and Burggren, W.W., 1985, Burggren, W.W. and Pinder, A.W., 1991, Seymour, R.S., 1999)</u>. Yet it has remained unclear whether and how hypoxic conditions affect the barrier function and tissue homeostasis in the developing epidermis.

Here we show that in the embryonic zebrafish epidermis, a bilayered epithelium responsible for oxygen uptake <u>(Rombough, P., 2002)</u>, exposure to hypoxia results in a change in cell morphology and reduction in E-cadherin levels leading to cell adhesion defects. However, the barrier function remains intact under hypoxic conditions. Our results further demonstrate that RhoA-ROCK signaling remodels epidermal tight junctions and the associated active Non-muscle Myosin-II (pNM-II) to maintain the barrier function and consequently alter cell morphology. The higher recruitment of pNM-II downstream of ROCK signaling is essential to offset the local heterogeneities in force imbalances to prevent Crumbs-dependent delamination and death of the outer peridermal cells. Our results suggest that reinforcing the barrier function, at the expense of minimising other cell adhesion mechanisms, is an effective survival strategy in a hypoxic environment. This aids in animal survival, allowing the epithelium to recover when conditions are favourable.

## Results

### A physical hypoxia treatment of 24 hours results in a depletion of the oxygen level in the epidermal cells

To systematically examine the effects of physical hypoxia on epithelial architecture, we modified the previously published protocol in zebrafish <u>(Kamei, H. and Duan, C., 2018)</u>. We used T-75 cell culture flasks, which allowed us to maintain an isolated environment with controlled dissolved oxygen (DO) concentration of 2-3mg/L as the hypoxia treatment and DO of 6-8mg/L as the normoxic control (see methods for details). To begin with, we asked if a 24-hour long exposure to hypoxia using this method would deplete oxygen at the cellular level. To test this, we exposed wild type embryos at 24 hours post fertilization (24hpf) to hypoxia for 24 hours and incubated the embryos in a live cell oxygen sensor, ImageiT^TM^ Red Hypoxia Reagent. At 24hpf, the developing epidermis is a bilayered tissue with an outer periderm in contact with the external environment and the basolateral domain of the peridermal cells adhering to the underlying basal epidermis, which is in contact with the basal lamina via cell-matrix adhesion (Fig. 1A). We observed increased cytoplasmic intensity of the sensor, which indicates lower oxygen levels in the periderm of the hypoxia treated embryos relative to their normoxic controls, establishing the effectiveness of hypoxia treatment regime used (Fig. 1B). We further characterized the overt developmental phenotypes at the end of the treatment. The hypoxic embryos displayed morphological phenotypes such as a larger otic vesicle length (OVL), increased head-trunk angle (HTA), reduction in yolk consumption, reduced pigmentation and a smaller head size relative to their 24 hour normoxic controls (Fig. 1C), suggesting that the development of certain tissues and organs is delayed. In this and subsequent experiments presented in the sections below, in addition to 48hpf, we kept embryos at 36hpf (Stage matched 1) and 42hpf (Stage matched 2) as controls to assess the phenotypes that may arise due to the developmental delay under hypoxia.

**Figure 1:**
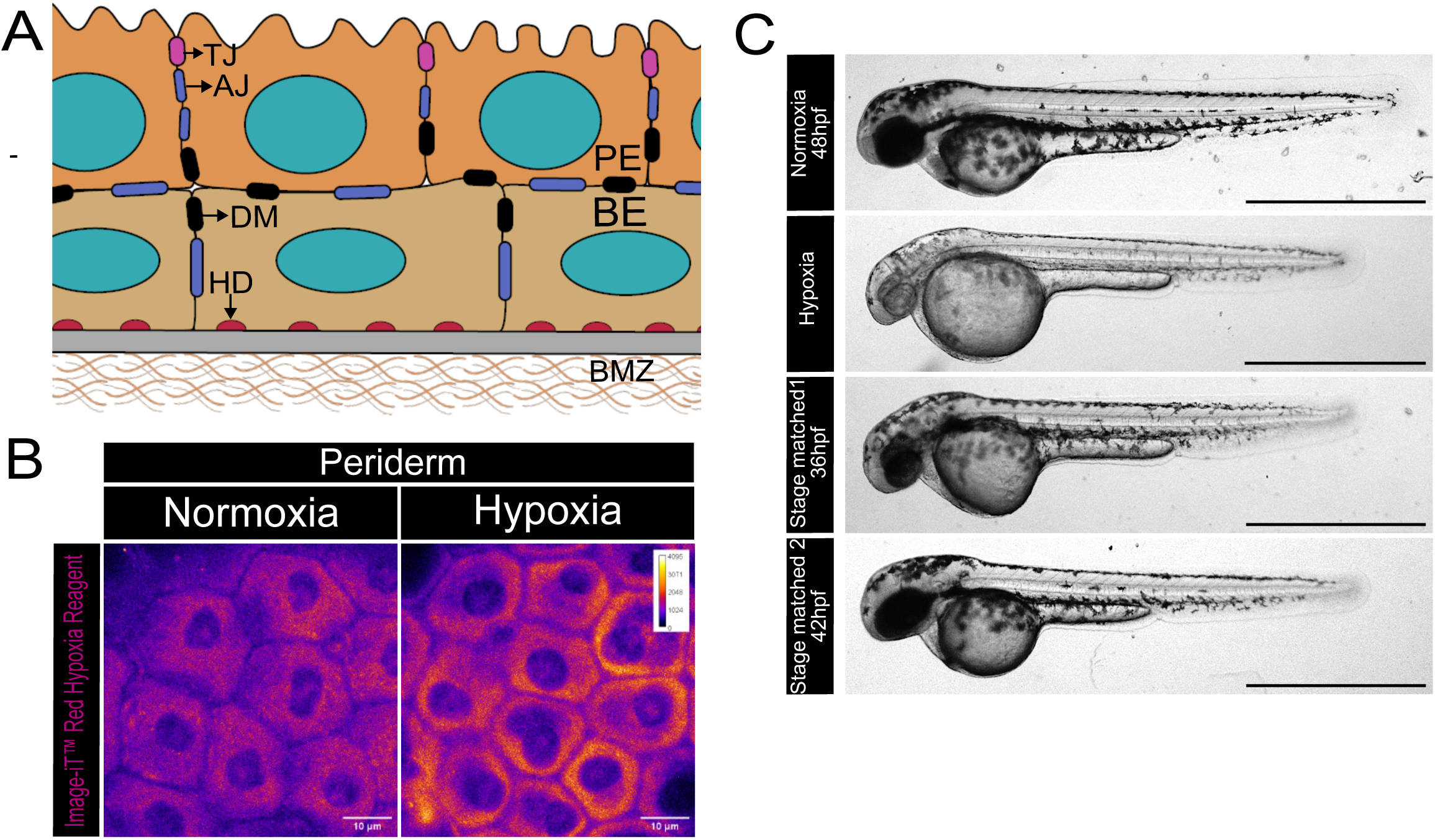
Assessment of hypoxia induced morphological changes in the developing zebrafish embryos. (A) Schematic representation of the zebrafish bi-layered embryonic epidermis (B) Confocal images of peridermal cells of normoxic and hypoxic embryos incubated with Image-iT^TM^ Red Hypoxia Reagent (Number of cells assessed for each condition=18, from experimental sets N=3). Note that the intensity of the dye is higher in the hypoxia treated embryos. Data represented in 12 bit-depth. (C) Representative brightfield images of normoxia (48hpf) and hypoxia treated embryos along with stage matched 1 (36hpf) and stage matched 2 (42hpf) normoxic embryos. Scale bars= 10µm in (B) and 1mm in (C). PE, periderm; BE, basal epidermis; TJ, tight junctions; AJ, Adherens junctions; DM, desmosomes; HD, hemi-desmosomes; BMZ, basement membrane zone.

To conclude, a 24-hour treatment with 2-3mg/L concentration of dissolved oxygen is sufficient to induce hypoxia at the cellular level in the developing epidermis. However, it did not result in any overt morphological phenotype in the epidermis.

### Hypoxic treatment results in disruption of the epithelial architecture without impacting tight junctions

To assess the effects of hypoxia on epithelial architecture and integrity, we used the *Tg(cldnB:lyn-EGFP)* line which fluorescently labels the peridermal cell membrane <u>(Haas, P. and</u> <u>Gilmour, D., 2006, Sonal, et al 2014)</u>. We observed a significant reduction in the apical cross-sectional area (Fig. 2A,B) and total surface area (Fig. 2C) in the peridermal cells of the hypoxia treated embryos as compared to the control normoxic embryos at 48hpf. In addition, there was an increase in cell height (Fig. 2E,F) but the basolateral surface area remained unaltered (Fig. 2D) at the end of a 24-hour treatment. The effect on the apical cell perimeter (Fig. 2A; Supp. Fig. 1A) and cell height (Fig. 2E; Supp. Fig. 1B) of the hypoxia treated embryos were different from all observed controls. These results indicate a transition of the peridermal cell morphology from squamous to cuboidal. Notably, this change in cell morphology was also accompanied by membrane gaps along the basolateral domain (Fig. 2H) of the peridermal cells but not in the apico-lateral domain (Fig. 2G). We further analysed the epithelial architecture in the head epidermis using Transmission Electron Microscopy (TEM), which confirmed the presence of intercellular gaps in the basal region of the peridermal cells (Fig. 2I). In addition, the TEM analysis also revealed a reduction in the thickness of basal lamina (Fig. 2I) indicating an impact of hypoxia on the basement membrane organisation.

**Figure 2:**
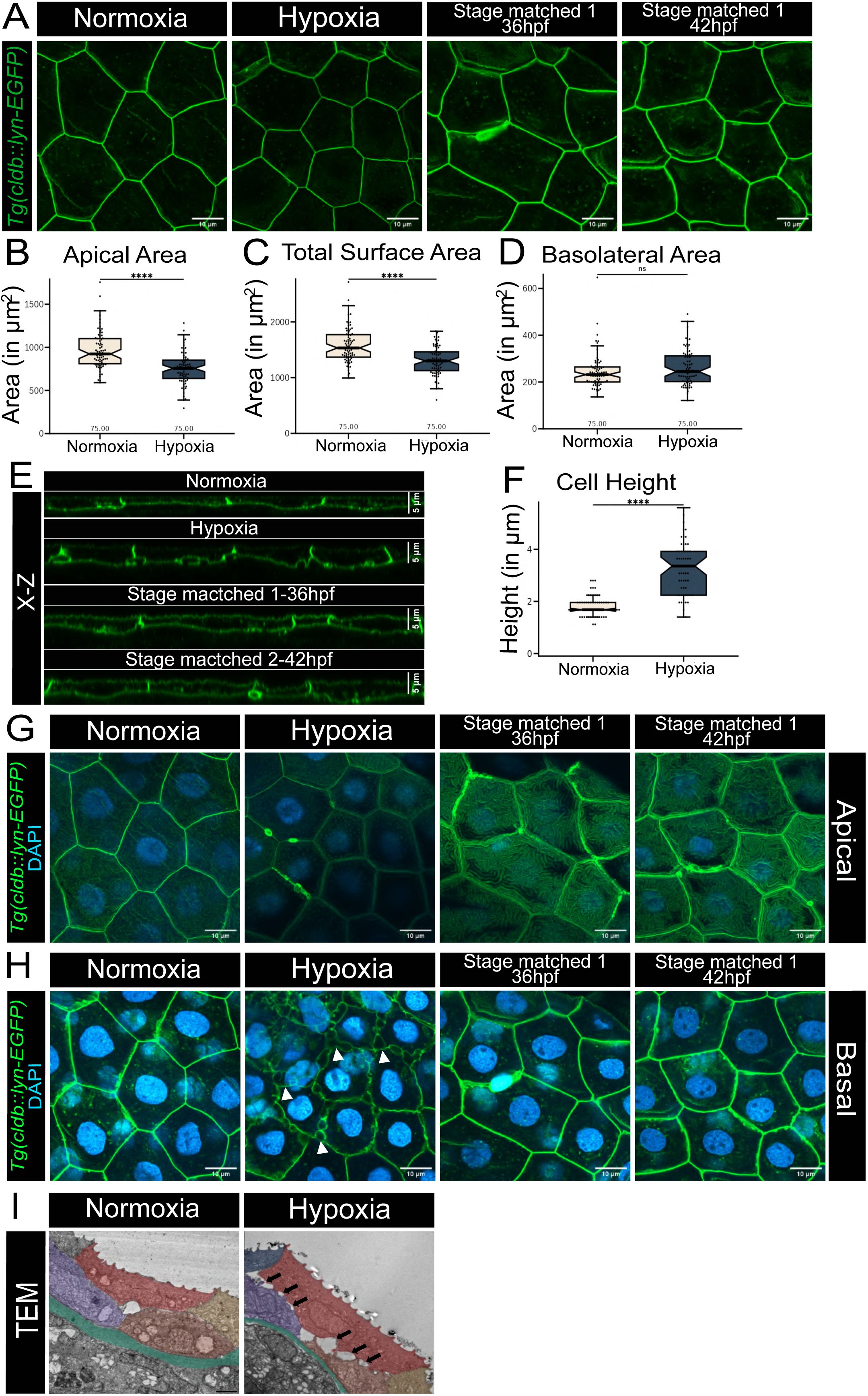
Epithelial architectural changes upon exposure to hypoxia. (A,E) Representative confocal images (A) and orthogonal (X-Z) sections (E) of peridermal cells in *Tg(cldnB:lynEGFP)* transgenic line exposed to normoxia (48hpf) and hypoxia along with the normoxic stage matched control 1 (36hpf) and stage matched control 2 (42hpf) embryos. (B,C,D,F) Estimation and comparison of apical (B), total (C) and basolateral surface area (D), and the cell height (F) under normoxic and hypoxic conditions. Data are median±interquartile range (no of cells assessed for each condition=75, N=3). \*\*\*\**P*<0.0001, n.s. not significant *P*>0.05 [Mann–Whitney U test]. (G,H) Confocal images of the apical (G) and basolateral slices (H) of the same peridermal cells with the membrane and the nucleus marked with Lyn-GFP and DAPI, respectively, under the conditions mentioned. White arrowheads mark the visible gaps in the basolateral domain of the hypoxia treated embryos. (I) Electron micrographs of the head epidermis of normoxia (48hpf) and hypoxia treated embryos. Peridermal cells false-coloured in red and yellow, basal epidermal cells false-coloured in purple and brown. Black arrowheads mark the intercellular gaps in the epidermis of hypoxia treated embryos. Green watershade marks the basement membrane zone. Scale bars= 10 µm in (A,G,H), 5µm in (E) and 2µm in (I).

Given the presence of intercellular gaps within the peridermal cells and between the two layers, we asked whether the E-cadherin localisation is perturbed under hypoxia treatment. E-cadherin is localised in a polarised fashion in the embryonic epidermis, where it displays increasing levels from the apical to basal, along the lateral domain, in the peridermal cells <u>(Arora, P., et al 2020)</u>. We used the previously reported method of assessing E-cadherin localisation <u>(Arora, P., et al</u> <u>2020</u>, see methods for details) in the normoxic versus hypoxic peridermal cells to gauge the impact of hypoxia on the polarised distribution of E-cadherin. We observed a uniform distribution of E-cadherin throughout the lateral domain (Fig. 3A) indicating clear loss of E-cadherin polarity (Fig. 3 B, C) and a reduction in average levels of E-Cadherin (Supp. Fig. 1G) following hypoxia treatment at 48hpf. While E-cadherin displayed a loss of polarized localisation in the hypoxic embryos relative to the stage matched controls as well (Supp. Fig. 1C, D, E, F), the average levels were not significantly different (Supp. Fig. 1G).

**Figure 3:**
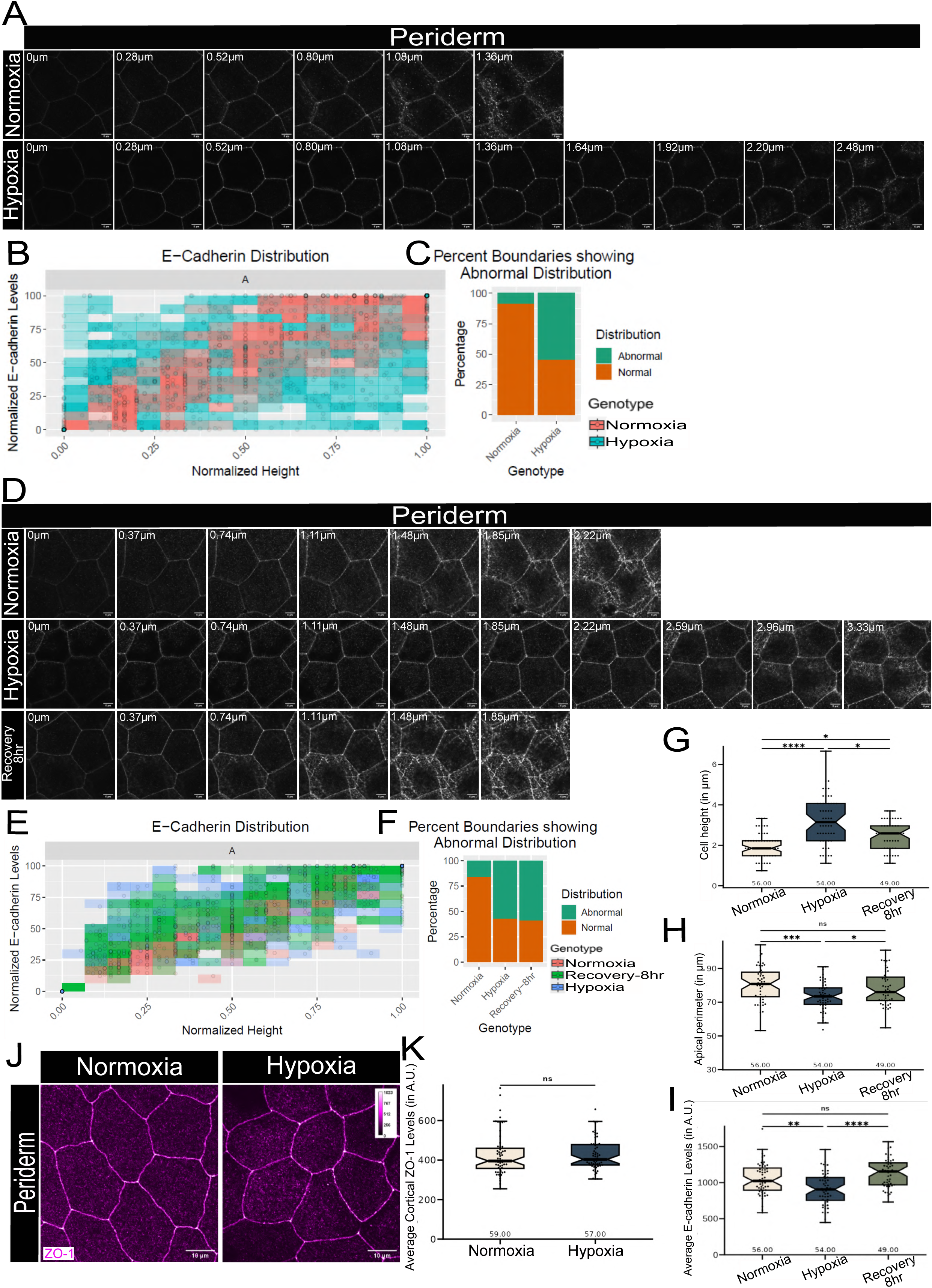
Under hypoxia, polarised distribution of E-cadherin is lost but tight junctions are maintained in the periderm. (A, D) Immunolocalisation of E-cadherin along the apicobasal axis (0 µm is apical) in the periderm of normoxia and hypoxia treated embryos (A) and in the embryos recovered post hypoxia along with normoxic and hypoxic controls (D). (B, E) Graphical representation showing polarised localisation of E-cadherin across normalised cell height in the periderm of normoxic and hypoxic embryos (B) and in embryos recovered post-hypoxia (E). (C,F) A graph showing noise index in E-cadherin localisation in normoxic and hypoxic embryos (C) and in the embryos recovered post hypoxia (F) in the periderm. (G-I) Box-plots showing comparison of cell height (G), apical perimeter (H), and average E-cadherin levels (I) in the normoxic, hypoxic and post-hypoxia recovered embryos in the periderm. Data are median±interquartile range (no of cells assessed for each condition =49-56, N=3). \**P*<0.05, \*\**P*<0.01, \*\*\**P*<0.001, \*\*\*\**P*<0.0001 [Kruskal Wallis with Dunn’s post-hoc test (G) and One-way ANOVA with Tukey’s HSD post-hoc test (H and I). (J) Immunolocalization of ZO-1 in the periderm of normoxic and hypoxic embryos. Data represented in 10 bit-depth. (K) Comparison of the average ZO-1 levels in the normoxic and hypoxic embryos. No of cells assessed for each condition= 57-59, N=3. n.s. not significant *P*>0.05 [Mann–Whitney U test]. Scale bars= 5µm in (A,D) and 10µm in (J).

To test whether these hypoxia-associated phenotypes were reversible, we replaced the hypoxia treated embryos into a normoxic environment for 8 hours. This resulted in restoration of normal squamous morphology characterized by a reduction in cell-height (Fig. 3G) and an increase in apical perimeter (Fig. 3H). In addition, the average E-cadherin levels also recovered in 8 hours (Fig. 3I). However, the polarized distribution of E-cadherin did not display an improvement (Fig. 3D, E, F) within this time window.

Across epithelial systems, exposure to physiological and pathological hypoxia has been reported to result in downregulation of tight junction components resulting in increased permeability and impaired barrier integrity <u>(Bouvry, D., et al 2006, Kimura, K., et al 2010, Luo, H., et al 2024, Huang, J., et al 2025, Springer, M., et al 2025)</u>. Therefore, we assessed the impact of hypoxia on tight junctions using ZO-1, a tight junction scaffolding protein, as a marker. Interestingly, we did not observe any change in ZO-1 levels or localisation under hypoxia as compared to the normoxic control embryos (Fig. 3J, K).

To conclude, exposure to a hypoxic environment during embryonic development results in perturbed epithelial architecture characterized by a reduction in the basement membrane zone and intercellular gaps plausibly due to the decreased cell-cell adhesion and loss of E-cadherin polarity. Several of these cellular characteristics improve upon resumption of oxygen availability, indicating their reversible nature in this developmental window. Notably, the tight junctions remained structurally intact and their localization to the apico-lateral domain did not change as revealed by ZO-1 immunostaining.

### RhoA/ROCK signalling regulates cell morphology and barrier integrity under hypoxia

Experimental evidence from various epithelial systems, supported by theoretical modelling, have shown that cell shapes in epithelia are driven by cell mechanics governed by cell-adhesive and contractile forces <u>(Lee, J.Y. and Goldstein, B., 2003, Dawes-Hoang, R.E., et al 2005, Käfer, J.,</u> <u>et al , 2007, Widmann, T.J. and Dahmann, C., 2009, Hannezo, E. et al, 2014, Tan, R.Z., et al</u> <u>2017, Dasbiswas, K., et al 2018, Herrera-Perez, R.M. and Kasza, K.E., 2018, Luciano, M., et al</u> <u>2022)</u>. We reasoned that the peridermal cell shape changes -from squamous to cuboidal-observed upon exposure to a hypoxic environment may be a consequence of altered force balance between cell-cell adhesive forces and contractility. While the effect on cell-cell adhesion in the basolateral domain was obvious, we asked whether there was any impact on acto-myosin contractility under hypoxia.

To test this, following a 24-hour hypoxia treatment starting at 24hpf, we assessed the levels and localization of phosphorylated myosin light chain (MLC) (Ser19) as a proxy for active NM-II (henceforth referred to as pNM-II) and hence, the acto-myosin contractility. We used ZO-1 as a marker to demarcate the tight junction domain. While pNM-II showed heterogenous localisation in the apical domain with some enrichment at the apico-lateral cortex at the tight junctions under normoxia, this apico-lateral localization was significantly increased upon exposure to hypoxia (Fig. 4A, B).

**Figure 4:**
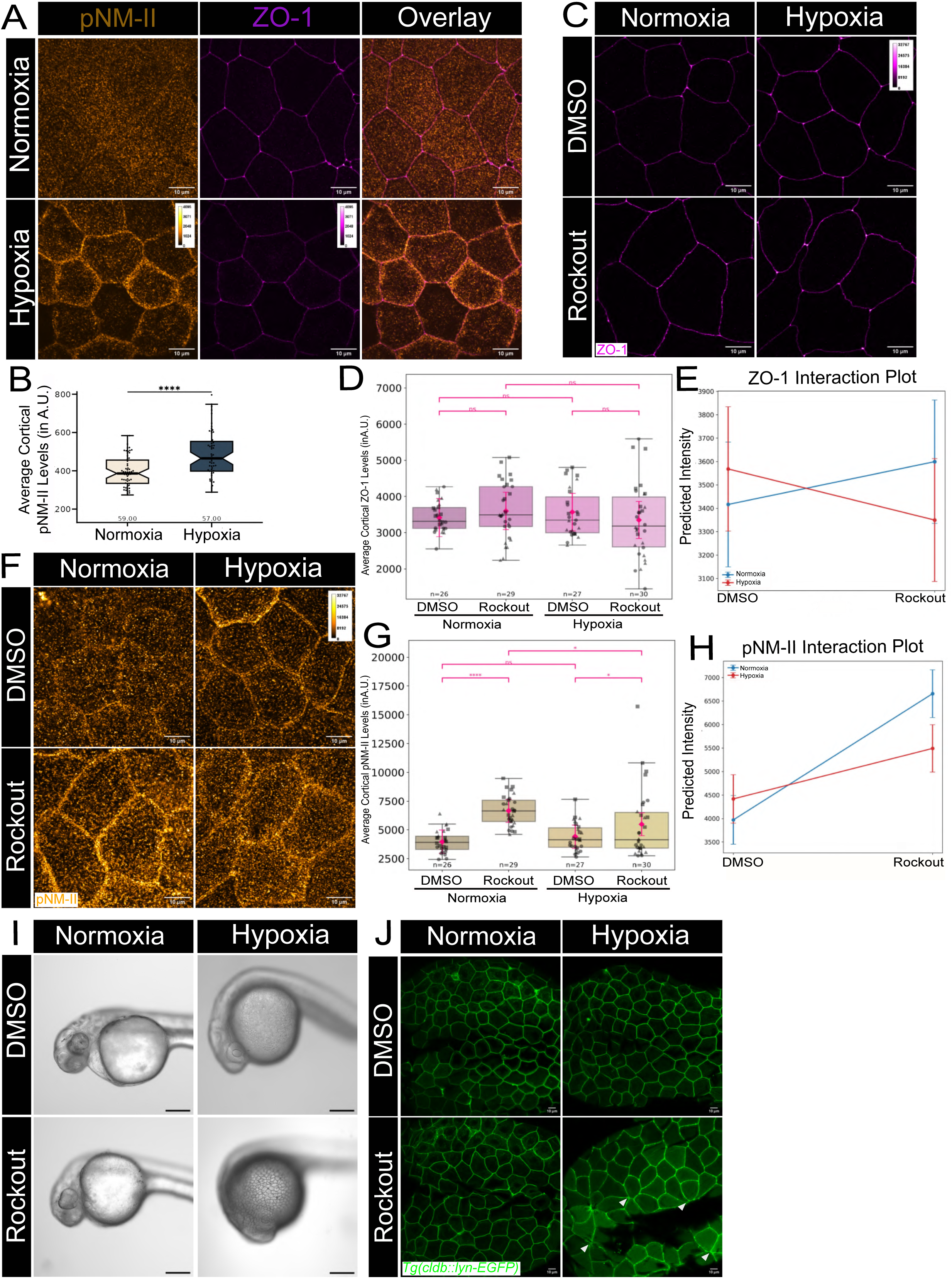
Hypoxia treated embryos display ROCK dependent reorganisation of pNM-II at the tight junctions to ensure barrier integrity. (A, C, F) Representative confocal images showing immunolocalisation of pNM-II and ZO-1 (A), ZO-1 (C) and pNM-II (F) in embryos maintained in embryo media (A) or treated with Rockout and vehicle control DMSO (C,F) under normoxia or hypoxia. (B) Box plot showing comparison of pNM-II levels in the periderm of normoxic and hypoxic embryos. Data are median±interquartile range (no of cells assessed for each condition =57-59, N=3). \*\*\*\**P*<0.0001 [Mann–Whitney U test]. (D,G) Box-plots showing comparison of average ZO-1 levels (D), cortical pNM-II levels (G) in the periderm of DMSO and Rockout treated normoxic and hypoxic embryos. Boxplot represents the median±interquartile range of the raw data (no of embryos assessed for each condition =26-30, N=3). Pink diamonds represent EMMs used for statistical analysis. n.s. not significant *P*>0.05, \**P*<0.05, \*\**P*<0.01, \*\*\*\**P*<0.0001 [Wald z-test]. (●, ▴, ◼ xsrepresent Set1, 2 and 3, respectively). (E,H) Graphs represent ZO-1 (E) and pNM-II (H) interaction plots to depict trends in normoxic and hypoxic embryos treated with DMSO and Rockout. (I) Representative brightfield images of DMSO and Rockout treated normoxic and hypoxic embryos incubated in Evans Blue showing the dye permeance through the epidermis of the hypoxic embryos treated with Rockout (no of embryos assessed for each condition =18, N=3). (J) Representative confocal images of peridermal cells from *Tg(cldnB:lynEGFP)* transgenic line treated with Rockout and vehicle control (DMSO) under normoxia and hypoxia. Note the presence of rosettes (white arrowheads) in Rockout treated hypoxic embryos. Scale bars= 10µm in (A,C,F,J). Data represented in 12-bit depth (A,J) and 15-bit depth (C,F).

RhoA, a central governor of cell shape regulation in epithelia, functions by promoting myosin activity via Rho-associated protein kinase, ROCK <u>(Leung, T., 1995, Fujisawa, K., et al 1996)</u>, a kinase responsible for the phosphorylation of the regulatory light chain of myosin on Ser19 of NM-II (pNM-II), either directly or via the regulation of the activity of MLCK (myosin light chain kinase) <u>(Sellers, J.R., et al 1981, Amano, M., et al 1996)</u>. ROCK also regulates myosin activity by phosphorylating myosin binding subunit (MBS) of MLCP thereby inhibiting myosin phosphatase activity and resulting in increased myosin phosphorylation <u>(Kimura, K., et al 1996)</u>. RhoA and ROCK activity has also been shown to localise at the apical junctional zone surrounding the tight junction <u>(Terry, S.J., et al 2011, Itoh, M. et al, 2012, Zihni, C. and Terry,</u> <u>S.J., 2015)</u>. Therefore, we hypothesized that under hypoxia, the enhanced recruitment of pNM-II to the tight junctions is a consequence of active RhoA/ROCK signalling contributing to reinforcement of tight junctional integrity and the associated cell shape changes.

To test this hypothesis, we treated embryos with Rockout under hypoxic conditions; Rockout is an ATP competitive inhibitor of ROCK activity <u>(Yarrow, J.C. et al, 2005, Sen, S., et al 2026)</u>. We reasoned that in the presence of Rockout, the cells would be unable to acquire the cuboidal morphology under hypoxia if NMII activity is the cause of this cell shape transition. Given the observed lethality associated with a 24-hour Rockout treatment in hypoxic embryos, the impact of ROCK inhibition under hypoxia was assessed after 12-hours of Rockout treatment, followed by evaluation of pNM-II and ZO-1 levels.

While boxplots were used to visualize the raw data, Estimated Marginal Means (EMMs) were computed using a two-way factorial design analyzed via the linear mixed model (LMM) (represented as pink diamonds in the box plots). The EMMs were further used to run pairwise comparisons (see methods for details). We also assessed interactions between independent variables (visualized using interaction plots) in addition to statistical analysis to observe overall treatment trends. The normoxic embryos treated with Rockout displayed an increase in ZO-1 levels (Fig. 4C, D; Supp. Fig. 2A) when compared to the vehicle controls, which was not statistically significant. However, in the subsequent experiments (Fig.5, Fig.6, Fig.7), we observed a statistically robust increase in ZO-1 levels upon Rockout treatment under normoxic conditions, suggesting that the increase we observed here (Fig. 4D) is real. We reason that the statistical insignificance between ZO-1 levels in the data presented (Fig. 4D) is likely due to low statistical power given the small sample size and high variance.

**Figure 5:**
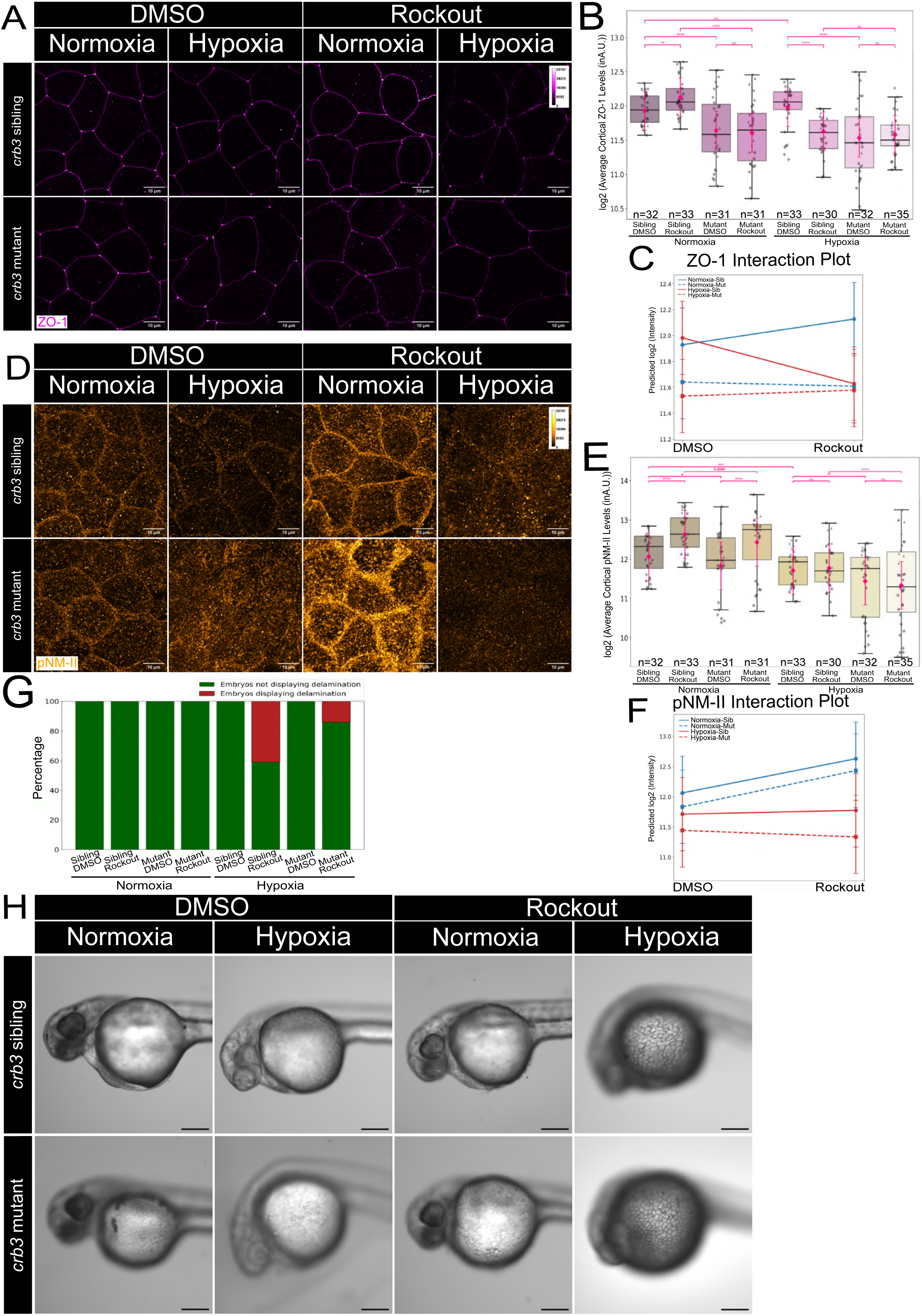
Loss of *crb3a;3b* function prevents peridermal delamination in the Rockout treated hypoxic embryos but not the loss of barrier integrity. (A,D) Representative confocal images showing immunolocalisation of ZO-1 (A) and pNM-II (D) in the periderm of *crb3* sibling and mutant embryos treated with DMSO or Rockout under normoxia and hypoxia for 12 hours. (B,E) Box plots showing comparison of ZO-1 (B) and pNM-II (E) levels in the Periderm of *crb3* embryos treated with DMSO or Rockout under normoxia and hypoxia. Boxplot represents the median±interquartile range of the raw data (no of embryos assessed per condition=30-35, N=4). Pink diamonds represent EMMs used for statistical analysis. n.s. not significant *P*>0.05, \**P*<0.05, \*\*\**P*<0.001, \*\*\*\**P*<0.0001 [Wald z-test]. (●, ▴, ◼, ◆ represent Set 1, 2, 3 and 4 respectively). (C,F) Graphs represent ZO-1 (C) and pNM-II (F) interaction plots to depict trends in *crb3* sibling and mutant embryos treated with DMSO or Rockout under normoxia and hypoxia. (G) Bar graph representing the percentage of embryos displaying rosettes in the *crb3* sibling and mutant embryos treated with DMSO and Rockout under normoxia and hypoxia. (H) Representative brightfield images of DMSO and Rockout treated normoxic and hypoxic *crb3* sibling and mutant embryos incubated with Evans Blue. Note the dye permeance through the epidermis of hypoxic embryos treated with Rockout. Scale bars= 10µm in (A,D) and 200µm in H. Data represented in 15-bit depth (A,D).

**Figure 6:**
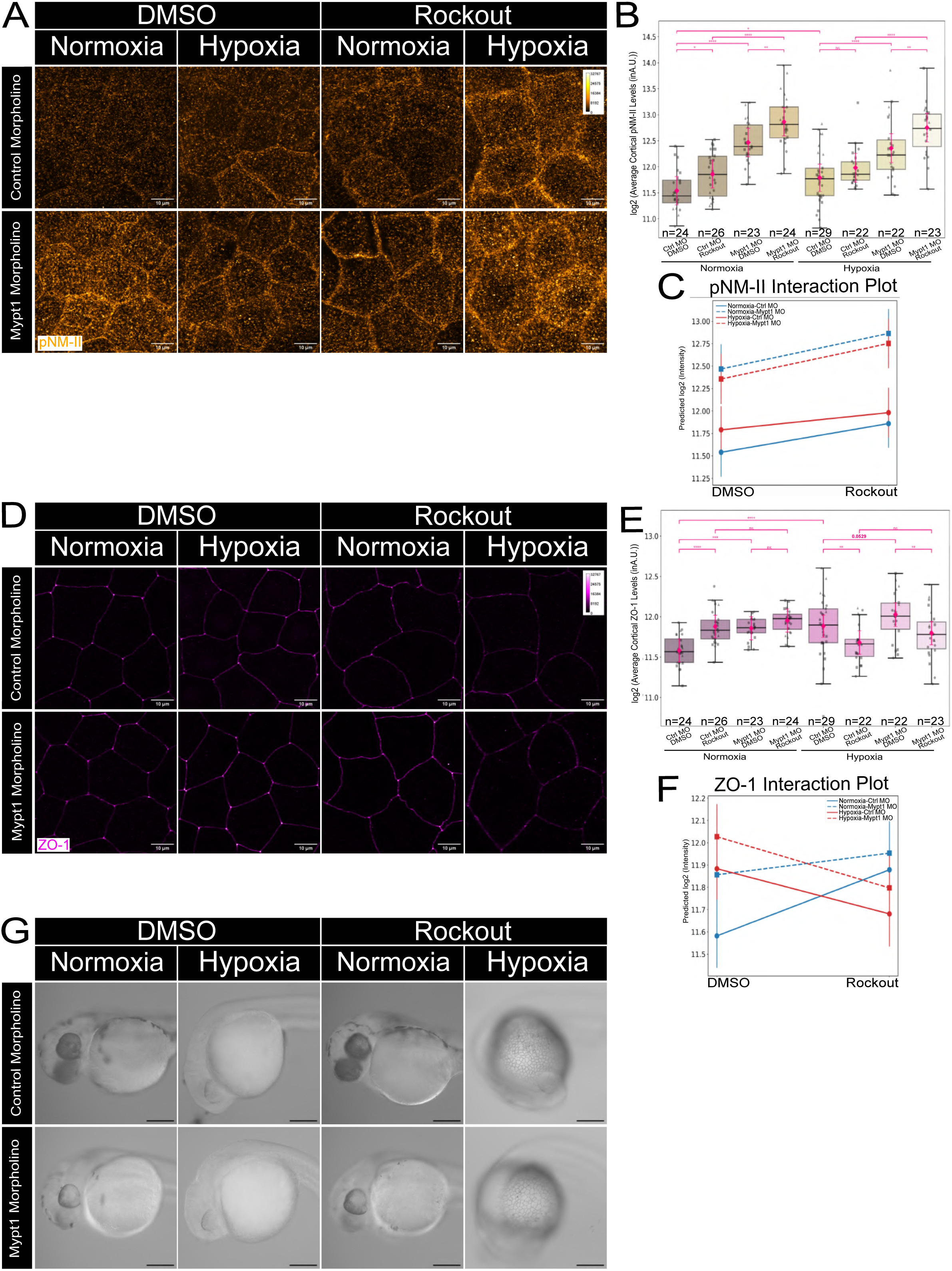
Enhanced Non-muscle myosin-II activity is not sufficient to prevent barrier function loss in hypoxic embryos treated with Rockout. (A,D) Representative confocal images showing immunolocalisation of pNM-II (A) and ZO-1 (D) in the periderm of Control and Mypt1 MO injected embryos treated with DMSO or Rockout under normoxia and hypoxia for 12 hours. (B, E) Box plots showing comparison of pNM-II (B) and ZO-1 (E) levels in the Periderm of Control and Mypt1 MO injected embryos treated with DMSO or Rockout under normoxia and hypoxia. Boxplot represents the median±interquartile range of the raw data (no of embryos assessed per condition=22-29, N=3). Pink diamonds represent EMMs used for statistical analysis. n.s. not significant *P*>0.05, \**P*<0.05, \*\**P*<0.01, \*\*\**P*<0.001, \*\*\*\**P*<0.0001 [Wald z-test]. (●, ▴, ◼ represent Set1, 2 and 3, respectively). (C,F) Graphs represent pNM-II (C) and ZO-1 (F) interaction plots to depict trends in Control and Mypt1 morphant embryos treated with DMSO or Rockout under normoxia and hypoxia. (G) Representative brightfield images of DMSO and Rockout treated normoxic and hypoxic embryos injected with Control and Mypt1 MO and incubated in Evans Blue. Note the dye permeance through the epidermis of hypoxic embryos treated with Rockout. Scale bars= 10µm in (A,D) and 200µm in G. Data represented in 15-bit depth (A,D).

**Figure 7:**
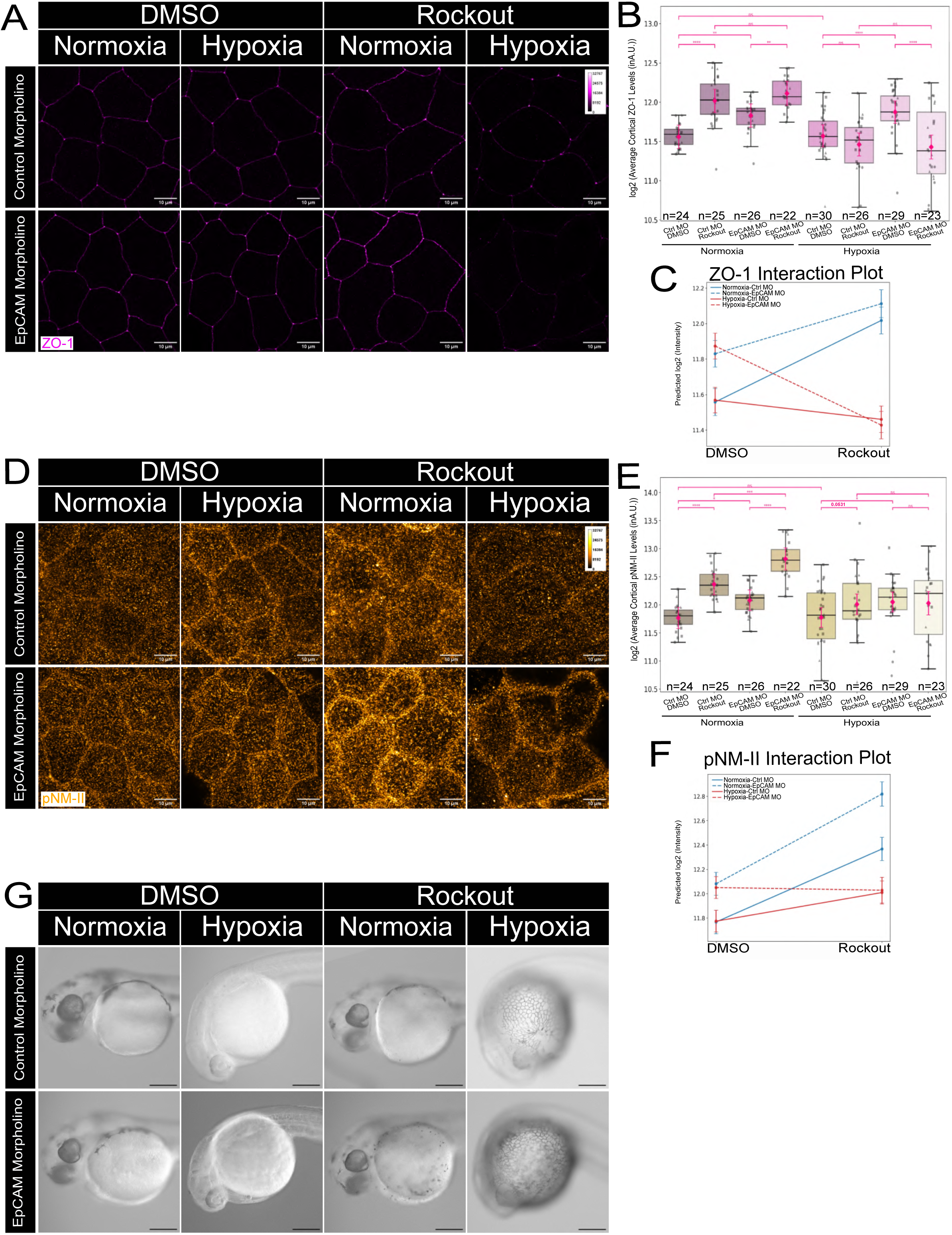
Tight junction augmentation does not prevent barrier defects in the hypoxic embryos treated with Rockout. (A, D) Representative confocal images showing immunolocalisation of ZO-1 (A) and pNM-II (D) in the periderm of Control and EpCAM MO injected embryos treated with DMSO or Rockout under normoxia and hypoxia for 12 hours. (B, E) Box plot showing the comparison of ZO-1 (B) and pNM-II (E) levels in the Periderm of Control and EpCAM MO injected embryos treated with DMSO or Rockout under normoxia and hypoxia. Boxplot represents the median±interquartile range of the raw data (no of embryos assessed per condition=22-29, N=3). Pink diamonds represent EMMs used for statistical analysis. \*\**P*<0.01, \*\*\**P*<0.001, \*\*\*\**P*<0.0001, n.s. not significant *P*>0.05 [Wald z-test]. ( ●, ▴, ◼ represent Set1, 2 and 3, respectively). (C,F) Graphs represent ZO-1 (C) and pNM-II (F) interaction plots to depict trends in Control and EpCAM morphant embryos treated with DMSO or Rockout under normoxia and hypoxia. (G) Representative brightfield images of DMSO and Rockout treated normoxic and hypoxic embryos injected with Control and EpCAM MO and incubated in Evans Blue. Note the dye permeance through the epidermis of hypoxic embryos treated with Rockout. Scale bars= 10µm in (A,D) and 200µm in G. Data represented in 15-bit depth (A,D).

Rather unexpectedly, pNM-II showed a significant increase in the apico-cortical localisation upon Rockout treatment under normoxic conditions (Fig. 4F, G; Supp. Fig. 2B). Despite the additional developmental delay seen in Rockout treated hypoxic embryos, the pNM-II localisation persisted at the apicolateral domain in Rockout treated hypoxic embryos in comparison to the vehicle control albeit displaying a more diffuse localisation at the apical cortex (Fig. 4F). Besides, although the increasing trend was observed (Fig. 4H), 12 hours of hypoxia did not result in statistically significant increase in pNMII levels (Fig. 4G) unlike that seen in embryos exposed to 24 hours of hypoxia (Fig. 4B). Notably, the levels of pNM-II in Rockout treated hypoxic embryos were significantly lower as compared to Rockout treated normoxic embryos (Fig. 4G), indicating that the compensatory mechanisms that act under normoxic conditions to enhance the pNM-II levels upon Rockout treatment do not operate under hypoxic conditions. The hypoxic embryos treated with Rockout did not display apically constricted peridermal cells as compared to the control (Supp. Fig. 2C, D) suggesting that the activation of RhoA-ROCK signaling is required to modulate the cell morphology under hypoxia.

While ZO-1 localisation did not show statistically significant change in the hypoxic embryos treated with Rockout (Fig. 4C, D), possibly due to low statistical power, the interaction plot (Fig. 4E) revealed a trend towards reduced ZO-1 levels in the hypoxic embryos treated with Rockout as compared to the hypoxic vehicle control. In subsequent experiments, presented later, we also observed robust decrease in ZO-1 localisation in Rockout treated hypoxic embryos suggesting that the effect of Rockout on ZO-1 is real. This along with the diffused cortical localisation of pNM-II in Rockout treated hypoxic peridermal cells mentioned above (Fig. 4F) raised the possibility of a functional consequence to the barrier integrity. Therefore, we asked whether ROCK signalling would be important for the maintenance of TJ-mediated barrier function. To test this possibility, we performed a permeability assay at the end of the 12-hour treatment of Rockout with hypoxia using Evans Blue Dye (EBD), a 960Da tracer molecule. Evans blue has been used as an indicator of leaky tight junctions in the epithelia lining the blood-brain barrier <u>(Waithe, O.L.Y., et al 2023, Nidavani, R.B. et al, 2014, Radu, M. and Chernoff, J., 2013)</u>. Similarly, in the epidermis, TJs constitute nearly “impermeable barriers” as mentioned before <u>(Citi, S., et al 2024)</u>. Upon incubation with this dye for 60 minutes at the end of the 12-hour treatment, we observed that 100% of the hypoxic embryos (n=6, N=3) treated with Rockout exhibit an increase in the permeance of the dye whereas all other controls displayed an intact barrier (Fig. 4I).

To conclude, under hypoxia, there is enhanced apical localisation of pNM-II, which results in a shift from squamous to cuboidal morphology of the peridermal cells. ROCK signaling is required for the maintenance of barrier function and the restricted cortical localisation of pNMII at the apico-lateral cortex under hypoxia. Under normoxic conditions, Rockout treatment results in an increase in ZO-1 and pNM-II localisation, possibly due to the presence of compensatory mechanisms (Discussed later in detail).

### Crb3 promotes cell-delaminations and functions redundantly with ROCK to regulate pNMII levels under hypoxia

In addition to a partial rescue in the cell morphology phenotype and appearance of permeability defects, we also observed the formation of cell-rosettes (Fig. 4J) in the periderm of approximately 50-60% of hypoxic embryos treated with Rockout. The rosettes were accompanied by delaminating cells labelled with lyn-EGFP indicating their peridermal origin. Furthermore, these delaminating cells showed nuclear fragmentation and were positive for Activated Caspase-3 (Supp. Fig. 2E). The presence of delaminating cells was surprising since the Rockout treatment was expected to reduce the actomyosin contractility required for apical constriction. We reasoned that in a few peridermal cells, the residual apical recruitment of pNM-II results in higher effective contractile force directed inwards, possibly due to a substantial reduction in countering cell adhesive force at the tight junction and local tension heterogeneities. This generates a force imbalance leading to apical constriction and stochastic delaminations seen in Rockout treated hypoxic embryos. To test this hypothesis, we used *crb3* mutant strain which shows decreased pNMII levels in the peridermal cells at 48hpf under normoxia (Supp. Fig. 3D, E; Bose et al, *manuscript in preparation*). The *crb3b* paralog is abundantly expressed in the periderm during early development <u>(Sur, A., et al 2023)</u>. However, to eliminate the possibility of genetic compensation by the *crb3a* paralog and maternal contribution, we used the *crb3a;crb3b* -maternal zygotic mutants in this study. Statistical analysis was performed on computed EMMs as before. We observed that *crb3a;crb3b m/z* (henceforth referred as *crb3*) mutant embryos at 48hpf under normoxia and hypoxia (24 hour treatment), displayed a significant reduction in the ZO-1 levels (Supp. Fig. 3A, B, C, G), and pNM-II levels (Supp. Fig. 3D, E, F, H) as compared to their respective siblings. These results indicate that Crumbs function is essential for the localisation of ZO-1 and pNM-II during development and hypoxia.

We further conducted the hypoxia treatment in the *crb3* background and combined it with Rockout treatment. Since hypoxia combined with Rockout treatment for 24-hours is lethal, the exposure to hypoxia and Rockout was done for 12-hours and hence, the embryos were fixed at 36hpf and analysed for ZO-1 and pNMII localisation. Under these conditions, the effect of *crb3* loss on ZO-1 and pNM-II localisation was evident under both normoxia and hypoxia (Fig 5A, B, D, E). Following Rockout treatment the normoxic *crb3* sibling embryos treated with Rockout showed a robust increase in ZO-1 and pNM-II localisation in comparison to their vehicle controls. However, while the normoxic *crb3* mutant embryos showed a significant increase in pNM-II localisation following a Rockout treatment as compared to its vehicle control, a similar robust increase was not observed in ZO-1 levels. Besides, under hypoxia, neither Rockout treated *crb3* siblings nor mutant embryos showed increase in the localisation of ZO-1 and pNM-II relative to their respective vehicle controls. (Fig. 5A, B, D, E; Supp. 3I, J). Importantly, under hypoxia, Rockout treated mutant embryos showed significant reduction in pNM-II levels as compared to Rockout treated sibling embryos. Consistently, we observed a reduction in the proportion of embryos that displayed rosettes and therefore, delamination events (Fig. 5G).

We further asked whether the permeability defect is a direct consequence of RhoA-ROCK signaling regulating tight junctions, or an indirect consequence of cell delaminations. To investigate this aspect, we treated the *crb3* mutant embryos with Rockout under hypoxia for 12 hours and assessed the effect on the barrier function. Interestingly, despite a decrease in the proportion of embryos showing rosettes under ROCK inhibition in *crb3* mutant hypoxic embryos (Fig. 5G), the permeability defect persisted (Fig. 5H), suggesting a direct regulation of tight junction permeability by RhoA-ROCK signaling under hypoxia.

To conclude, ROCK mediated high pNM-II activity is essential to prevent cell delaminations and apoptosis under hypoxia. Upon ROCK inhibition, Crb3 contributes to residual pNM-II recruitment, which results in delamination and apoptosis of peridermal cells likely due to imbalance between the inward actomyosin contractile force and the cell adhesive force.

### ROCK signalling regulates tight junction remodelling to maintain barrier function under hypoxia

A RhoA/ROCK mediated programme is known to operate during morphogenesis or wound healing responses to repair and reinstate the barrier function. It exerts these effects on barrier integrity primarily by two ways, via regulating the actomyosin cytoskeleton and by controlling the availability of components such as Claudins for tight junction formation <u>(Amano, M., et al</u> <u>1996. *Journal of Biological Chemistry*, Kawano, Y., et al 1999. *The Journal of cell biology*, Yu,</u> <u>D., et al 2010. *PNAS*, Terry, S.J., et al 2011. *Nature cell biology*, Itoh, M. et al, 2012. *PNAS*, Arnold, T.R., et al 2017, Stephenson, R.E., et al 2019, Higashi, T., et al 2022, Craig, Z., et al</u> <u>2025, Cho, Y., et al 2025)</u>. Therefore, we asked whether ROCK signalling controls the barrier function in the developing zebrafish epidermis by regulating tight junction maintenance or actomyosin contractility.

To test whether ROCK signaling operates via regulating actomyosin contractility in hypoxic embryos to maintain the barrier function, we asked whether increasing NM-II activity in the absence of RhoA-ROCK signaling would be sufficient to maintain the barrier function. To achieve this, we used the previously published myosin phosphatase-targeting subunit 1 (*mypt1*) morpholino <u>(Huang, H., et al 2008)</u>. We reasoned that the loss of *mypt1* function would result in higher pNM-II levels, due to dampened MLC dephosphorylation during the window of ROCK inhibition, allowing us to test the importance of pNM-II in barrier function maintenance under hypoxia. We observed that the Mypt1 morphants displayed significantly higher levels of pNM-II and ZO-1 (Fig. 6A, B, D, E; Supp. Fig. 4C, D) under vehicle-treated normoxic and hypoxic conditions as compared to the control morphant embryos. Notably, in this condition, there was also a significant increase in ZO-1 localisation (Fig. 6D, E, Supp. Fig. 4C), suggesting possible augmentation of tight junctions due to improved cytoskeletal support. Importantly, normoxic Rockout treated Mypt1 morphants showed pNMII levels over and above the Rockout treated control morphants. However, this increase in pNMII did not translate into an increase in ZO-1 localisation in Rockout treated Mypt1 morphants, indicating that mere gain in pNMII level may be insufficient to augment tight junctions when ROCK signaling is inhibited. Further, under hypoxia, Rockout treated Mypt1 morphants showed an increase in pNM-II (Fig. 6 A, B, C; Supp. Fig. 4D) as compared to both the Rockout-treated control and the vehicle-treated Mypt1 hypoxic morphants. Interestingly, unlike their vehicle-treated counterparts, the hypoxic Mypt1 morphants treated with Rockout were unable to display an increase in ZO-1 and these levels were not significantly different from the Rockout treated hypoxic control morphants (Fig. 6 D, E, F; Supp. Fig. 4C), again driving the point that Mypt1 loss is sufficient for pNMII gain in the ROCK inhibited embryos under hypoxia but without promoting tight junction augmentation. Similar to the previous experiments (Fig. 4F), pNM-II showed diffused localisation at the apico-lateral cortex despite the increase in levels in Rockout-treated Mypt1 morphants under hypoxia. Consistent with this localisation defect in pNM-II and significant decrease in ZO-1 levels, the Rockout treated Mypt1 and control morphant embryos showed the barrier function defects under hypoxia (Fig. 6G).

We reasoned that diffused localisation of pNM-II in Rockout treated hypoxic Mypt1 and control morphants is a consequence of dysregulated tight junctions and reduced ZO-1 localisation, suggesting that RhoA-ROCK signaling primarily controls tight junction formation under hypoxia. Indeed, RhoA-ROCK signalling is known to promote the cleavage of EpCAM by activating Matriptase, thereby promoting the release of Claudin sequestered with EpCAM complexes <u>(Higashi, T., et al 2022, Cho, Y., et al 2025)</u> and increasing its availability for *de novo* tight junction formation. Interestingly, however, it was shown in the enveloping layer (EVL) of zebrafish embryos (precursor of the outer epidermal layer) that the loss of EpCAM function results in increased levels of ZO-1 <u>(Slanchev, K., et al 2009)</u>. This increase in ZO-1 localisation in *epcam* morphant embryos is likely to be a consequence of the increased availability of Claudin pool for the formation of tight junctions. We reasoned that if tight junction remodelling is indeed controlled by RhoA-ROCK signalling, this augmentation in ZO-1 localisation in *epcam* deficient embryos should get rescued or reduced in Rockout treated hypoxic embryos. To test this, we used the previously published morpholino to knockdown EpCAM function <u>(Slanchev, K., et al</u> <u>2009)</u>. We observed that, under normoxic and hypoxic conditions, the vehicle-treated EpCAM morphants displayed significantly higher levels of ZO-1 (Fig. 7 A, B; Supp. Fig. 4A) as well as pNM-II (Fig. 7 D, E, Supp. Fig. 4B) as compared to the control morphant embryos indicating augmentation in tight junction formation. Rockout treatment in both the EpCAM and control morphants under normoxia resulted in an increase in ZO-1 and pNM-II localisation relative to their respective vehicle controls (Fig. 7B, E). Furthermore, EpCAM morphants treated with Rockout under normoxia did not show an increase in ZO-1 levels when compared with their Rockout treated control morphant counterparts, suggesting that under normoxia ROCK independent mechanisms could operate in developing embryos (Fig. 7B, C). Importantly, under hypoxia, Rockout treated EpCAM morphants showed significant reduction in ZO-1 levels as compared to the vehicle-treated EpCAM morphants (Fig. 7B,C; Supp. Fig. 3A). In contrast to normoxic embryos and despite the heterogeneity in levels, the Rockout-treated hypoxic EpCAM morphants did not show an increase in pNM-II as compared to both the Rockout-treated control and vehicle-treated EpCAM morphants (Fig. 7 E, F). Consistent with their inability to improve either pNM-II or ZO-1 levels or localisation, the Rockout treated hypoxic EpCAM morphants displayed barrier function defects like its control morphant counterparts (Fig. 7G).

Taken together, while EpCAM loss promotes increased ZO-1 recruitment indicative of improved tight junctions, ROCK inhibition, specifically under hypoxia, reduces ZO-1 localisation and increased permeance. Notably, the loss of ZO-1 recruitment upon ROCK inhibition persisted even when bulk myosin activity was restored via Mypt1 knockdown, indicating that ROCK’s role in junctional remodelling under hypoxia cannot be explained by increased myosin activity alone. Our data thus suggests that ROCK regulates tight junction integrity under hypoxic conditions through a mechanism that extends beyond generalized cytoskeletal regulation and primarily involves tight junction remodelling. However, we cannot rule out the possibility of ROCK’s involvement in localized cytoskeletal organisation at the junction itself.

## Discussion

How epithelial tissues -which contribute to respiration, barrier function and animal survival-cope with hypoxic environments has remained poorly understood. The developing epidermis, similar to the lung and gill epithelia, participates in gas exchange in addition to acting as a barrier during development. In this study, we uncover the ROCK mediated protective mechanism that maintains the barrier function specifically under hypoxic conditions in a developing epidermis.

Tight junctions are the primary regulators of paracellular transport and therefore, barrier function in epithelial tissues. The homeostatic regulation of tight junctions under physiological conditions has primarily been attributed to the activity of RhoA-ROCK signalling which eventually impinges upon cytoskeletal regulation by controlling NM-II activity and actin organisation <u>(Mack, N.A. and Georgiou, M., 2014, Terry, S.J., et al 2011, Itoh, M., et al 2012</u>; <u>Amano, M., et</u> <u>al 1996, Kawano, Y., et al 1999, Yu, D., et al 2010, Acharya, B.R., et al 2018, Stephenson, R.E.,</u> <u>et al 2019, Guan, G., et al 2023)</u>. More recently, it has been shown that RhoA-ROCK signaling regulates tight junctions via the activation of membrane anchored serine protease Matriptase, resulting in the cleavage and release of Claudins sequestered in Claudin-EpCAM/TROP2 complexes <u>(Higashi, T., et al 2022, Cho, Y., et al 2025)</u>. While the paracellular transport via tight junctions is largely regulated by the transmembrane barrier proteins, the tension exerted by the underlying cytoskeleton can result in altered barrier properties, highlighting the importance of scaffolding protein, ZO-1. ZO-1 links the transmembrane tight junction complex with the circumferential actomyosin <u>(Fanning, A.S., et al 1998, Rouaud, F., et al 2023)</u>, and contributes to the maintenance of barrier function <u>(Umeda, K., et al 2006, Van Itallie, C.M., et al 2009, Citi, S.,</u> <u>2019)</u>. This crosstalk between the core tight junction components and the associated actomyosin cytoskeleton is essential in maintaining barrier properties <u>(Madara, J.L., et al 1987, Turner, J.R.,</u> <u>et al 1997, Ivanov, A.I., et al 2004, Steed, E., et al 2010, Spadaro, D., et al 2017)</u>. In contrast to these published studies, our results show that during development under normoxic conditions, ROCK signaling is dispensable for tight junction maintenance and barrier function. In fact, ROCK inhibition results in augmentation in both pNMII as well as ZO-1 levels under normoxia. It is likely that in a developing epithelial system such as the epidermis, additional regulators of myosin phosphorylation such as MLCK, ZIPK, citron kinase could participate in its recruitment and subsequent activity at the tight junctions, thereby ensuring barrier maintenance <u>(Ikebe, M.</u> <u>and Hartshorne, D.J., 1985, Murata-Hori, M., et al 1999, Yamashiro, S., et al 2003)</u>.

Further, we show that under hypoxic conditions, active NM-II is enriched at the tight junctions in the developing epidermis. While E-cadherin polarity is lost and cell-cell adhesion is compromised, the tight junction integrity and the barrier function remain intact. We further demonstrate that ROCK signalling is essential for the tight apico-cortical localisation of pNMII and maintenance of ZO-1 levels. Importantly, it functions as a major regulator of the tight junction mediated barrier function under hypoxic environments. Our data reveals that upon ROCK inhibition, increased pNMII levels in *mypt1* deficient embryos display diffused cortical organisation in the hypoxia-treated embryos. Besides, these increased pNM-II levels in *mypt1* morphants do not translate into increased ZO-1 localisation, a proxy for tight junctions. Furthermore, despite augmented formation of tight junctions in EpCAM deficient embryos, the ROCK inhibition results in the loss of tight junctions and barrier function defects under hypoxia. These data indicate that ROCK signalling primarily regulates the tight junction formation under hypoxia (Figure 8). Given the importance of tight junctions in organising apico-lateral actomyosin, it is conceivable that loss of tight junctions in hypoxic embryos upon ROCK inhibition results in diffused apico-cortical localisation of pNM-II in all the genetic conditions that we analysed. The RhoA-ROCK-Matriptase-EpCAM axis is shown to regulate tight junction formation in cultured simple and stratified epithelial systems <u>(Higashi, T., et al 2022, Cho, Y., et</u> <u>al 2025)</u>. However, it is not clear how EpCAM functions in tight junction formation in the developing zebrafish epidermis. Given that ROCK inhibition in hypoxic EpCAM deficient embryos results in a stark decrease in tight junction formation and barrier function defects, it is unlikely that under hypoxia RhoA-ROCK functions alone in tight junction remodelling via matriptase mediated EpCAM cleavage and release of Claudins. Further investigation will be required to understand the mechanism of tight junction remodelling downstream of ROCK signaling under hypoxia.

**Figure 8:**
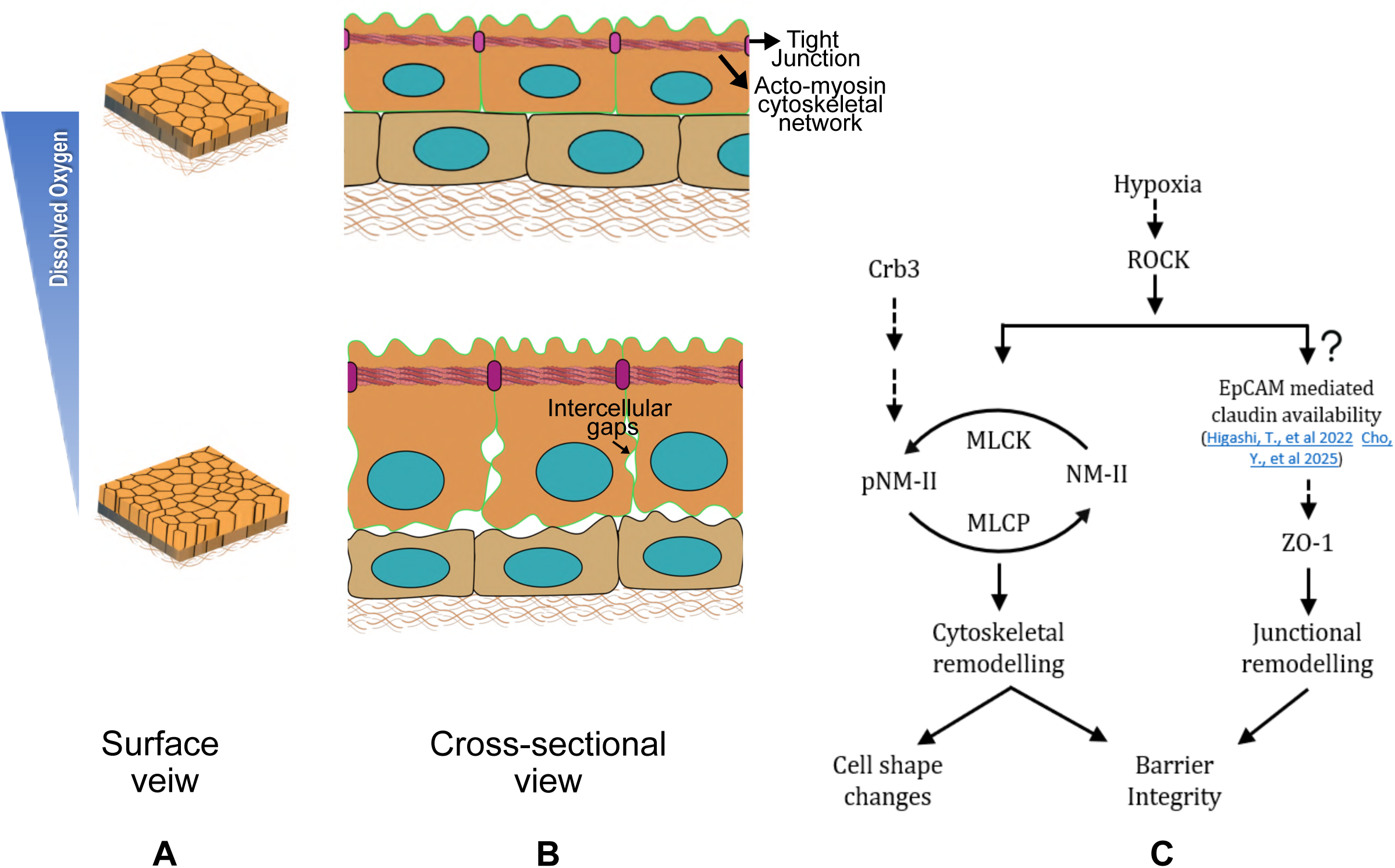
Schematic representation of the response of a developing epithelium to a hypoxic stress and its underlying molecular regulation. The squamous morphology of the periderm cells in normoxic embryos changes to cuboidal morphology upon hypoxic exposure (Surface view) (A) accompanied by the appearance of intercellular gaps (Cross-section) (B). ROCK regulates both junctional and cytoskeletal remodelling to ensure barrier integrity under hypoxia, while Crumbs participates in cytoskeletal regulation and modulates cell morphology. Inhibition of ROCK results in perturbed barrier integrity, evidenced by reduced ZO-1 and pNM-II and the resultant force imbalance due to residual pNM-II at the tight junction results in peridermal cell delamination (C; see text for details).

The ROCK mediated phosphorylation of NM-II, either directly or via MLCK <u>(Sellers, J.R., et al</u> <u>1981, Amano, M., et al 1996)</u>, and its increased tight localisation at the tight junction combinatorially fortify the tight junctions under the hypoxic condition. Our data indicate that one of the consequences of an increased apical localisation of the active NM-II under hypoxia is the change in cell morphology from squamous to cuboidal (Fig. 8). While assessing the role of ROCK mediated NM-II activity in this morphology change, we observed delaminations and apoptosis of a few peridermal cells, since both tight and adherens junctions are compromised. We reason that these delaminations are a consequence of altered balance between cell adhesive and contractile forces at the apical cortex. Under hypoxia, the strong contractile force operational in the apico-lateral cortex due to pNM-II enrichment, is likely balanced by the tight junction reinforcement (due to ROCK activity) in the peridermal cells. Perturbed junctional remodelling during ROCK inhibition under hypoxia results in an imbalanced NM-II activity at the apical cortex leading to apical constriction, followed by delamination and apoptosis. This may further be aided by the local heterogeneities in tissue tension, resulting in stochasticity seen in delaminations. Indeed, in *crb3* mutant embryos, the dampening of NM-II localisation substantially reduced the frequency of peridermal delaminations. Crumbs, an apical polarity determinant, is known to interact with FERM domain containing proteins like 4.1/Ezrin/Radixin/Moesin by its cytoplasmic C-terminal domain <u>(Pocha, S.M. and Knust, E.,</u> <u>2013)</u> and to regulate myosin activity primarily through a Rho/ROCK/pNM-II mediated regulation either directly or indirectly <u>(Röper, K., 2012, Flores-Benitez, D. and Knust, E., 2015, Silver, J.T., et al 2019)</u>. The hypoxia and Rockout analysis in *crb3* mutant is consistent with barrier defects persisting even when delamination frequency is reduced and highlights the importance of sustained junctions to ensure barrier integrity.

Our findings show that under hypoxia, the developing epidermis displays a loss of E-cadherin polarity along with a reduction in its levels in the periderm (Fig. 3B) plausibly resulting in the intercellular gaps in the epidermis. Both, simple and stratified epithelial systems highlight the importance of E-cadherin mediated adhesion in regulating the formation and maintenance of tight junctions and vice versa, which are essential for the mechano-response and barrier function of epithelia <u>(Gumbiner, B., et al 1988, Tunggal, J.A., et al 2005, Ikenouchi, J., et al 2007, Ooshio, T., et al 2010, Maiers, J.L., et al 2013, Rübsam, M., et al 2017, Shigetomi, K., et al</u> <u>2018)</u>. Interestingly, our findings show that in the bilayered epithelium of the developing zebrafish epidermis, while E-cadherin polarity is perturbed and its levels are reduced upon exposure to hypoxia, this does not impact the levels or localisation of tight junctions, indicating that E-cadherin is dispensable in regulating tight junction maintenance. Further, given that overactivation of ROCK has been associated with the disruption of adherens junctions <u>(Sahai, E.</u> <u>and Marshall, C.J., 2002)</u>, our work raises an interesting possibility of ROCK signaling regulating both, the tight junctions to maintain the barrier function and disassembly of adherens junctions to reduce the energy requirement of their maintenance. In the energy constrained cellular environment under hypoxia, this reinforcement of tight junctions, possibly at the expense of adherens junction maintenance, promotes the survival of the organism. This further provides organisms with the opportunity to recover when normoxic conditions are restored. Indeed, our data clearly show that most of the hypoxic phenotypes recover within 8 hours, upon resumption of normoxic conditions, except the E-cadherin polarity, which possibly requires recovery in synthesis and trafficking, which likely operates on different timescales.

Physiologically, hypoxia is routinely experienced by epithelia. A conserved response across different epithelia involves modulation of the expression of junctional proteins and regulation of cytoskeleton in a Hif-dependent manner highlighting the importance of maintaining barrier integrity <u>(Jain, M. and Sznajder, J.I., 2005, Zhou, G., et al 2009, Rezvani, H.R., et al 2011, Saeedi, B.J., et al 2015, Manresa, M.C. and Taylor, C.T., 2017, Muenchau, S., et al 2019, Zieseniss, A., 2014, Flood, D. and Taylor, C.T., 2025)</u>. Furthermore, most of the hypoxia mediated cellular responses are largely studied under pathological conditions and using cell culture models. Our study establishes an *in vivo* epithelial model to understand the mechanisms underlying the tissue response under physiologically relevant hypoxic conditions. We establish RhoA-ROCK signalling as a major driver of the epithelial response to developmental hypoxia, where it elegantly rewires the junctional and cytoskeletal machinery to ensure survival of the organism.

## Materials and methods

### Ethics Statement

Zebrafish husbandry and experimental handling were conducted with the guidelines recommended by the Committee for the purpose of Control and Supervision of Experiments on Animals (CPCSEA), Govt. of India, and approved by the institutional Animal Ethics Committee (TIFR/IAEC/2019-6 and TIFR/IAEC/2024-5).

### Fish strains

Experiments were done in Tübingen (Tü) wild-type strain unless mentioned otherwise. Fish were housed at 28°C on a 13h:11h light:dark cycle. The transgenic fish, *Tg(cldb:lyn-EGFP)* <u>(Haas and</u> <u>Gilmour, 2006)</u> were used for experiments that involved visualization of the plasma membrane. The *crb3a;3b m/z* mutant lines used in this study were generated by Satu Kujawski and Elisabeth Knust at the Max-Planck Institute for Molecular Cell Biology and Genetics and will be published elsewhere (Bose et al, *Manuscript under preparation*).

### Physical hypoxia chamber for zebrafish embryos

A hypoxic microenvironment was created by the controlled infusion of Nitrogen in the embryo medium (E3) placed in T75 Nunc™ EasYFlask™ Cell Culture Flasks (Thermo Scientific; Catalog No: 156472) until a dissolved oxygen concentration of 2-3 mg/L was attained, which was measured with the help of a DO meter (Vernier Optical DO Probe; Catalog No: ODO-BTA) and subsequently monitored for 24 hours. All embryos were incubated at 28.5° C and staged according to <u>Kimmel et. al.1995</u> and the treatment was initiated at 24 hours post fertilization (hpf) for a duration of 24 hours except for experiments involving a treatment with Rockout, which were for a duration of 12 hours.

### Microinjections

EpCAM morpholino (5′-GTGCAGAGACTTTCCGGCCATATTT-3′) and its 5bp mismatch control (5’-GTCCAGACAGTTTCCCGCGATATTT-3) (1.5nl of 200µM) <u>(Slanchev, K., et al</u> <u>2009)</u>, and Mypt1 morpholino (mypt1-MO, 5′-CGTAACGCAACGCTCTTCTTACCTG-3′) <u>(Huang, H., et al 2008)</u> along with the standard control morpholino (5’-CCTCTTACCTCAGTTACAATTTATA-3’) (1nl of 500µM) were microinjected in embryos at 1-2 cell stage using the World Precision Instruments PV830 Pneumatic PicoPump. All injections in a single set were performed in a single session. All morpholinos were procured from Gene Tools, LLC (Eugene, Oregon, USA).

### Rockout treatment

Rockout was prepared in DMSO and the final DMSO concentration during each experiment was 1% in both the test and control conditions. The treatments were administered in embryo medium (E3) without methylene blue. For every condition, 15-20 embryos at 24hpf were taken in either a 6-well plate (3ml; Tarsons 6-Well Cell Culture Plate; Catalog No. 980010) for normoxia or a T25 cell culture flask (10ml; Falcon® 25cm² Rectangular Canted Neck Cell Culture Flask with Blue Plug Seal Screw Cap; Catalog No: 353014) to maintain hypoxia. Embryos were treated with Rockout (100µM; SCBT, Catalog No: sc-203237; CAS No:7272-84-6) along with hypoxia for 12 hours. The treatment was terminated with three washes of E3 buffer without methylene blue for 5 min each before fixation in 4% PFA in PBS for 30 min followed by overnight at 4°C.

### Evans Blue Permeability Assay

Briefly, embryos were treated with Rockout for 12 hours followed by three washes in E3 without methylene blue and incubated in a 0.02% Evans Blue solution (Stock=2% w/v) (Catalog No: Sigma, E2129) for 60 mins. Due to the fragility exhibited by the epidermis post-dye incubation, embryos were imaged directly in the dye containing media using Olympus MVX10 microscope. Only the Green channel images were presented in figures.

### Electron Microscopy (TEM)

For TEM, embryos were fixed in 2.5% glutaraldehyde (Sigma, Catalog No: G6257) in 0.1 M Phosphate buffered saline (PBS) at pH 7.2 for 30 minutes at RT followed by 4°C overnight. This was followed by PBS washes, 3 times for 15 minutes each then stained in 0.2% Uranyl acetate (EMS; Catalog No: 22400) for 2 hours at 4°C followed by 1% OsO_4_ (Sigma, Catalog No: 60H0150) at 4°C on a rotor overnight. The embryos were then washed 6 times (each wash 10 mins) with autoclaved distilled water and processed for dehydration in acetone series prepared in autoclaved distilled water (30%, 50%, 70%, 90%, 100%, 100%, 100%) for 5 min each. Embryos were equilibrated in acetone: Durcupan (2:1), acetone: Durcupan (1:1) and Durcupan (EMS, Catalog No: 14040) for 60 mins each at RT and then in fresh Durcupan overnight. This was accompanied with Durcupan coating in several moulds which were cured at 60°C overnight, followed by embryo placement in the moulds with Durcupan and further curing at 60°C overnight. For standard EM, 70–100 nm sections cut with a diamond knife (DiATOME) were collected on formvar coated slot or mesh grids or uncoated mesh grids and post stained with uranyl acetate and lead citrate and imaged on a JEM 1400 PLUS, JEOL Transmission Electron microscope.

### Immunostainings

Embryos were fixed in 4% PFA in PBS for 30 mins at RT followed by overnight in 4°C. The subsequent day, fixative was washed off with two 5-minute PBS washes followed by permeabilization in 0.8% Triton X-100 in PBS (PBT) for five 10-minute washes. 10% Normal Goat Serum (Jackson Immuno Research Labs; Catalog No: 005-000-121) in PBT, was used for blocking at room temperature (RT) for 3-4 hours, followed by primary antibody incubation in PBT for 12-16 hours (20hr for pMLC), at 4°C on a rotor. Embryos were then washed with five 30-minute PBT washes at RT on the rocker followed by incubation with secondary antibodies or fluorescent conjugates made in PBT for 3-4 hours at RT on the rotor. This was followed by seven 10-minute washes with PBT and post-fixation with 4% PFA in PBS for 30-minutes at RT. The fixative was washed off with two 5-minute washes in PBS followed by serial upgrade in glycerol (30%, 50%, 70%) and kept in 4°C until imaged.

The following primary antibodies and dilutions were used in this study: pMLC(Ser19) (1:50; Cell Signaling Technologies; Catalog No: 3671), chicken anti-GFP (1:250; Abcam; Catalog No: ab13970), mouse anti-E cadherin (1:200; BD Biosciences; Catalog No: 610182), rabbit anti-active Caspase 3 (1:200; BD Biosciences; Catalog No: 559565), mouse anti-ZO-1 (1:200; Invitrogen; Catalog No: 339100). The following secondary antibodies and fluorescent conjugates and the dilutions were used in this study: Alexa 488 conjugated goat anti-rabbit IgG and goat anti-mouse IgG antibodies (1:250; Invitrogen; Catalog No: A-11034 and A-11029); Alexa 488 conjugated goat anti-chicken IgG (1:250; Invitrogen; Catalog No: A11039); Cy3- and Cy5-conjugated goat anti-rabbit IgG antibody (1:750; Jackson ImmunoResearch; Catalog No: 111-165-144 and 111-175-144); Cy3- and Cy5-conjugated goat anti-mouse IgG antibody (1:750; Jackson ImmunoResearch; Catalog No: 115-165-146 and 115-175-146).

### Image Acquisition

#### Live Imaging

Brightfield images were taken on stereomicroscope (Olympus MVX10), after anesthetizing the embryos with 0.04% MESAB (Sigma-Aldrich, Catalog No: E10521) and mounting in low melting agarose (Sigma-Aldrich, Catalog No: A9414) on a glass slide.

Live imaging to assess the effectiveness of our physical hypoxia regime was performed using Hypoxia Sensor- Image-iT™ Red Hypoxia Reagent (10µM; Invitrogen; Catalog No: H10498). The embryos were incubated for the last 1 hour of the treatment with the dye and fluorescent images were acquired immediately after from the dorsal head epidermis with embryos mounted in a home-made imaging mould with a screw-cap seal for hypoxic embryos using an Olympus FV3000 at 60x/1.40 oil immersion objective lens and 2.4x optical zoom. Embryos were imaged in fresh E3 without methylene blue with 0.04% MESAB. Images were captured in a 12-bit depth in a 512 × 512 grid with a 4µs pixel dwell time.

#### Fixed Imaging

Fluorescent confocal images were acquired from the dorsal head epidermis of zebrafish embryos on Zeiss LSM 980 confocal microscope with a PlanApo 63x/1.40 NA oil immersion objective lens at 1.5x optical zoom and z-step of 0.37µm (Fig. 4K; 5A,D; 6A,D; 7A,D; Supp. Fig. 3A,D) and Olympus FV3000 at 60X/1.40 NA oil immersion objective lens and 2.4x optical zoom (Fig. 1C; 2A,E,G,H; 3A,D; Supp. Fig. 2E) and z-step of 0.37µm for fixed imaging. A z-step of 0.28µm for E-cadherin quantification (Fig. 3A) and 0.37µm for E-cadherin quantification in the recovery experiment (Fig. 3E) was used. A PlanApo 40X/1.3 NA oil immersion objective lens at 0.6x optical zoom was used for low magnification imaging of rosettes in the Rockout treatment experiments. All fixed images were acquired with a resolution of 1024 × 1024, with 4 times frame averaging. All images acquired on Olympus FV3000 (previously mentioned) were imaged and quantified at 12-bit depth; on Zeiss LSM980 (previously mentioned) were imaged and quantified at 16-bit depth.

The pinhole was kept at 1 Airy Unit for both live and fixed imaging. The PMT gains were set using the condition giving the brightest signal to prevent saturation and were kept constant within the experiments.

#### Quantification

Fiji image analysis software was used for image analysis and quantification.

For measuring surface area of plasma membrane, *Tg(cldnB:lynEGFP)* embryos were either treated with hypoxia or normoxia and fixed at different stages and stained with anti-GFP to obtain total surface area and basolateral area, respectively. To quantify surface area, cell outlines were traced in each slice by monitoring the Measure Stack plugin in ImageJ. The perimeter in each slice was multiplied by the slice thickness and added to the area of the first and last slice to obtain the total surface area. To quantify the area of the micro-ridges, images were smoothened and thresholded such that edges of the ridges were neatly defined. This was followed by the Analyze Particle command to detect and measure edge lengths. The perimeter of all the edges was summed up and multiplied with slice thickness and number of slices the ridges extend to in order to get an estimate of the area sequestered in the membrane folds. Five cells from five embryos were analyzed for each condition. The apical area was further determined by subtracting the basolateral surface area from the total surface area.

The quantification and analysis of E-cadherin levels in the periderm cells was based on previously published protocol <u>(Arora, P. et al, 2020)</u>. Briefly, the cell boundaries across z-stacks were marked using the segmented line tool with a fixed width of 7 pixels to cover the cell membrane and the levels were measured at each z-level. For each condition, 4–6 non-neighbouring cells were quantified from each embryo. 7–10 embryos were quantified with a minimum of 30 cells per set. Note that quantification was done where the lateral domains were clearly visible.

For pNM-II and ZO-1 intensity quantification, ROIs were drawn using the segmented line tool with a fixed width of 5-pixels at the apical junctional cortex using ZO-1 as a membrane marker. Intensities were measured in the ZO-1 channel and then on the pNM-II channel for the same ROI. The mean intensity per slice was measured and the average per cell was calculated and plotted. For each condition, 1-4 non-neighbouring cells were quantified from each embryo. 8-12 embryos were quantified and embryo-wise data was plotted. Where required, representative images have been displayed with 15-bit depth to improve signal-to-noise but all quantifications have been performed on the raw data with 16-bit depth.

For cross-sectional area quantification (Supp. Fig. 2A), ZO-1 was used as a membrane marker. Briefly, ROIs were drawn using the polygon tool in the ZO-1 channel and the area was measured. For each condition, 1-4 non-neighbouring cells were quantified from each embryo. 8-10 embryos were quantified and embryo-wise data was plotted.

The quantification of delamination events (Fig. 5G) was carried out by counting the number of embryos that display rosettes accompanied with intercellularly (between the two layers) localized fragmented nuclei assessed using DAPI.

### Statistical analysis

Quantitative analysis of cell morphological features (Fig. 2B,C,D,F; 3G,H; Supp. Fig. 1A,B) and cortical intensities of E-cadherin (Fig. 3I, Supp. 1G), ZO-1 (Fig. 3K) and pNM-II (Fig. 4B) was carried out using R and RStudio <u>(Sen, S., et al 2026)</u>. The data were compared using statistical tests such as Mann-Whitney U test and Kruskal Wallis test with the appropriate post hoc tests. The box-jitter plots compare the median amongst conditions, and it ranges from 25th and 75th percentile while the error bar represents the 95% confidence interval.

Quantitative analysis of cortical intensity obtained (Fig. 4D,G; 5B,E; 6B,E; 7B,E; Supp. Fig. 2D,G) was carried out using Python v3.9.12, linear mixed-effects modeling was employed to account for both fixed experimental factors and hierarchical data structure. Briefly, mean intensity values were first calculated at the embryo level by averaging measurements across quantified cells within each embryo. These embryo-level values were used as the unit of analysis. All data represented are pooled from at least three independent replicate experiments, with replicate number and sample size listed in each figure legend.

A linear mixed-effects model (LMM) described by the following equation:

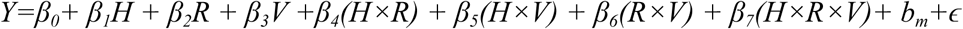

where Y is observed log_2_ intensity, H and R correspond to Hypoxia and Rockout, and V represents the genetic condition (EpCAM MO/Mypt1 MO/*crb3a;3b* m/z) used for the experiment. *b_m_* accounts for the random clutch effects, *ɛ* corresponds to residual noise and *β* are the model coefficients with *β_0_*being the baseline coefficient, was fitted to assess combinatorial effects on protein localization. Estimated marginal means (EMMs) are calculated from the fitted linear mixed-effects model by predicting the mean response for each experimental condition while accounting for the effects of all model terms and clutch-to-clutch variability. The EMMs therefore represent model-adjusted condition means rather than raw observed averages or medians. Data were analysed using linear mixed-effects models implemented in Python (**statsmodels**), with experimental conditions treated as fixed effects and clutch included as a random intercept to account for inter-clutch variability. EMMs and their 95% confidence intervals were obtained from the fitted model using the fixed-effect parameter estimates and covariance matrix. Pairwise comparisons between EMMs were performed using Wald z-tests and resulting p-values were corrected for multiple comparisons using the Benjamini–Hochberg false discovery rate (FDR) procedure. Holm correction was also evaluated and yielded qualitatively similar trends, but FDR-adjusted p-values were reported to balance sensitivity with control of false discoveries. The box-jitter plots show the median and range from the 25th to the 75th percentile, while the error bar represents the 95% CI. Pairwise treatment effects were visualized as differences in EMMs with 95% confidence intervals, obtained from the linear mixed-effects model using forest plots. All quantified data was plotted in Python.

The code used for data analysis and plotting of graphs using LMM is provided in the Supplementary Information.

All the numerical data used for plotting the graphs in this study are presented in the Dataset S1. The statistical tests and comparisons conducted on the datasets are shown in the Dataset S2.

## Acknowledgements

The authors thank the Advanced Centre for Treatment, Research and Education in Cancer (ACTREC) for their contribution to TEM imaging in this work. The *crb3* mutant lines were shared by Satu Kujawski and Elisabeth Knust from Max-Planck Institute of Molecular Cell Biology and Genetics. We also thank Dr. Kalidas Kohale and Boby K.V. for fish facility and confocal microscope facility maintenance, respectively, and TIFR-DAE (RTI4003: 1303/2/2019/R&D-II/DAE/2079) for funding.

## Author’s contributions

Performed experiments, discovered the phenotype and optimized experiments, M.F.; prepared data analysis code, A.K; conceived and conceptualized the project and interpreted the data, M.F and M.S.; secured the funding and supervised the project, M.S.; wrote the manuscript, M.F and M.S.

---

**Supplementary Figure 1:**
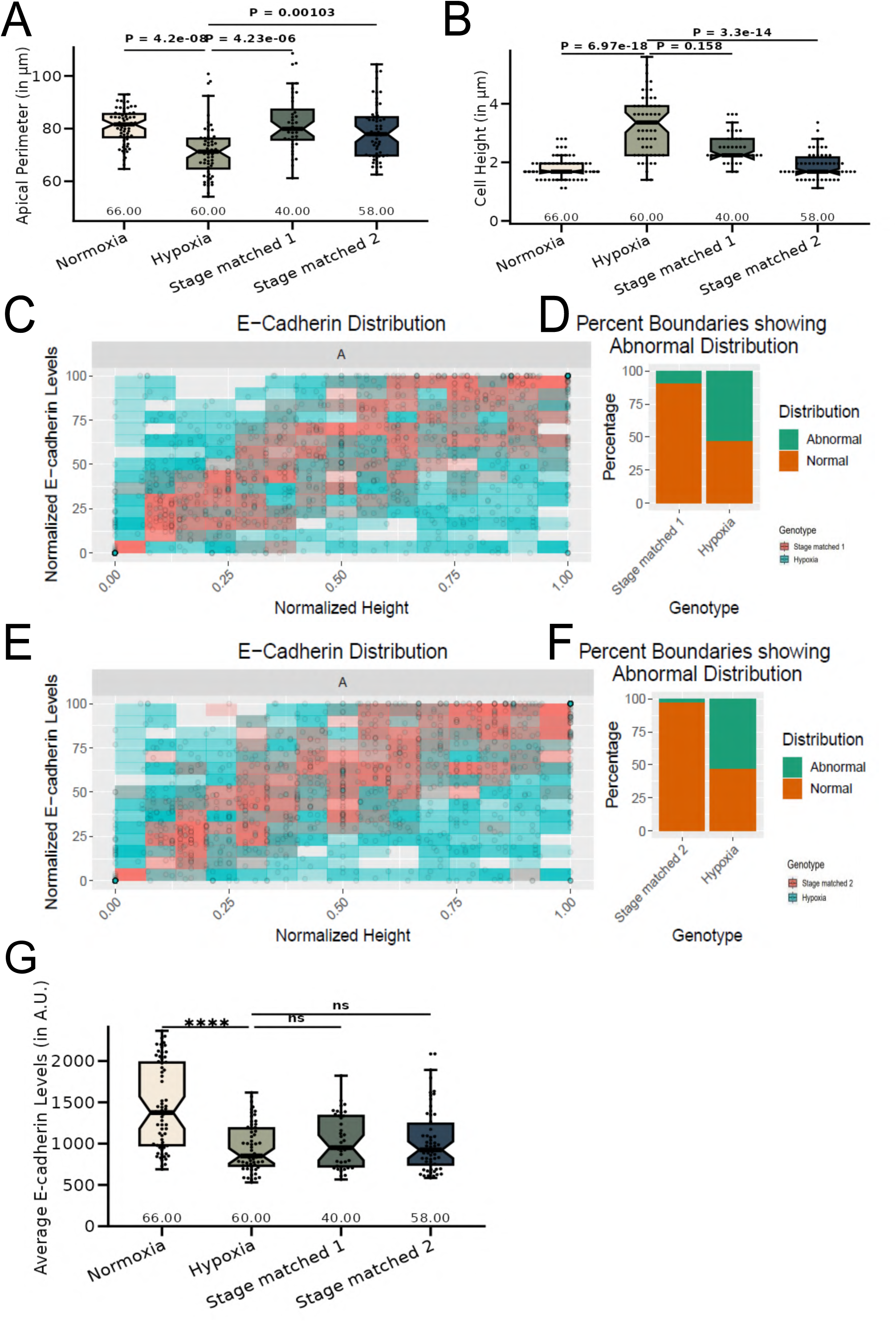
Epithelial architectural changes in the hypoxic embryos relative to the stage matched controls. (A,B) Estimation and comparison of apical perimeter (A) and cell height (B) in the 48hpf normoxic, hypoxic, stage matched 1 (36hpf) and stage matched 2 (42hpf) normoxic embryos in the periderm. Data are median±interquartile range (no of cells assessed per condition=40-66, N=3). \*\**P*<0.01, \*\*\*\**P*<0.0001, n.s. not significant *P*>0.05 [Kruskal Wallis with Dunn’s post-hoc test]. (C,E) Graphical representation showing polarised localisation of E-cadherin across normalised cell height in the periderm of stage matched 1 and hypoxia treated embryos (C) and stage matched 2 and hypoxia treated embryos (E) in the periderm. (D,F) A graph showing noise index in E-cadherin localisation in stage matched 1 and hypoxia treated embryos (D) and stage matched 2 and hypoxia treated (F) in the periderm. (G) Box plot showing comparison of average E-cadherin levels in the 48hpf normoxic, hypoxic, stage matched 1 (36hpf) and stage matched 2 (42hpf) embryos. Data are median±interquartile range (no of cells assessed per condition=40-66, N=3). \*\*\**P*<0.001, \*\*\*\**P*<0.0001, n.s. not significant *P*>0.05 [Kruskal Wallis with Dunn’s post-hoc test].

**Supplementary Figure 2:**
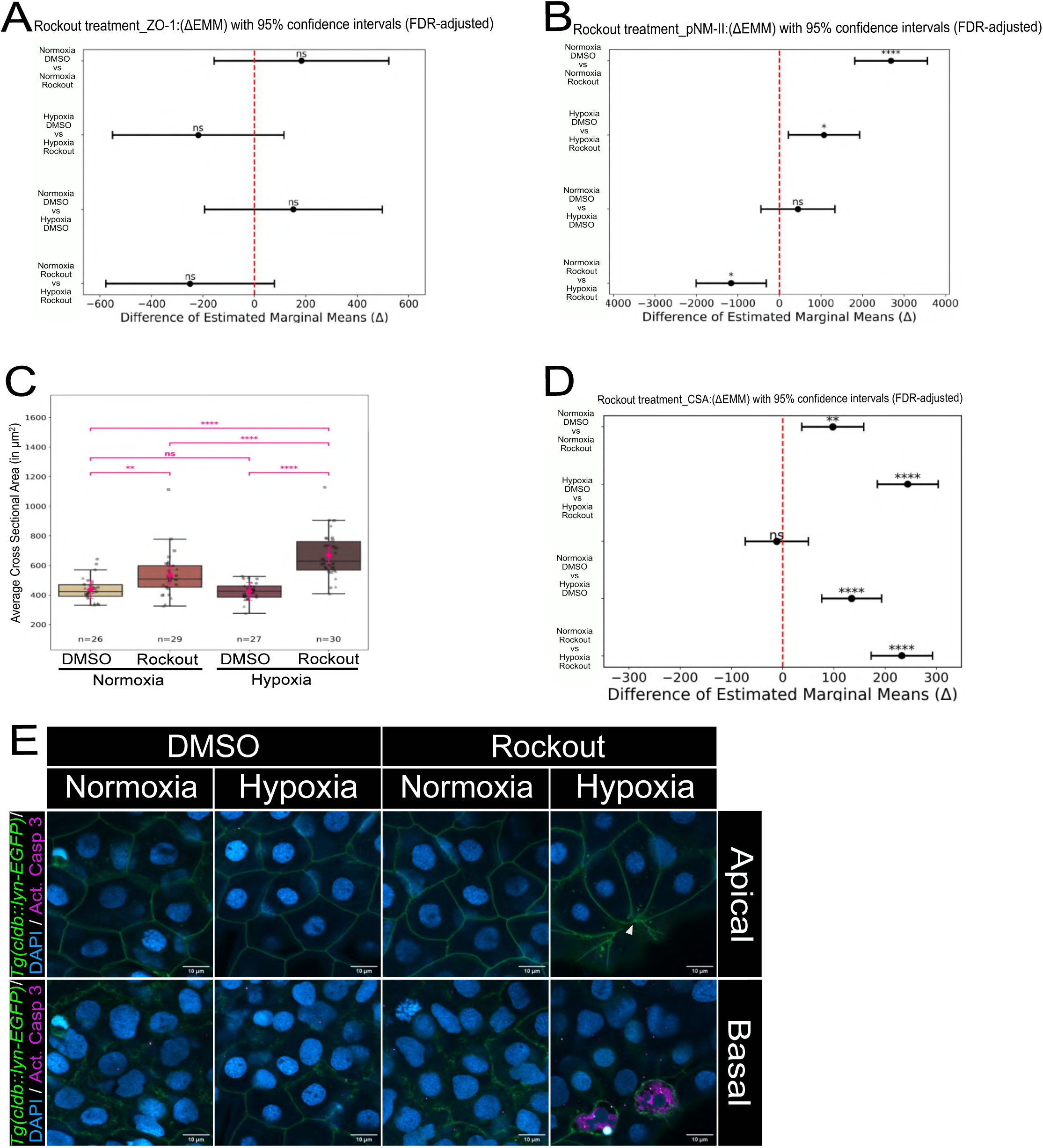
Assessment of ROCK inhibition on peridermal features in hypoxic embryos. (A,B) Forest plots showing pairwise treatment effects in ZO-1 (A) and pNM-II (B) visualized as differences in EMMs with 95% confidence intervals in embryos treated with normoxia and hypoxia with DMSO and Rockout. (C) Boxplot showing the average cross-sectional area in wild type embryos treated with DMSO and Rockout under normoxia and hypoxia. Boxplot represents the median±interquartile range of the raw data (no of embryos assessed for each condition=26-30, N=3). Pink diamonds represent EMMs used for statistical analysis. \*\**P*<0.01, \*\*\*\**P*<0.0001 [Wald z-test]. ( ●, ▴, ◼ represent Set1, 2 and 3, respectively). (D) Forest plot showing pairwise treatment effects on cross-sectional area visualized as differences in EMMs with 95% confidence interval. (E) Confocal images of the apical slice (upper row) and the basal slice (lower row) of the peridermal cells from *Tg(cldnB:lynEGFP)* stained for Activated Caspase 3 and DAPI treated with Rockout and vehicle control embryos under normoxia and hypoxia. White arrowheads indicate delamination events and associated apoptotic cells marked with LynEGFP and Activated Caspase 3. Scale bars= 10µm in (E).

**Supplementary Figure 3:**
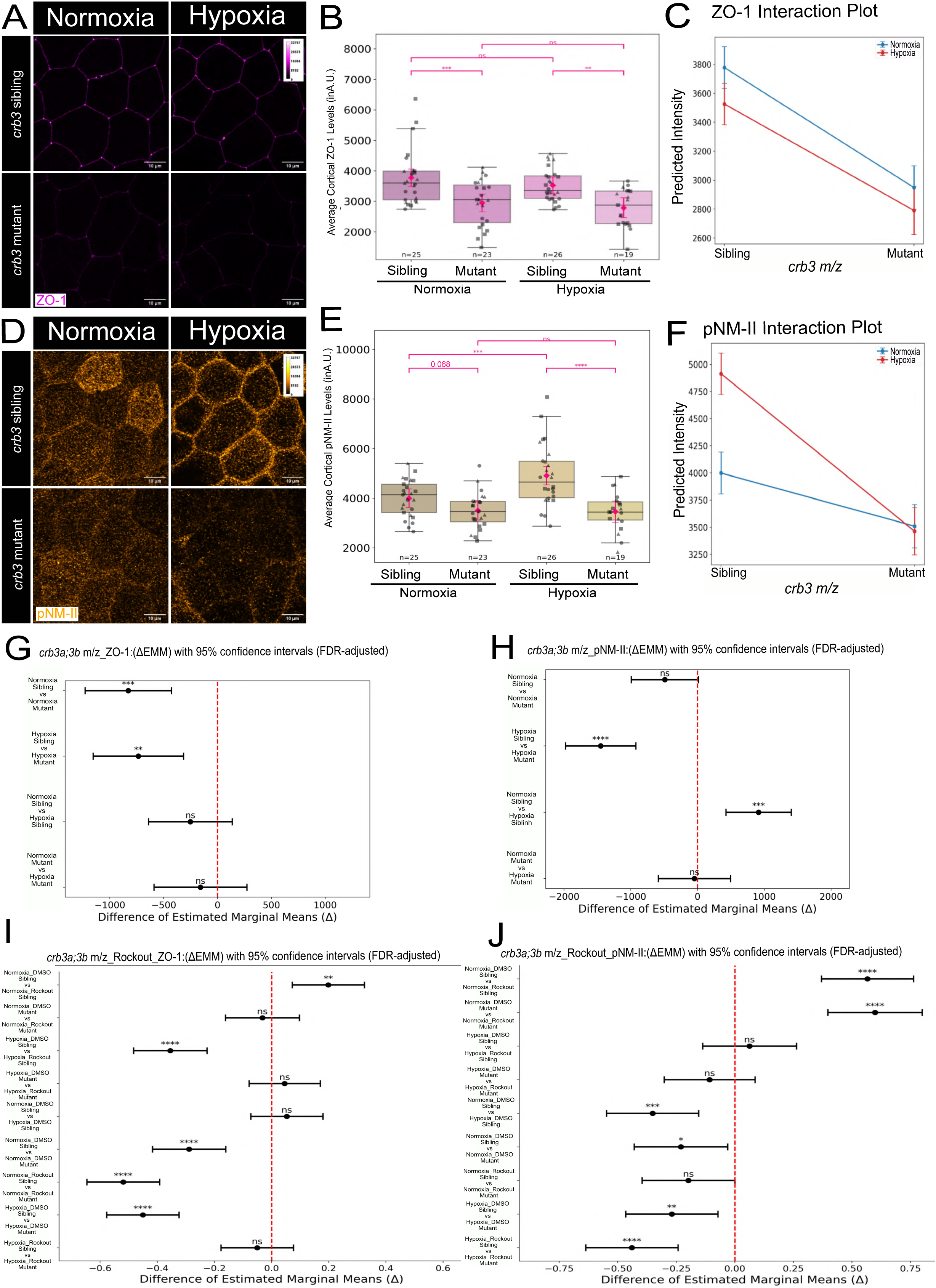
Assessing the impact of loss of *crb3a;3b* function on peridermal ZO-1 and pNM-II localisation. (A,B) Representative confocal images showing immunolocalisation of ZO-1 (A) and pNM-II (B) in the periderm of *crb3* sibling and mutant embryos exposed to normoxia and hypoxia for 24 hours. (B,E) Box plot showing comparison of levels of ZO-1 (B) and pNM-II (E) levels in the periderm of *crb3* embryos under normoxia and hypoxia. Boxplot represents the median±interquartile range of the raw data (no of embryos assessed for each condition=19-26, N=3). Pink diamonds represent EMMs used for statistical analysis. \**P*<0.05, \*\*\*\**P*<0.0001 [Wald z-test]. ( ●, ▴, ◼ represent Set1, 2 and 3, respectively). (E, H) Graphs represent ZO-1 (C) and pNM-II (F) interaction plots to depict trends in *crb3a;3b* sibling and mutant embryos reared under normoxic and hypoxic conditions. (G,H,I,J) Forest plots showing pairwise treatment effects in *crb3* sibling and mutant embryos on ZO-1 following a 24 hour treatment (G) and a 12 hour treatment with Rockout (I) and pNM-II following a 24 hours treatment (H) and a 12 hour treatment with Rockout (J) visualized as differences in EMMs with 95% confidence intervals. \**P*<0.05, \*\**P*<0.01, \*\*\*\**P*<0.0001, n.s. not significant *P*>0.05 [Wald z-test]. Scale bars= 10µm in (A,D). Data represented in 15-bit depth (A,D).

**Supplementary Figure 4:**
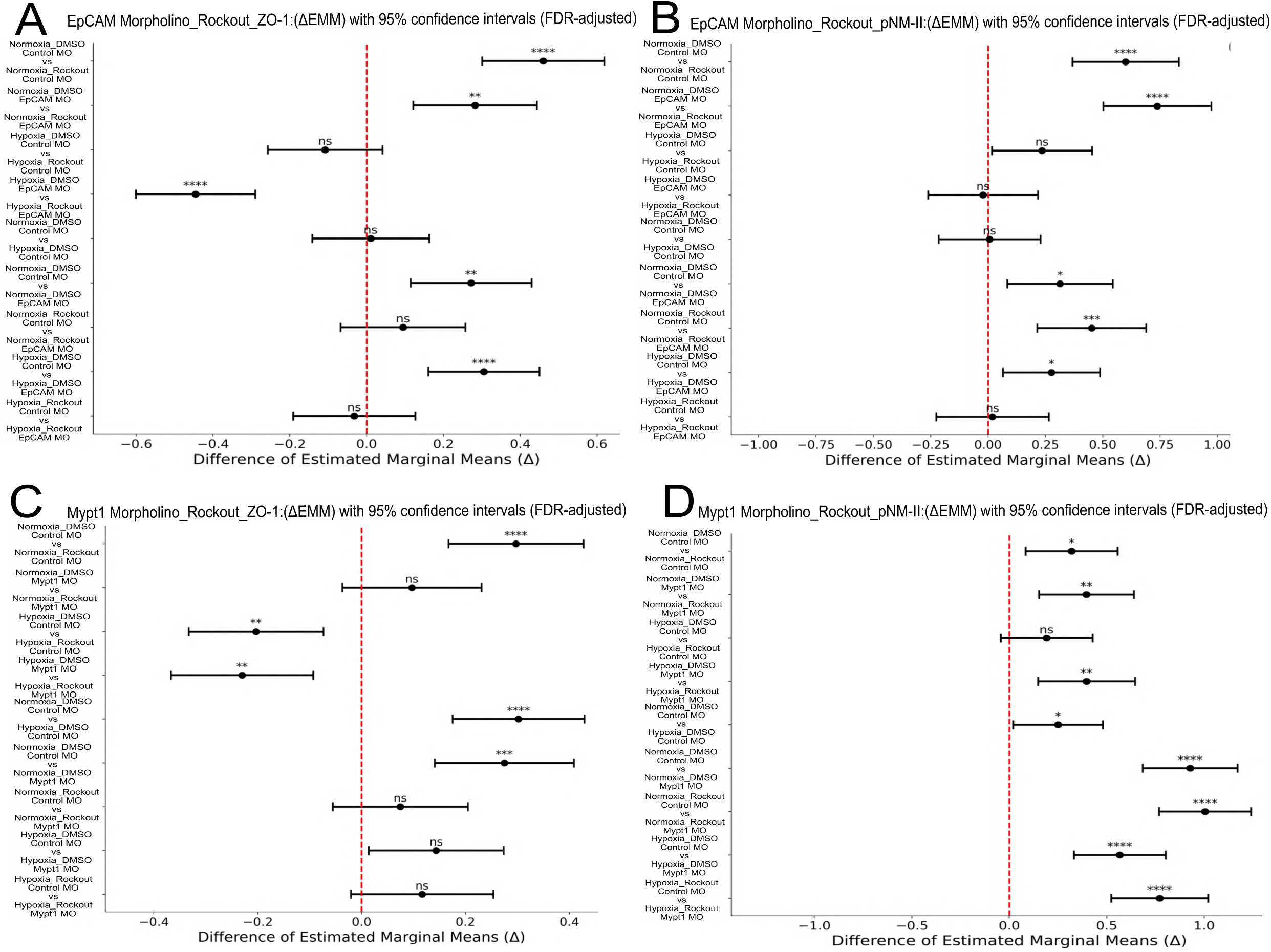
Assessment of EpCAM and Mypt1 knockdown on ZO-1 and pNM-II levels in Rockout treated hypoxic embryos. (A-D) Forest plots showing pairwise treatment effects visualized as differences in EMMs of different conditions for ZO-1 levels in EpCAM morphants (A) and Mypt1 morphants (C), and pNM-II levels in EpCAM morphants (B) and Mypt1 morphants (D) treated with DMSO and Rockout under normoxia and hypoxia. \**P*<0.05, \*\**P*<0.01, \*\*\**P*<0.001, \*\*\*\**P*<0.0001, n.s. not significant *P*>0.05 [Wald z-test].

## Supplementary Information

### CODE FOR LMM ANALYSIS (4 CONDITIONS)

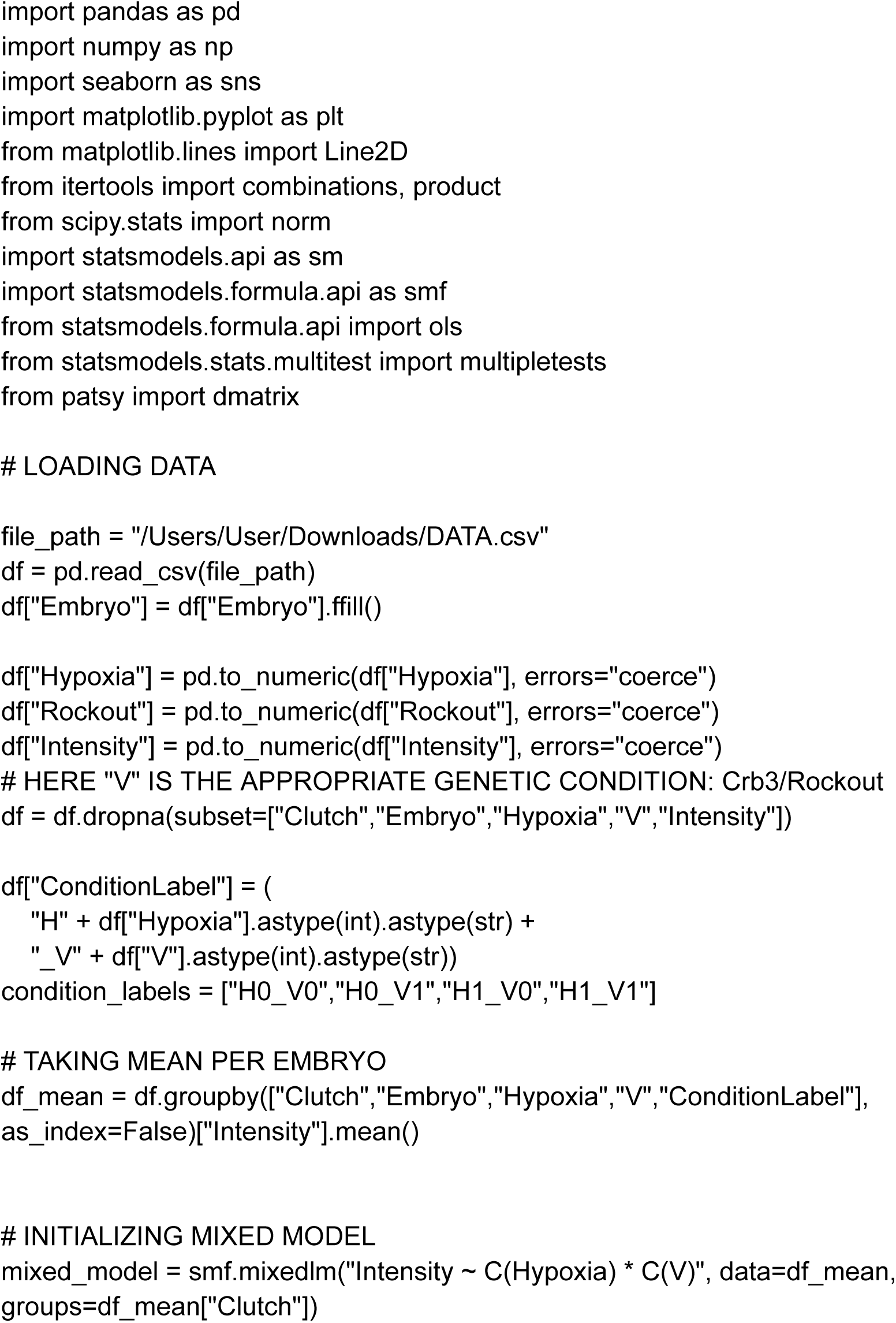
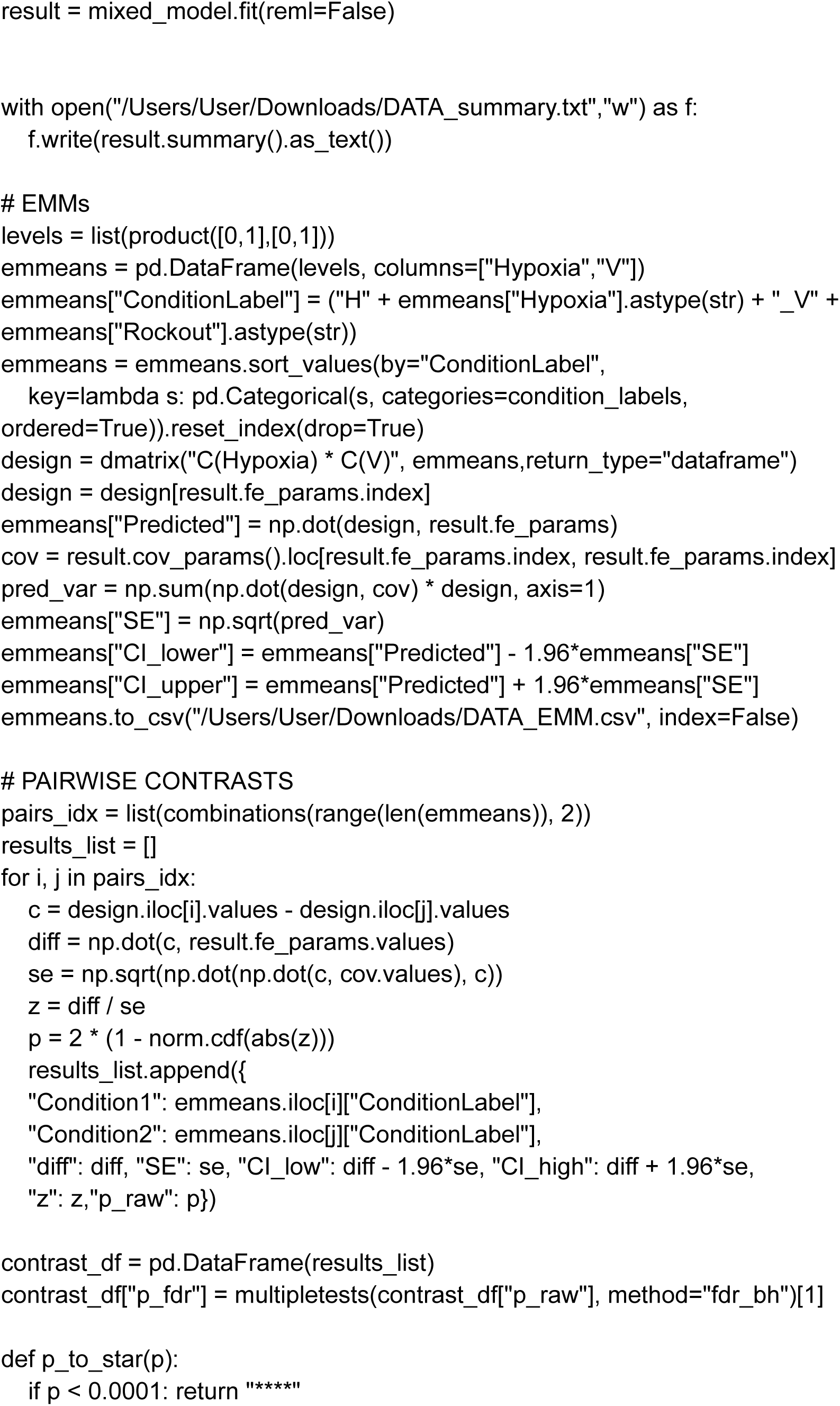
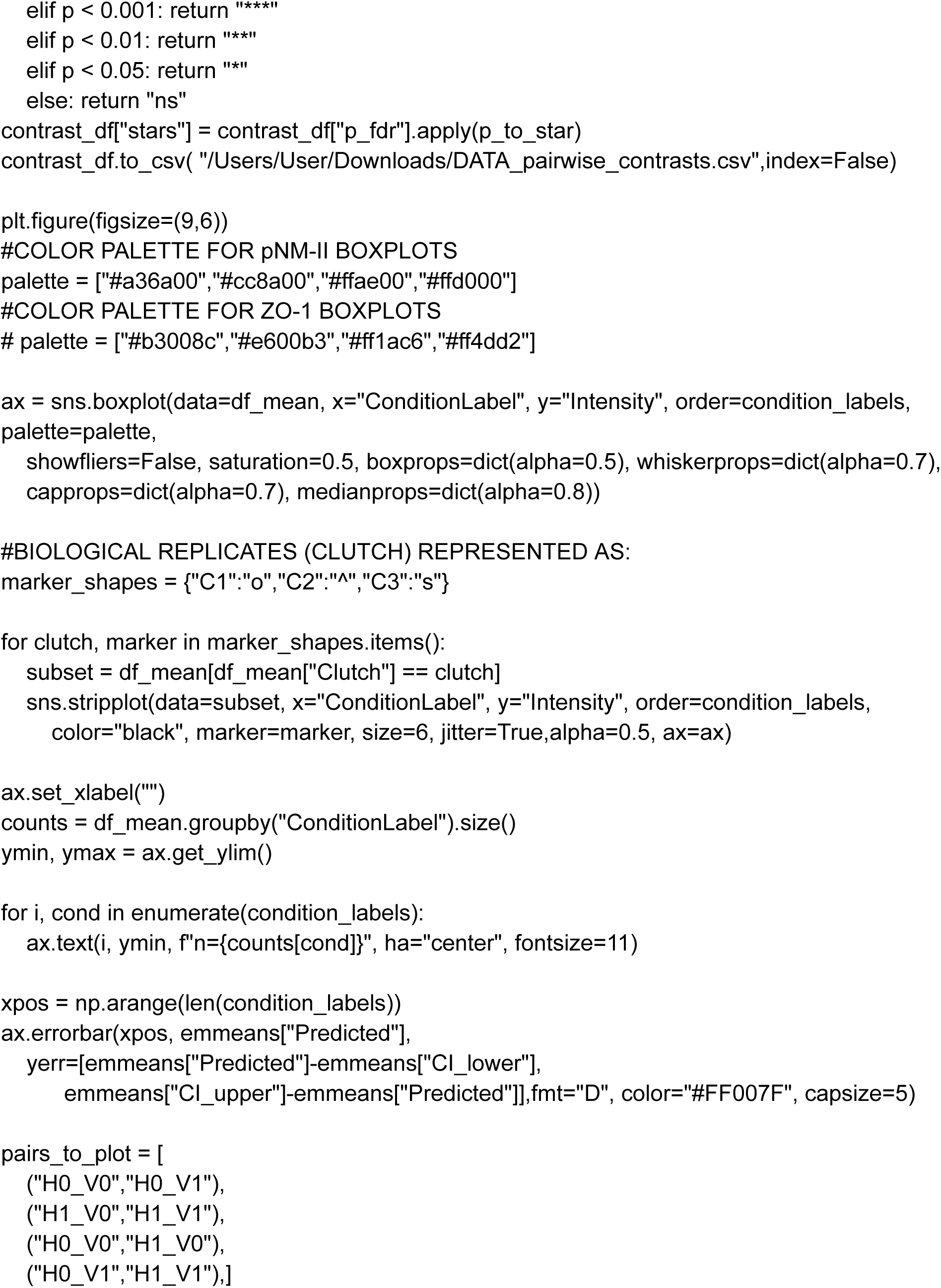
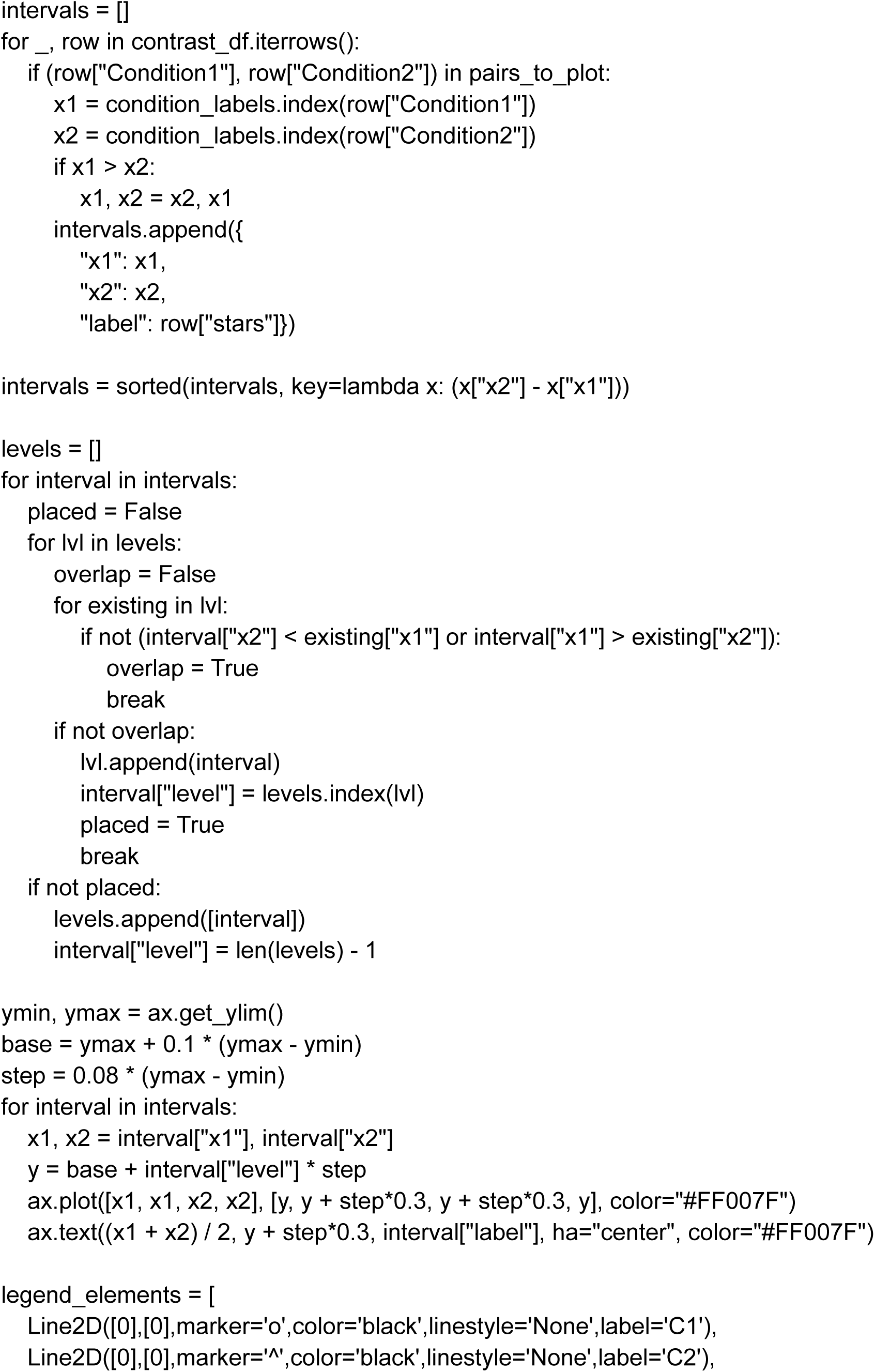
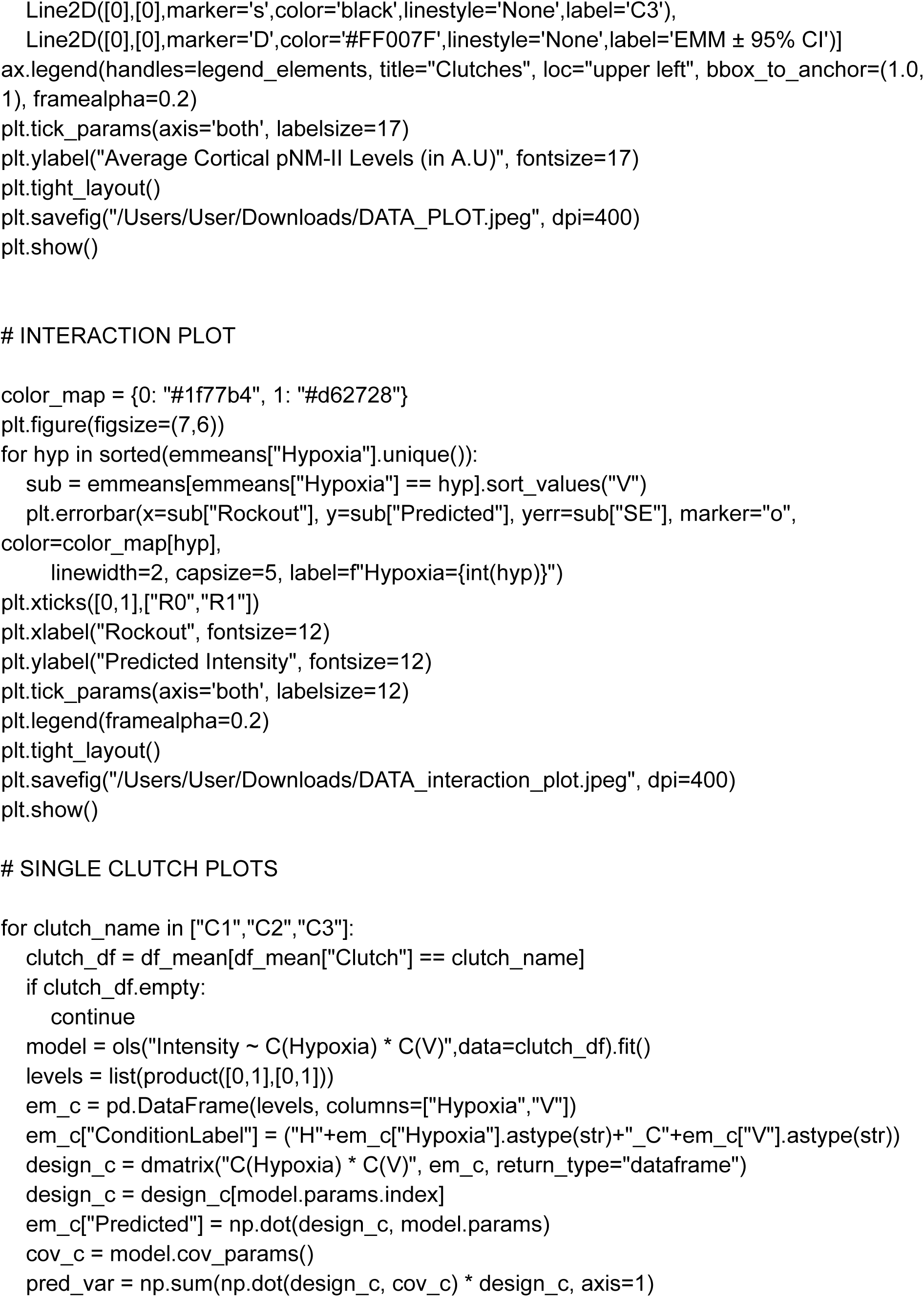
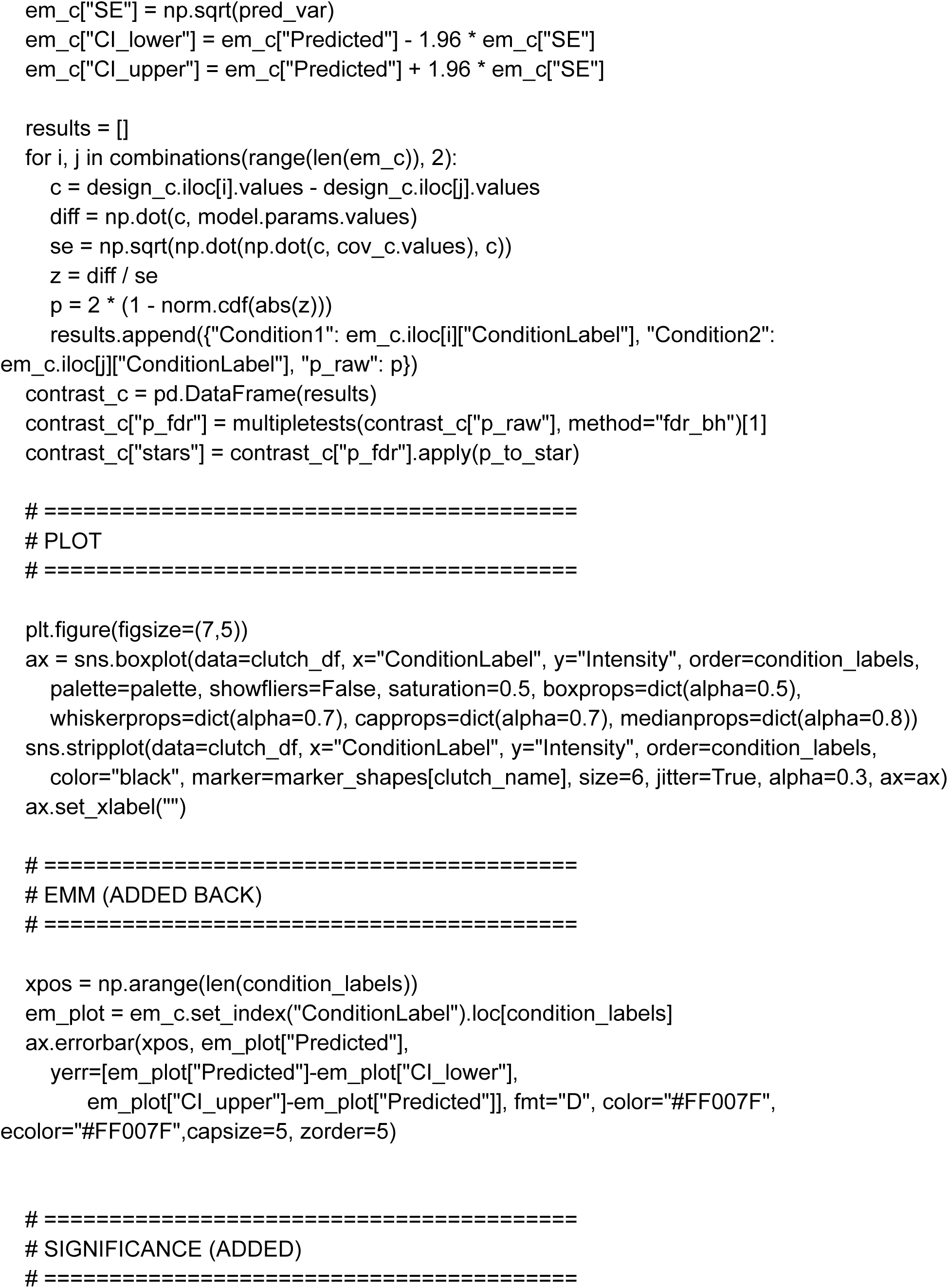
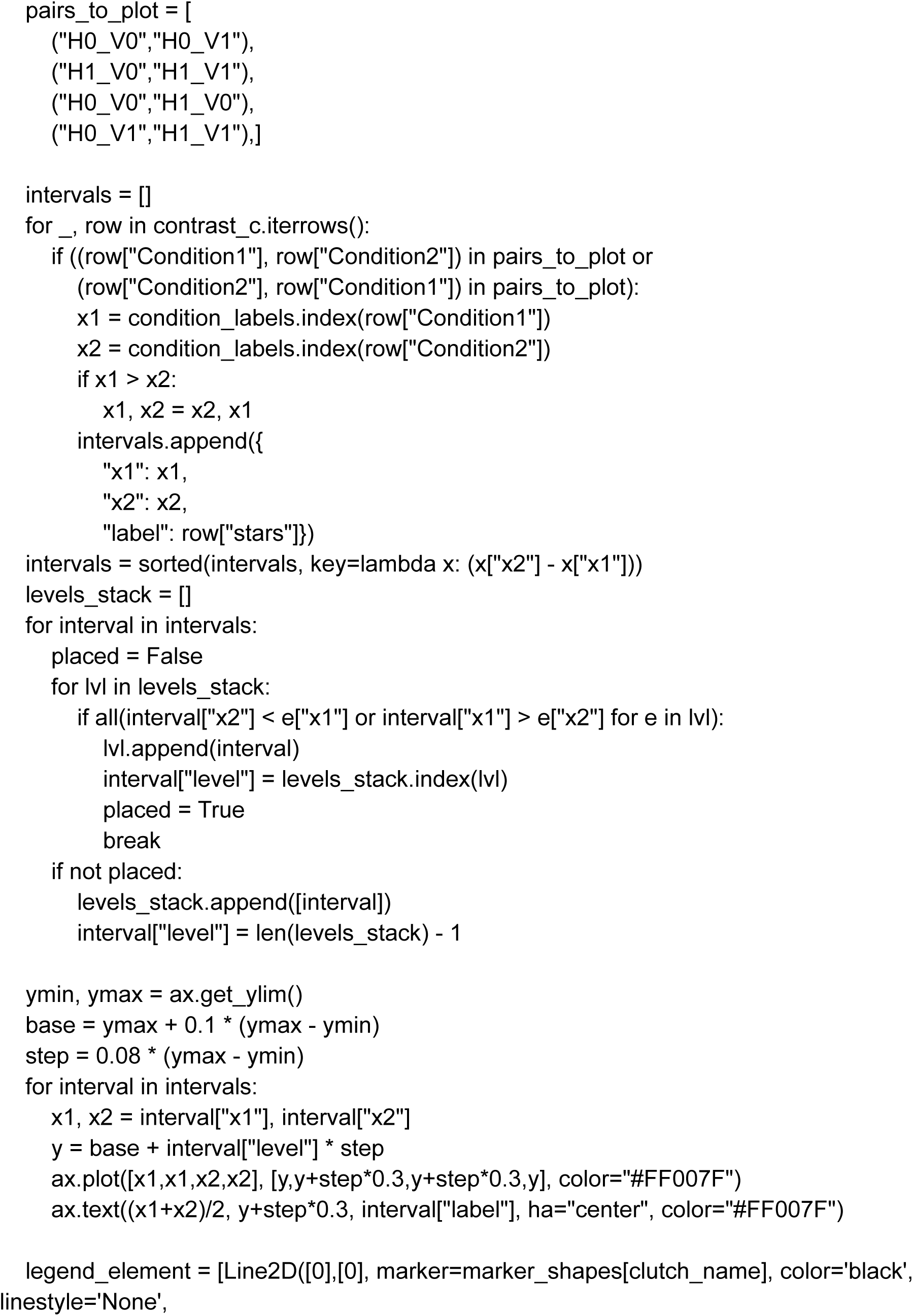
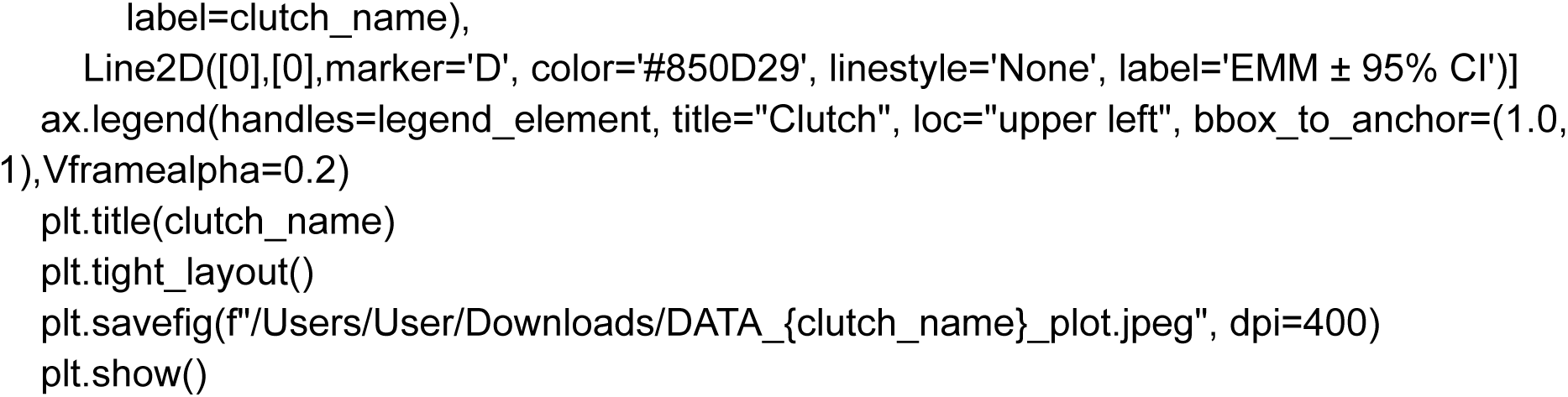

### EMM DIFFERENCE AND FOREST PLOT

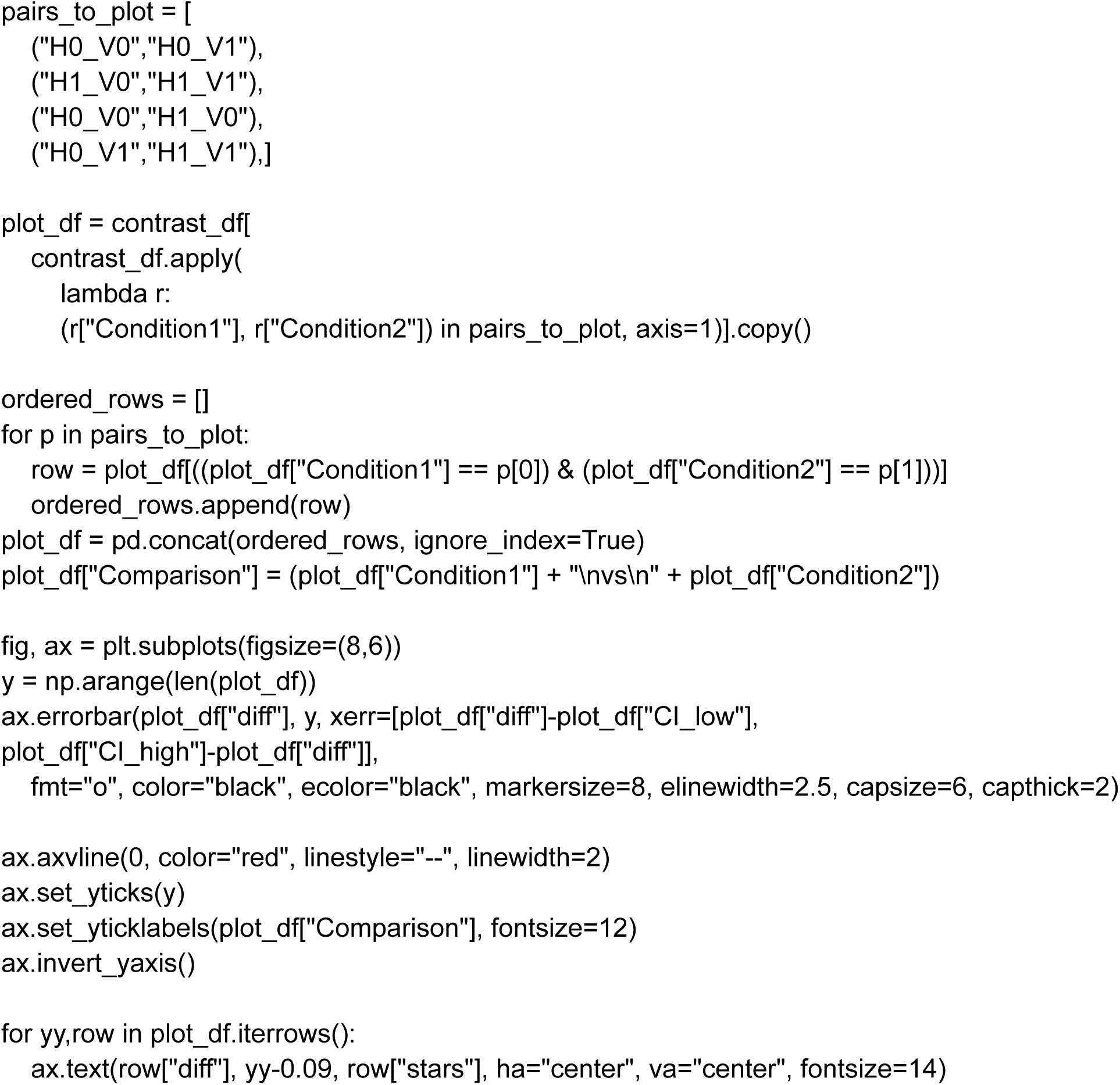
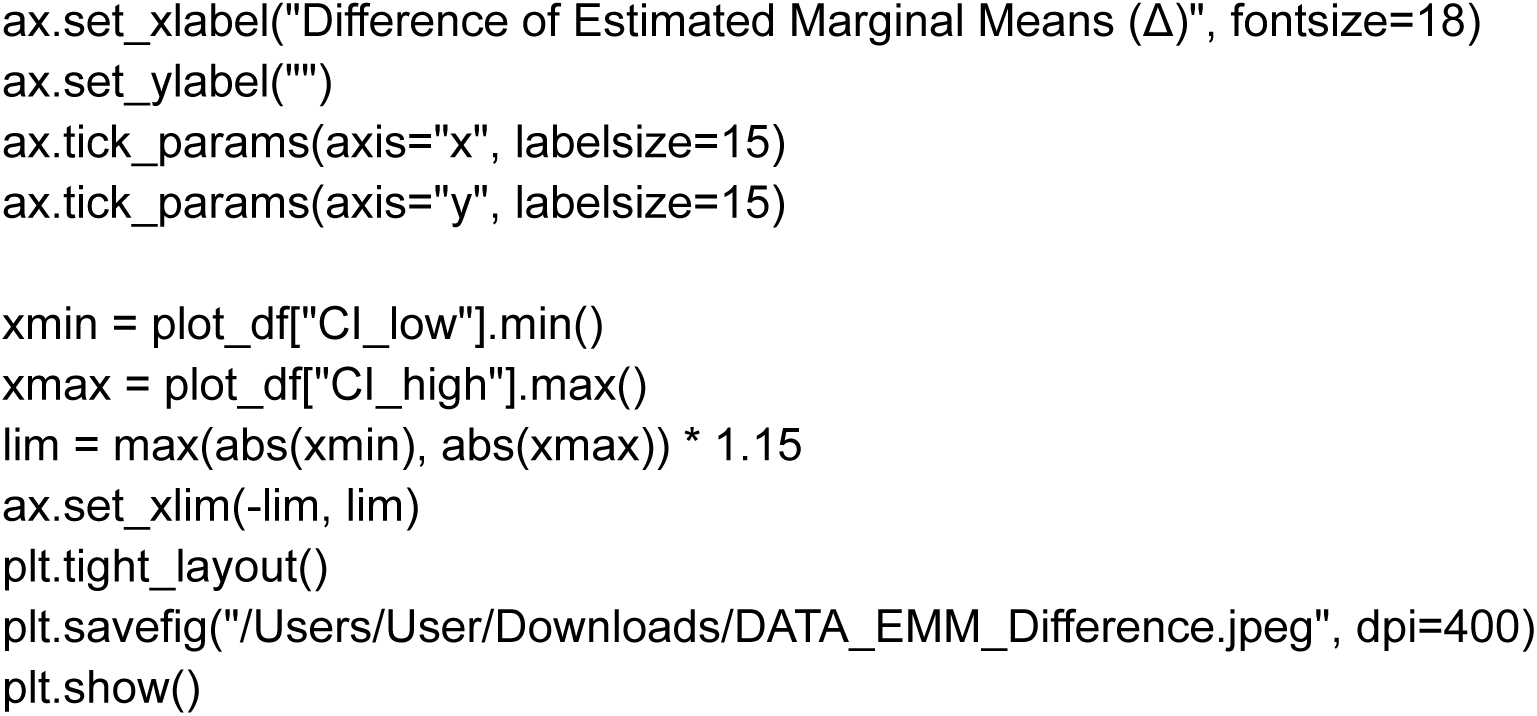

### CODE FOR LMM ANALYSIS (8 CONDITIONS)

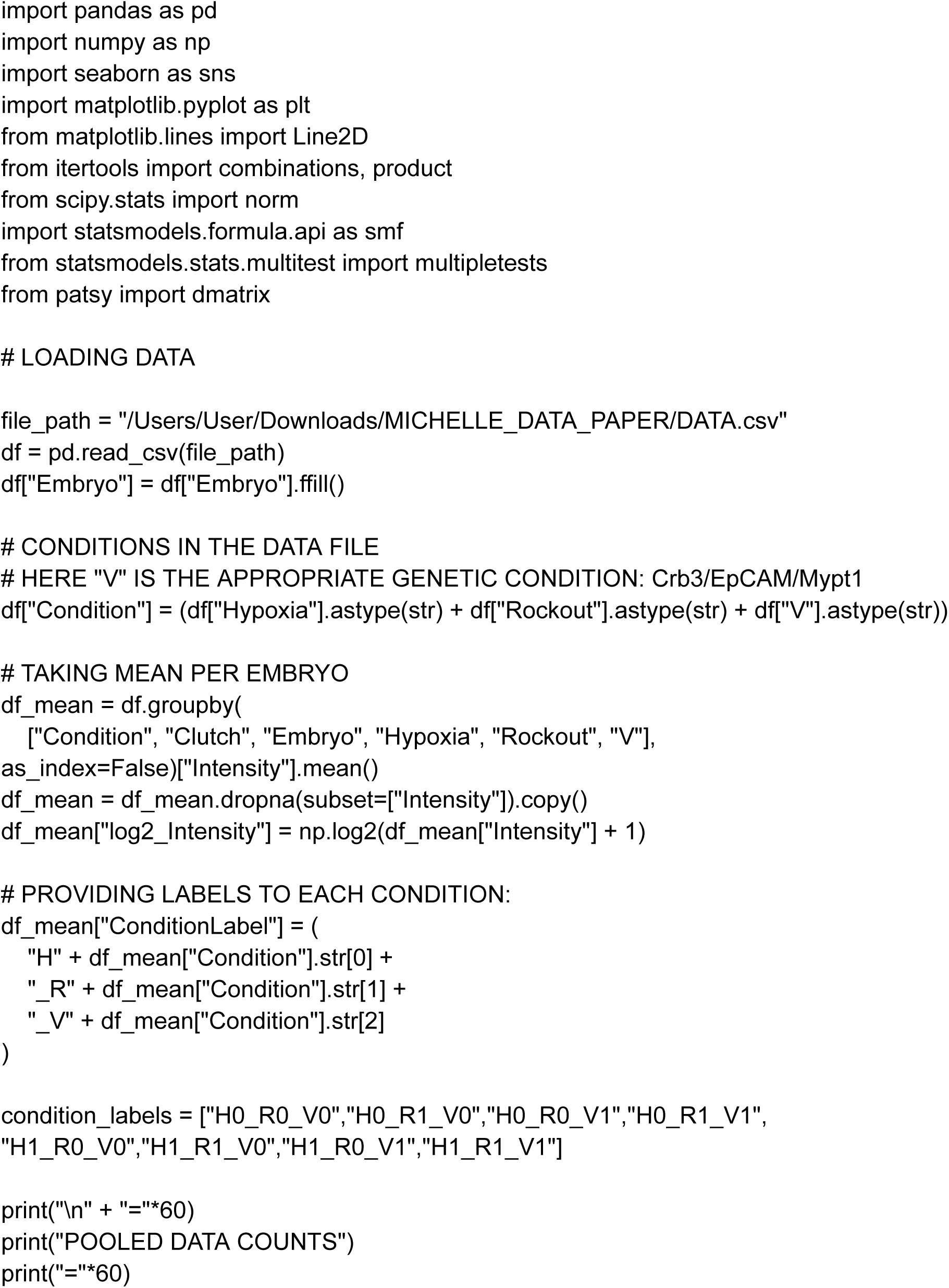
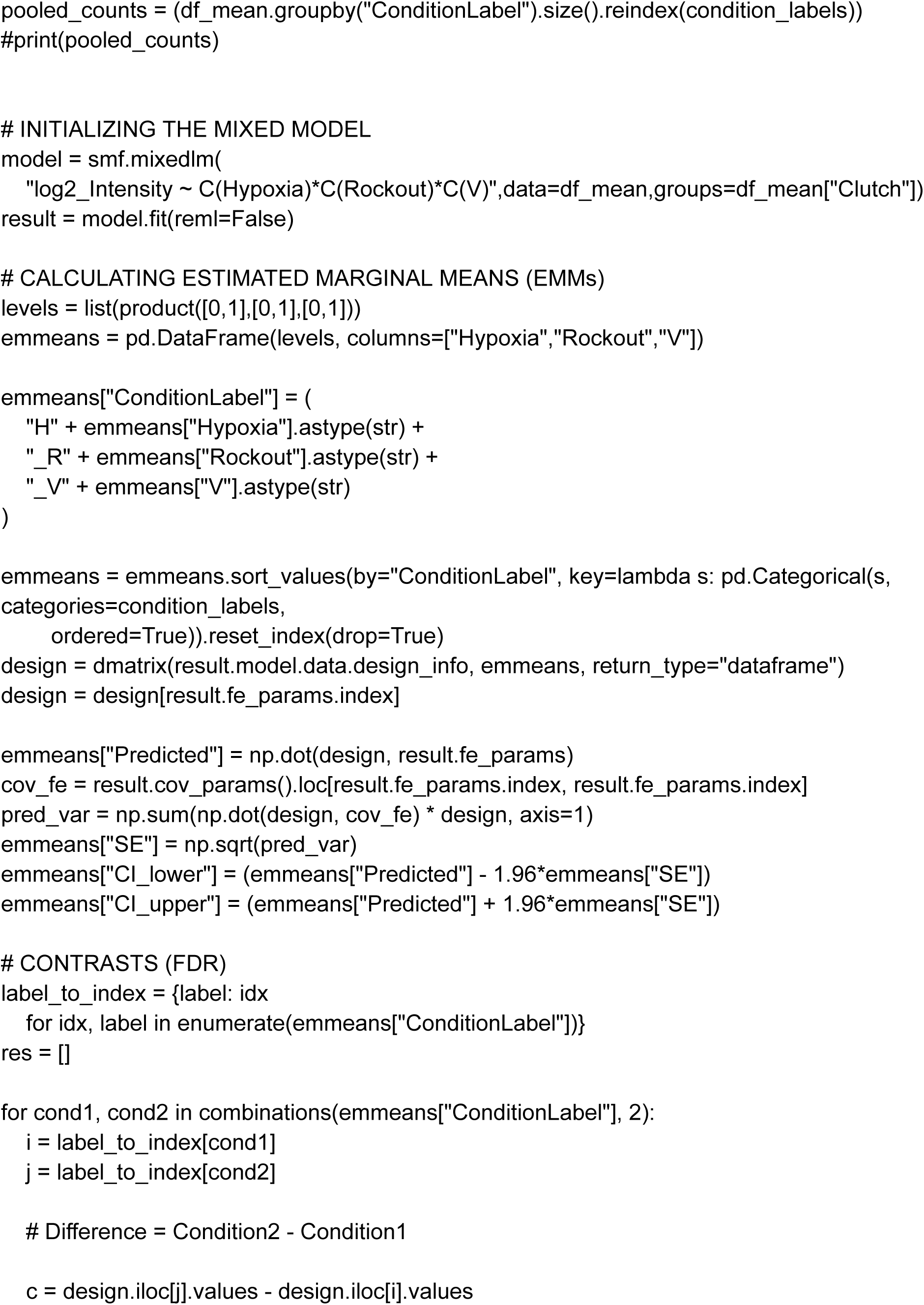
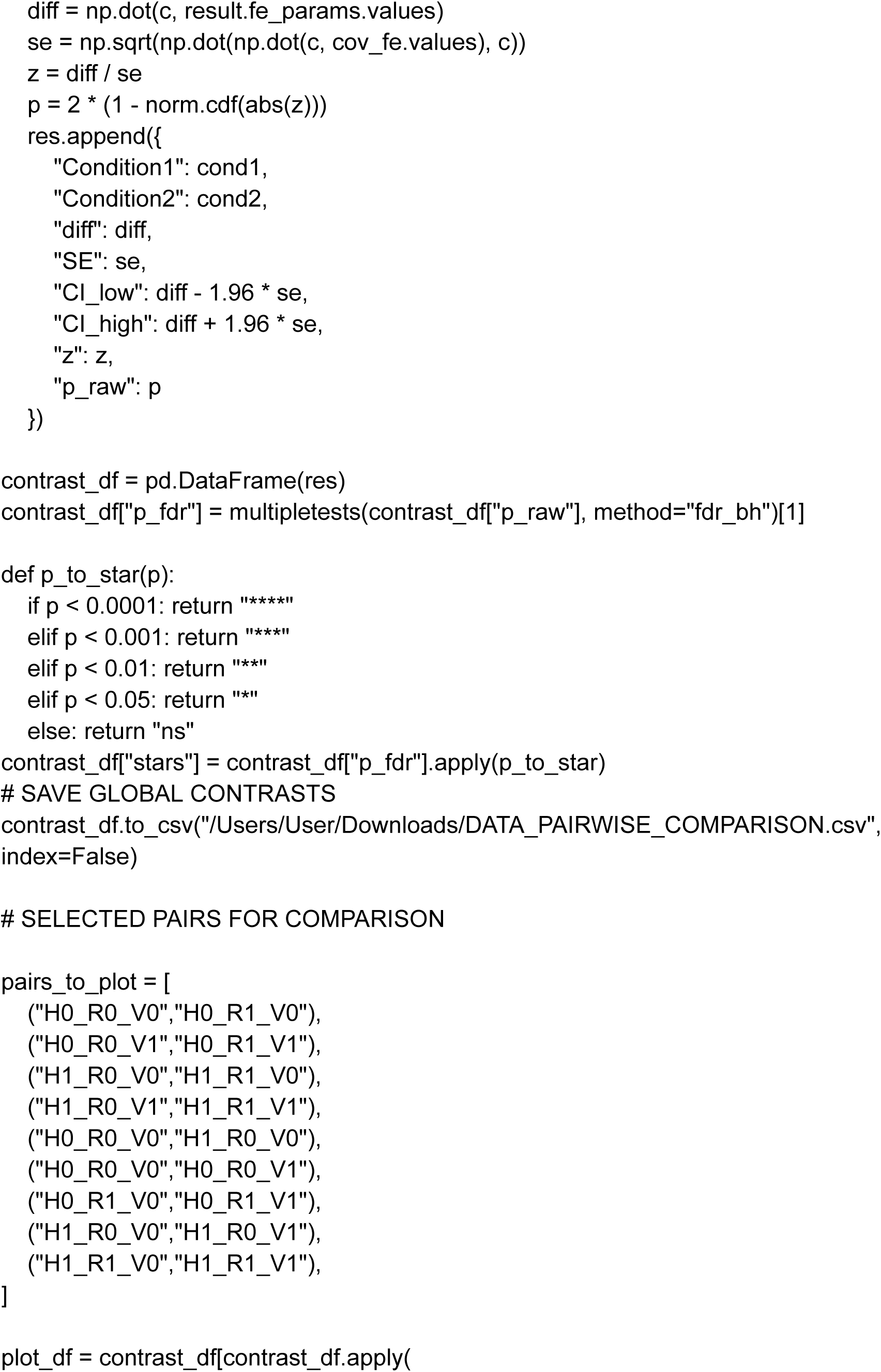
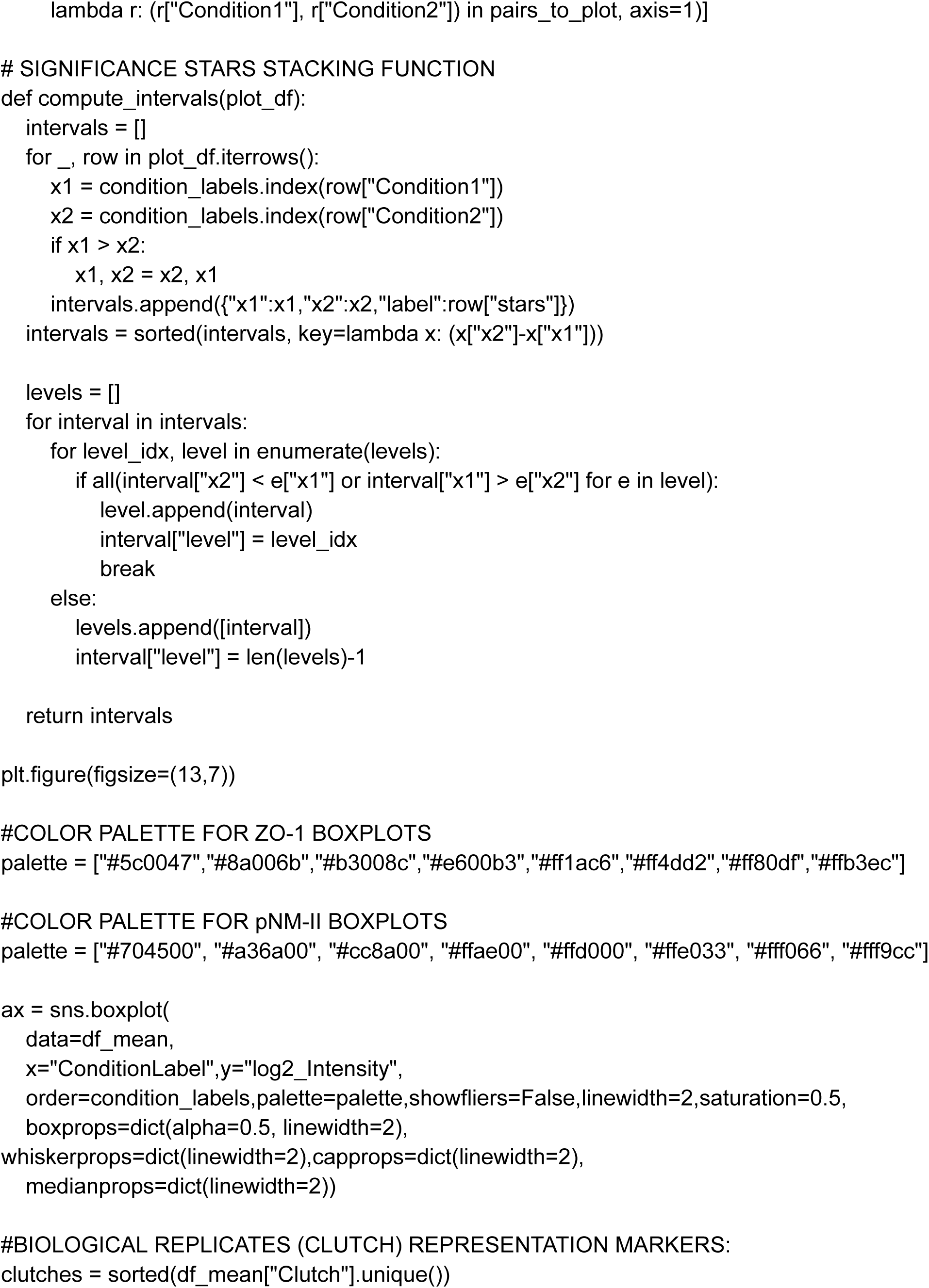
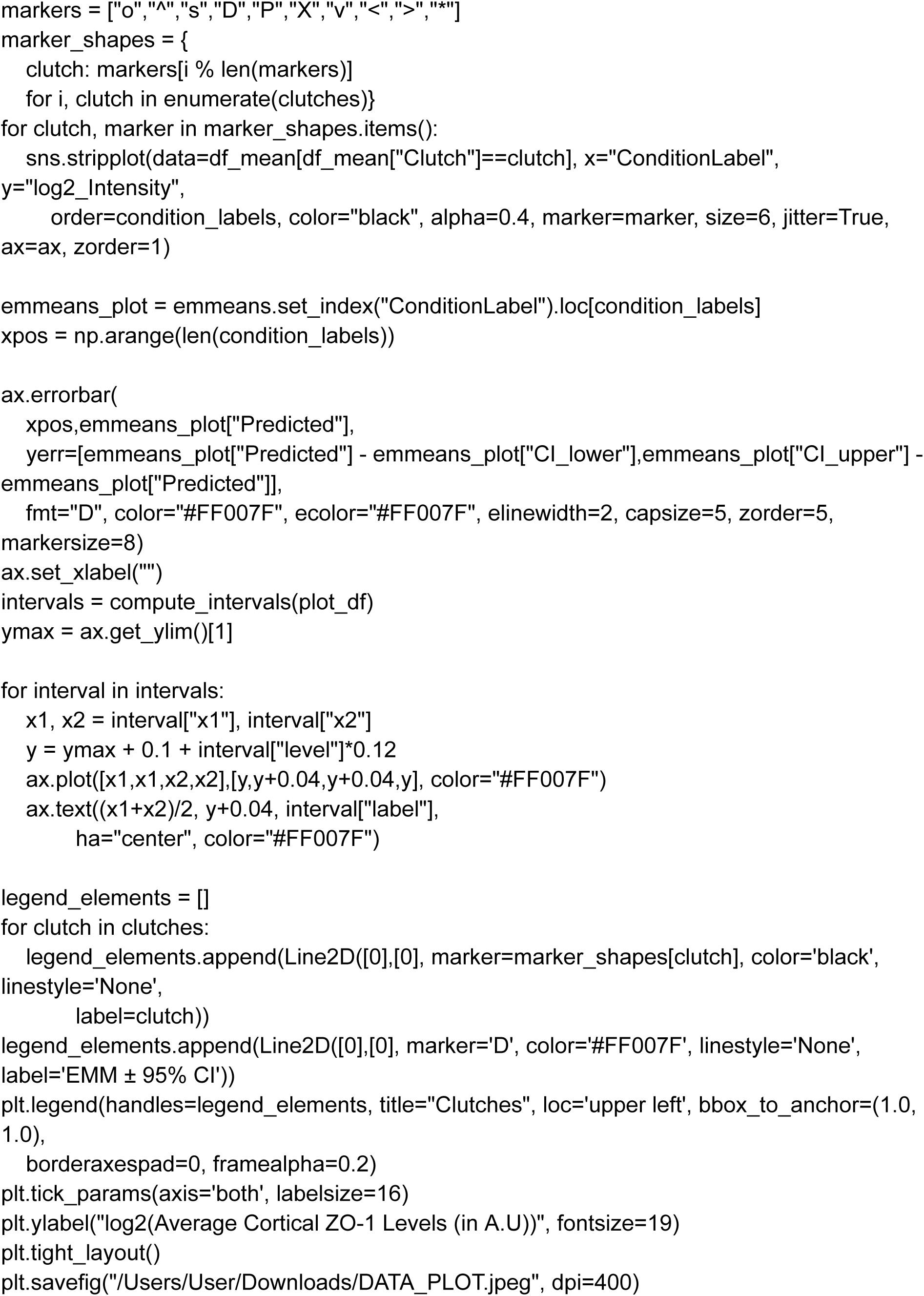
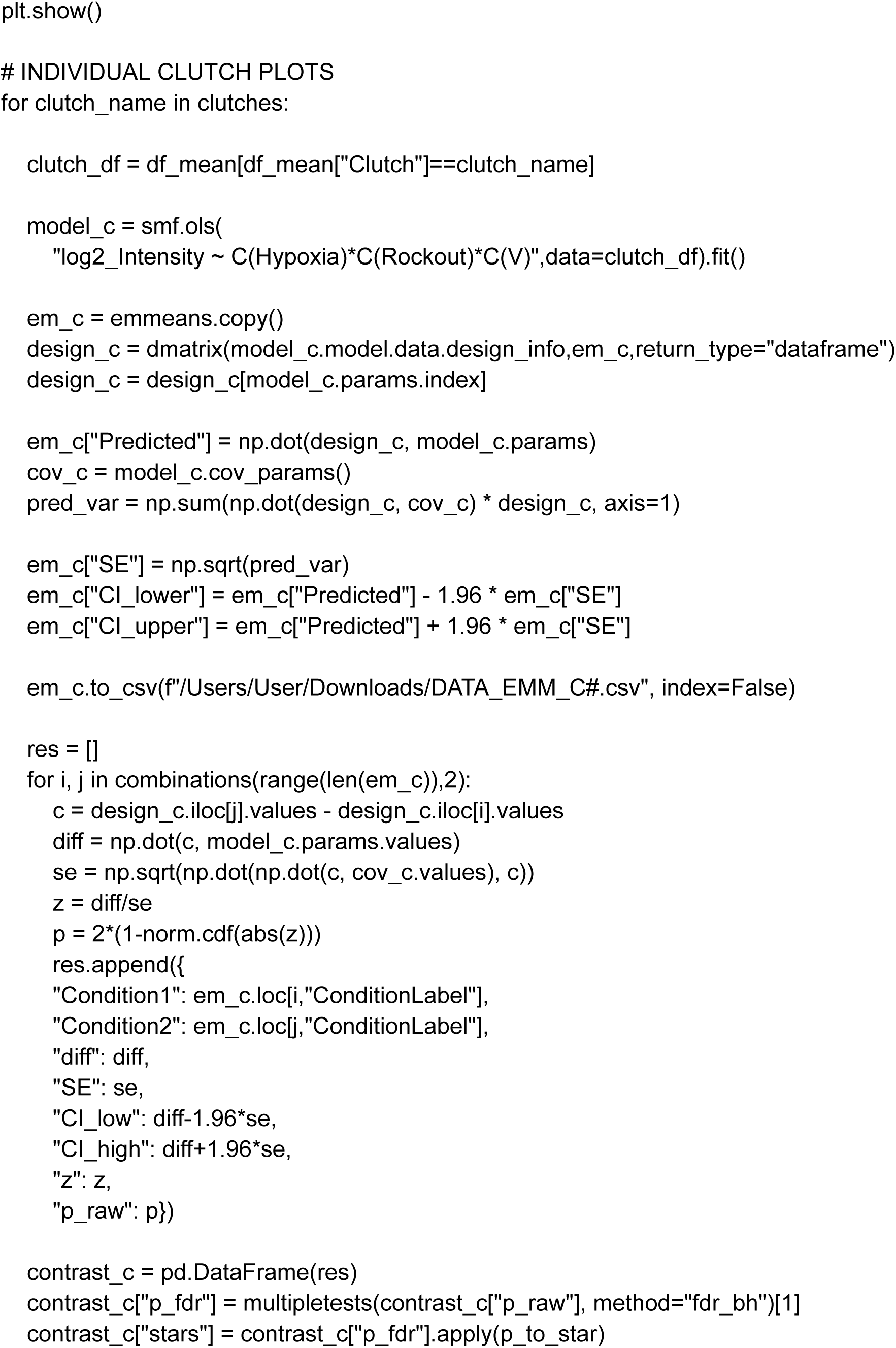
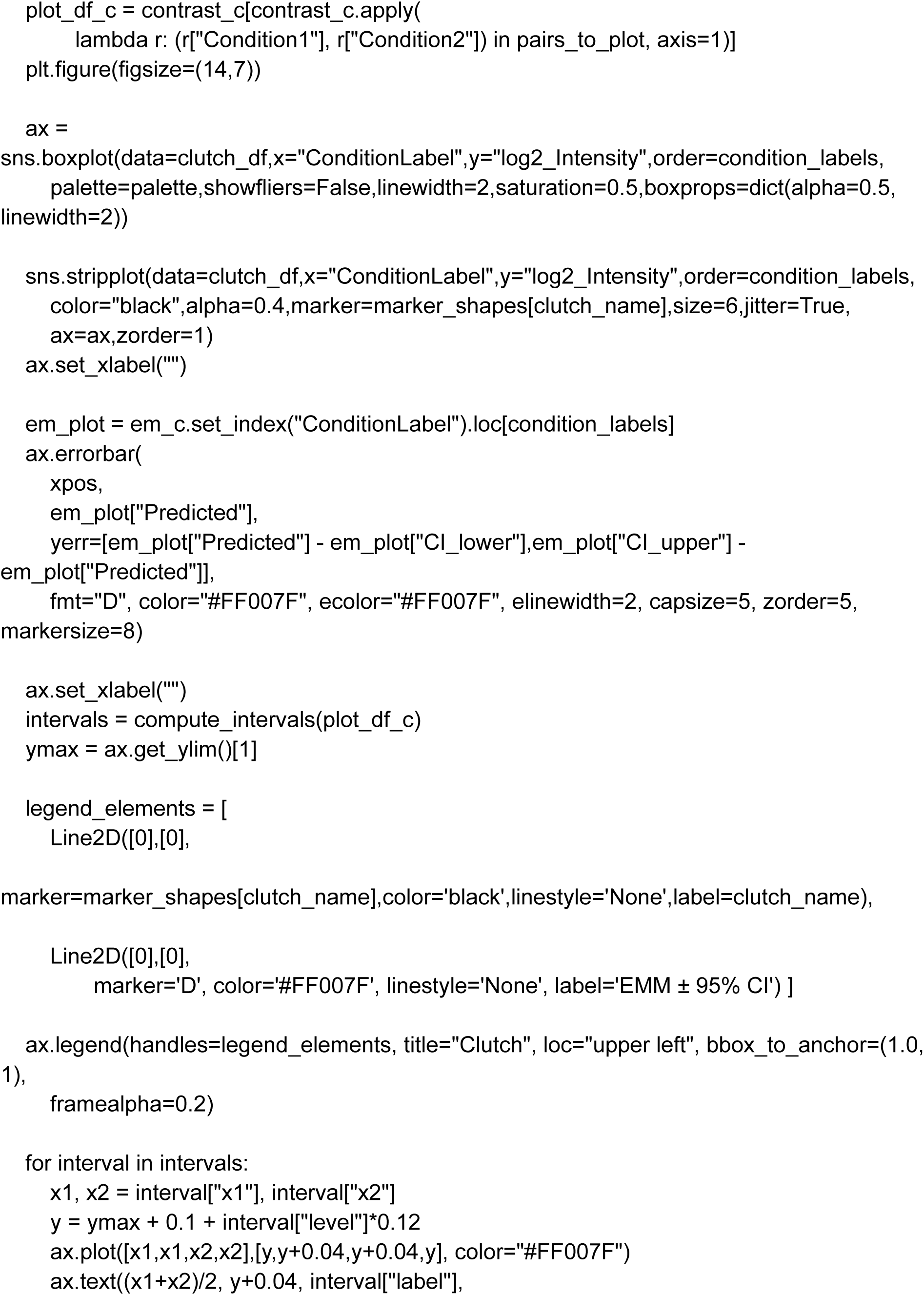
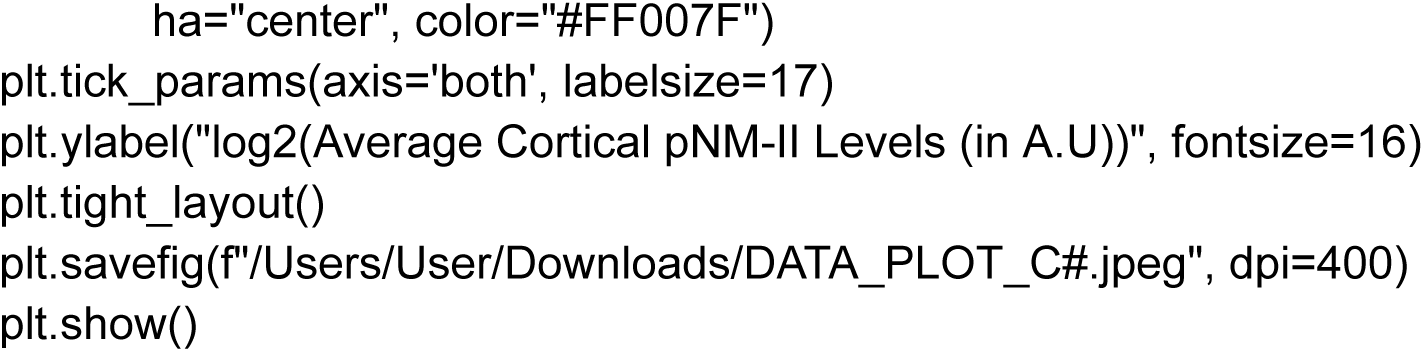

### EMM DIFFERENCE AND FOREST PLOT

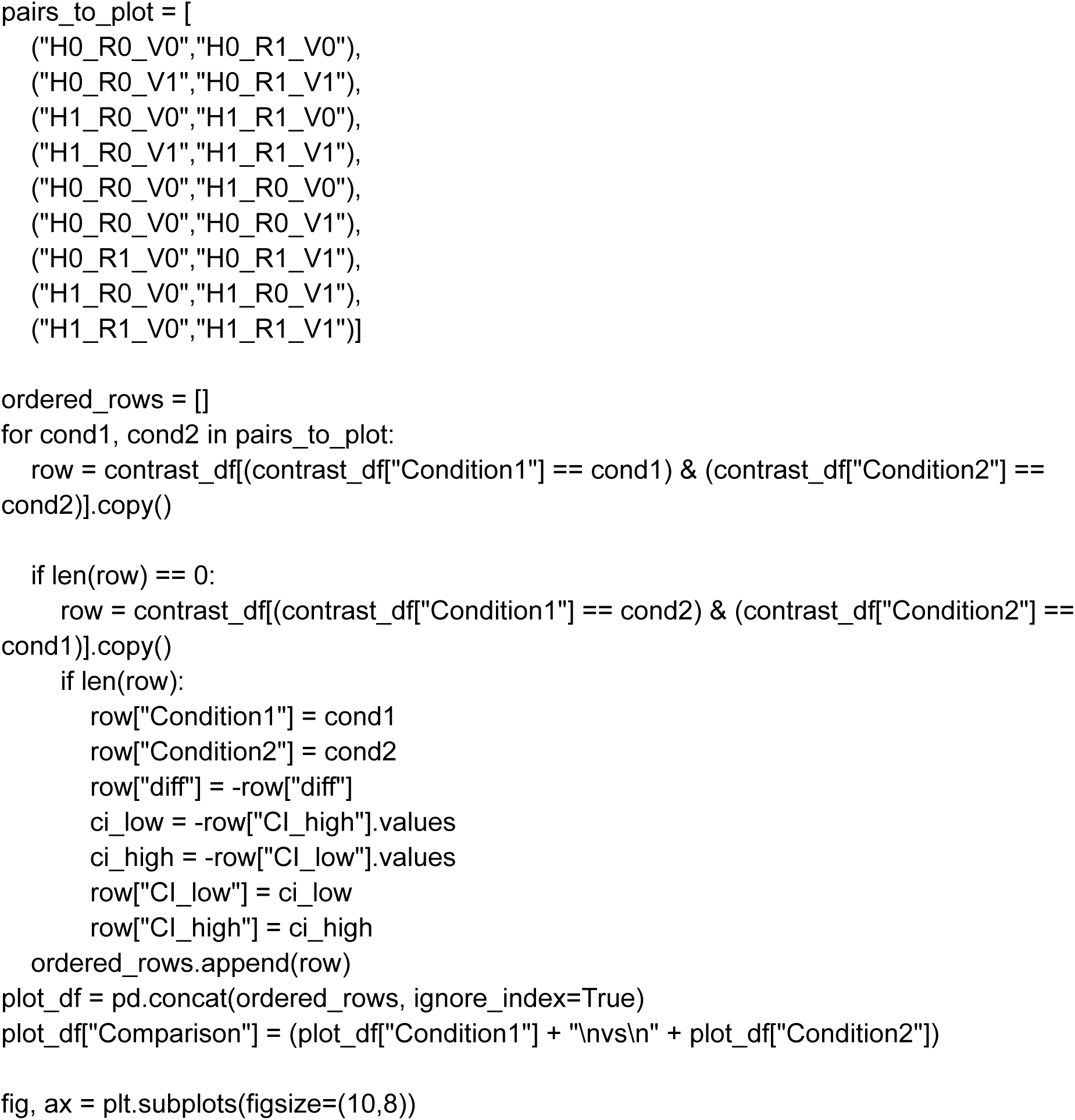
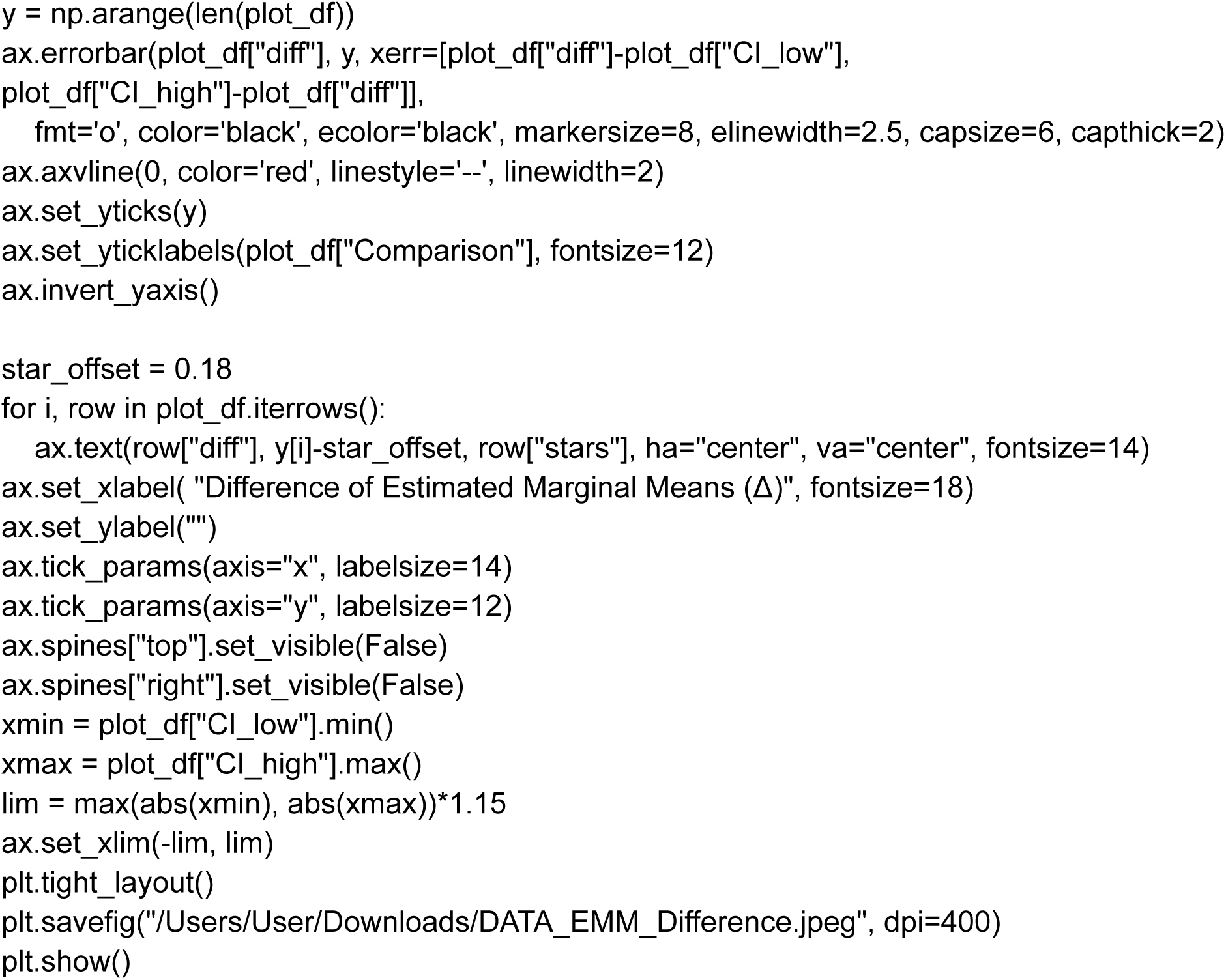

### CODE FOR CROSS-SECTIONAL AREA ANALYSIS

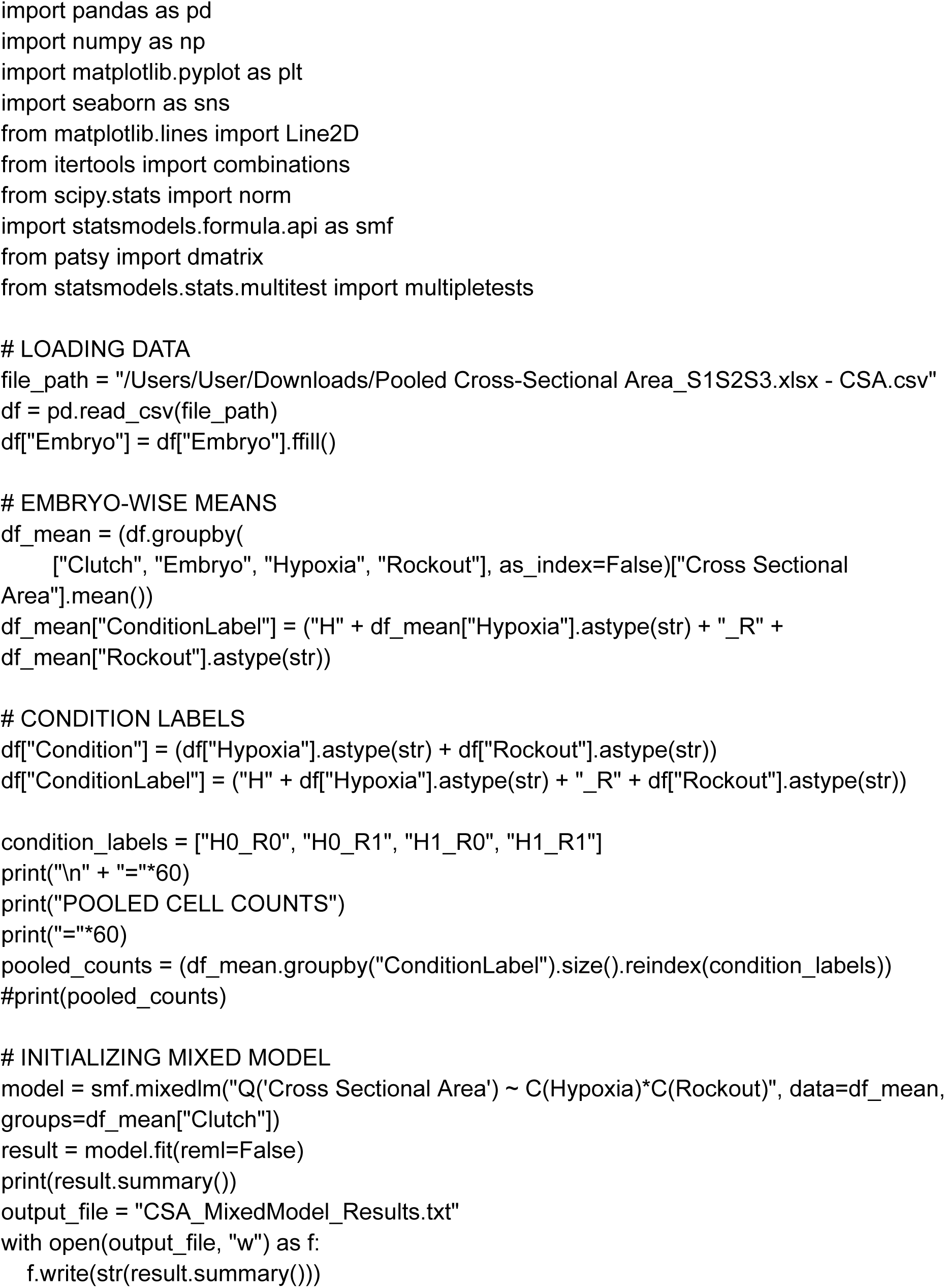
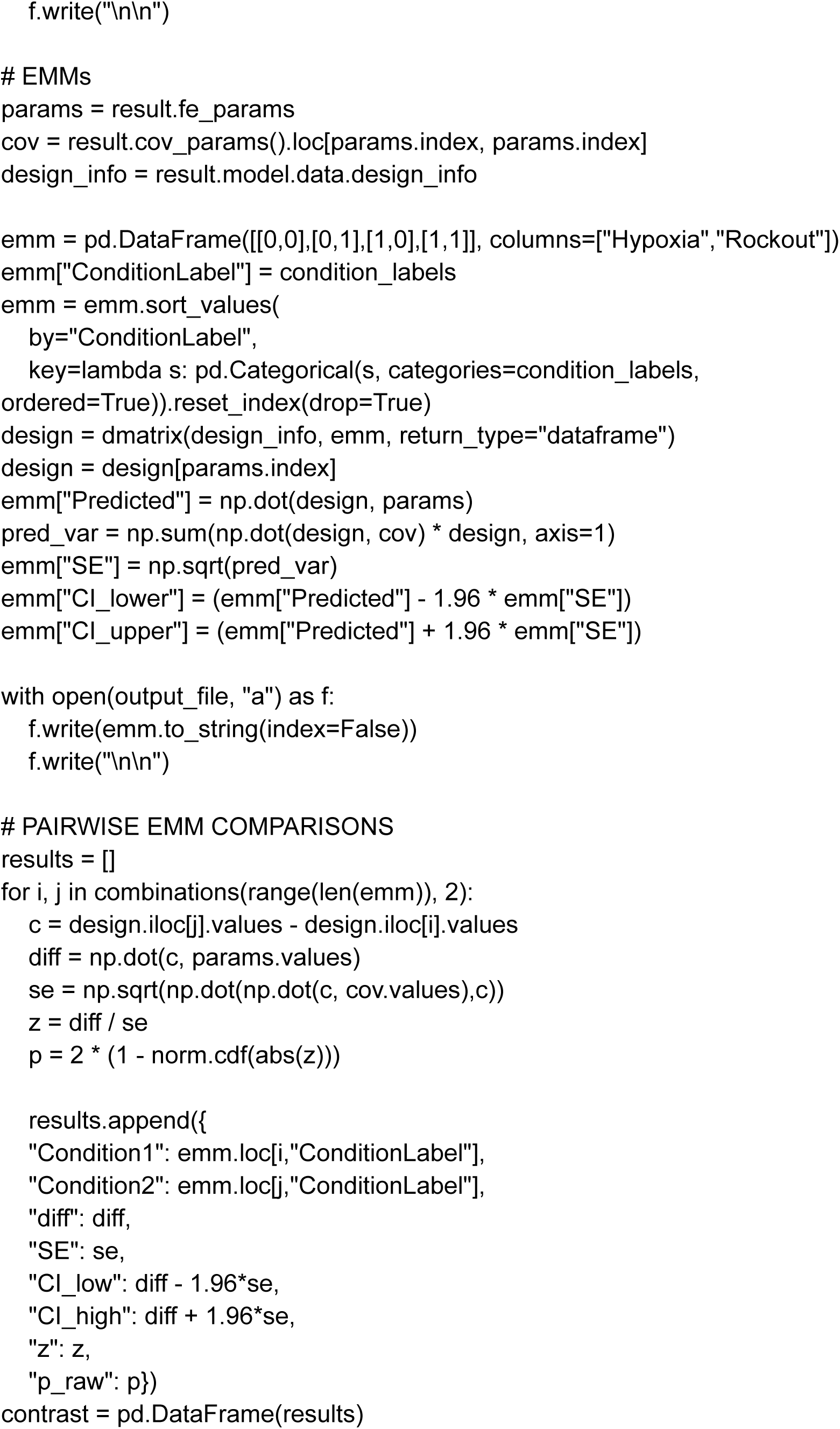
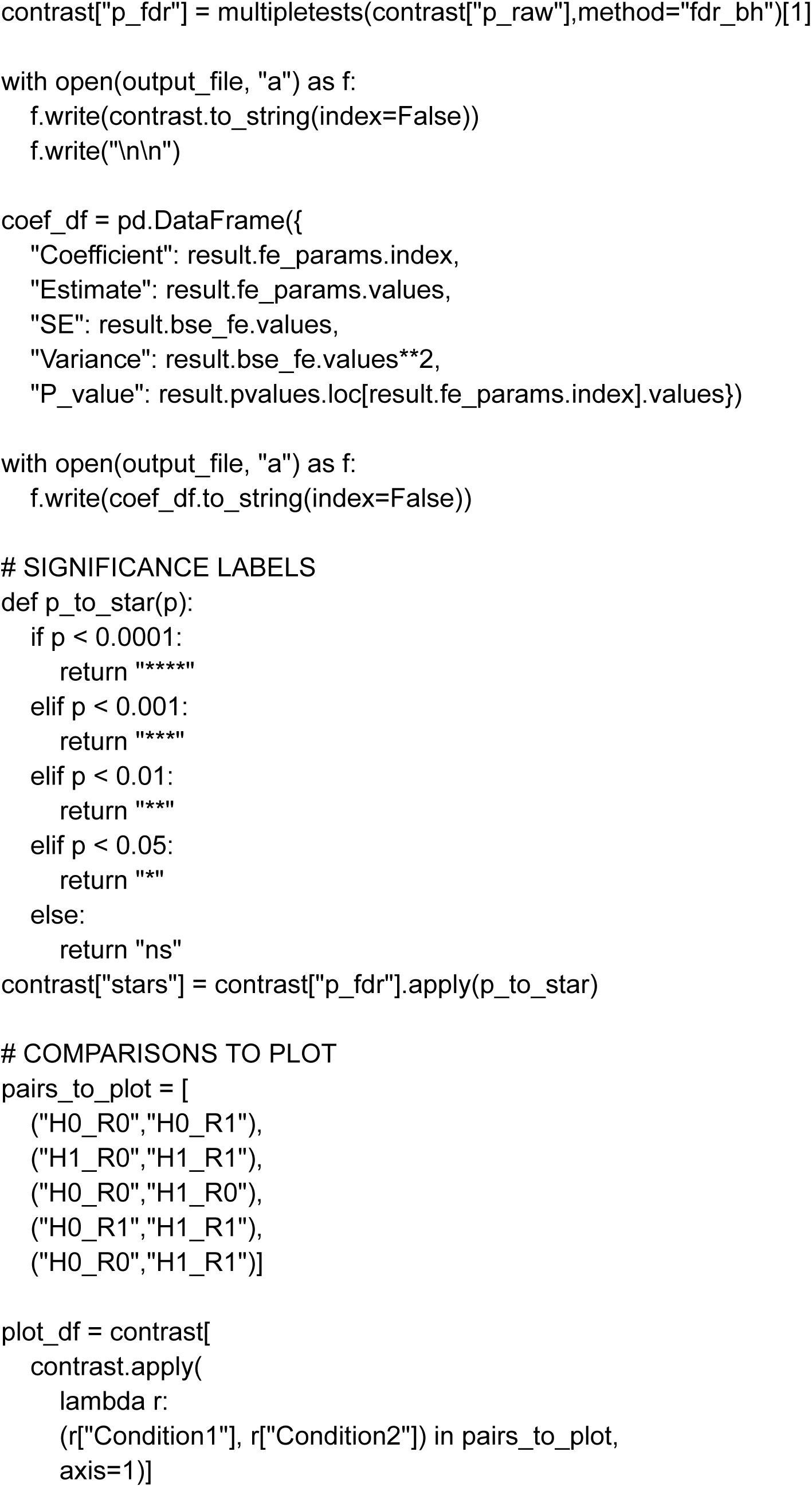
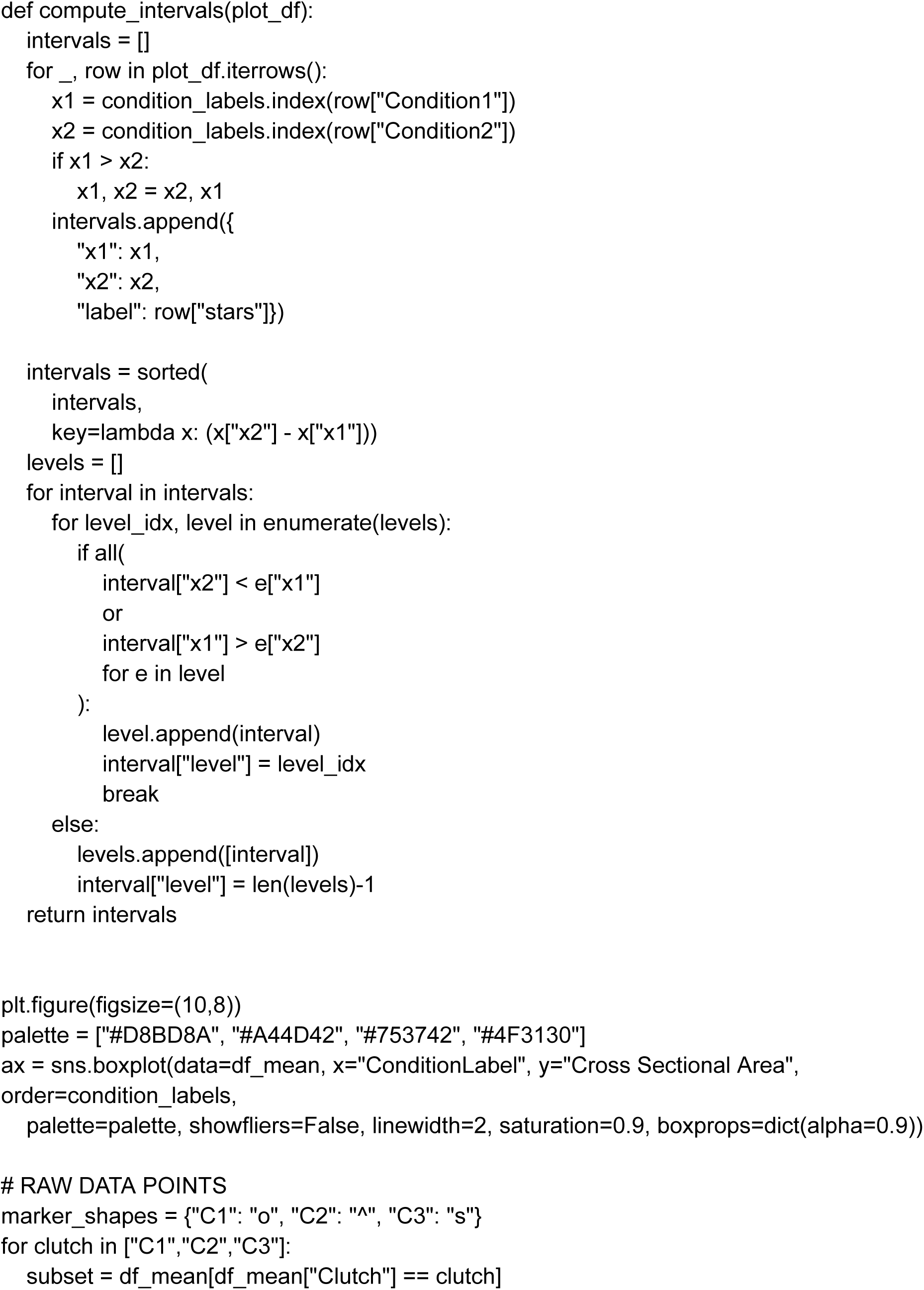
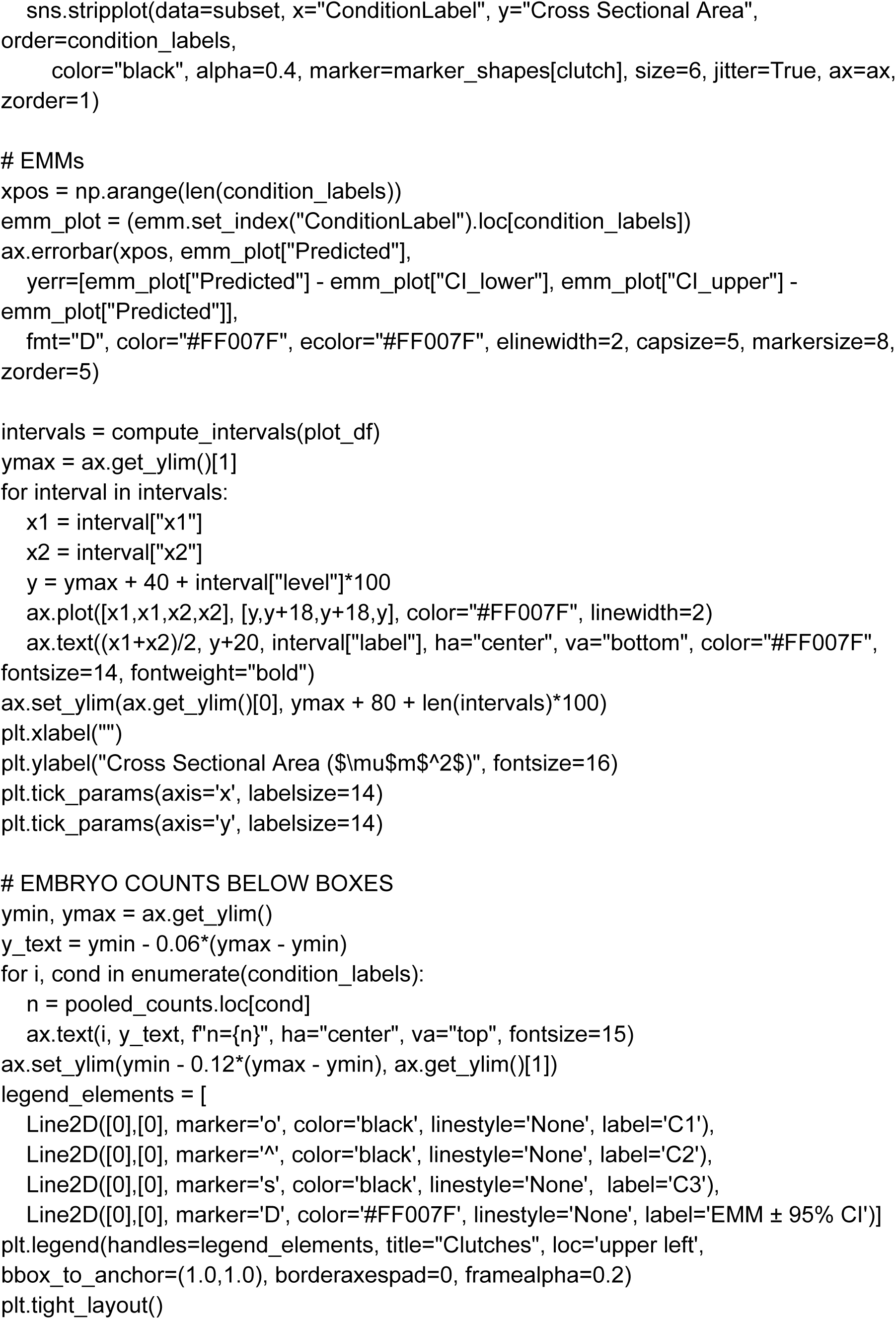
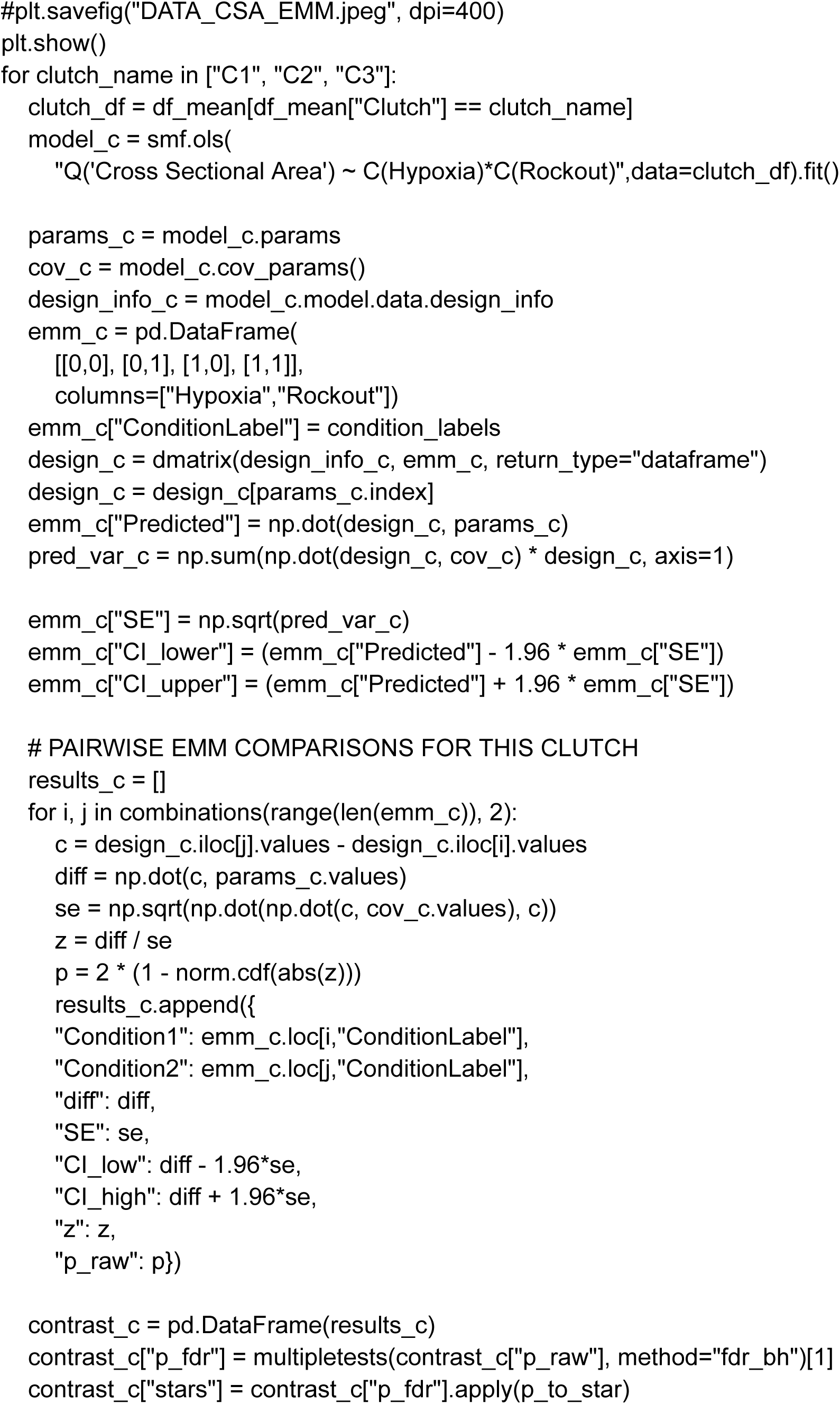
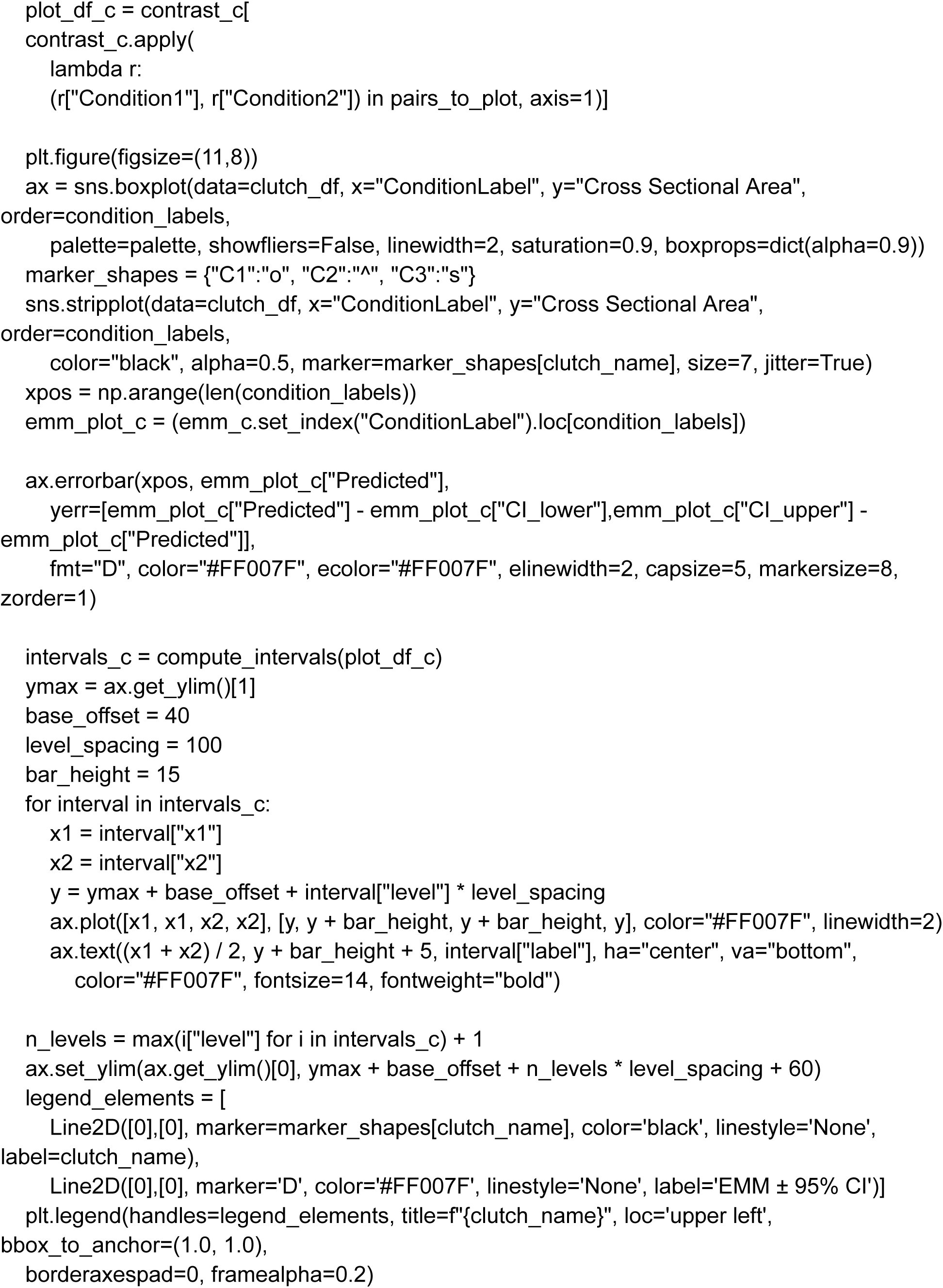
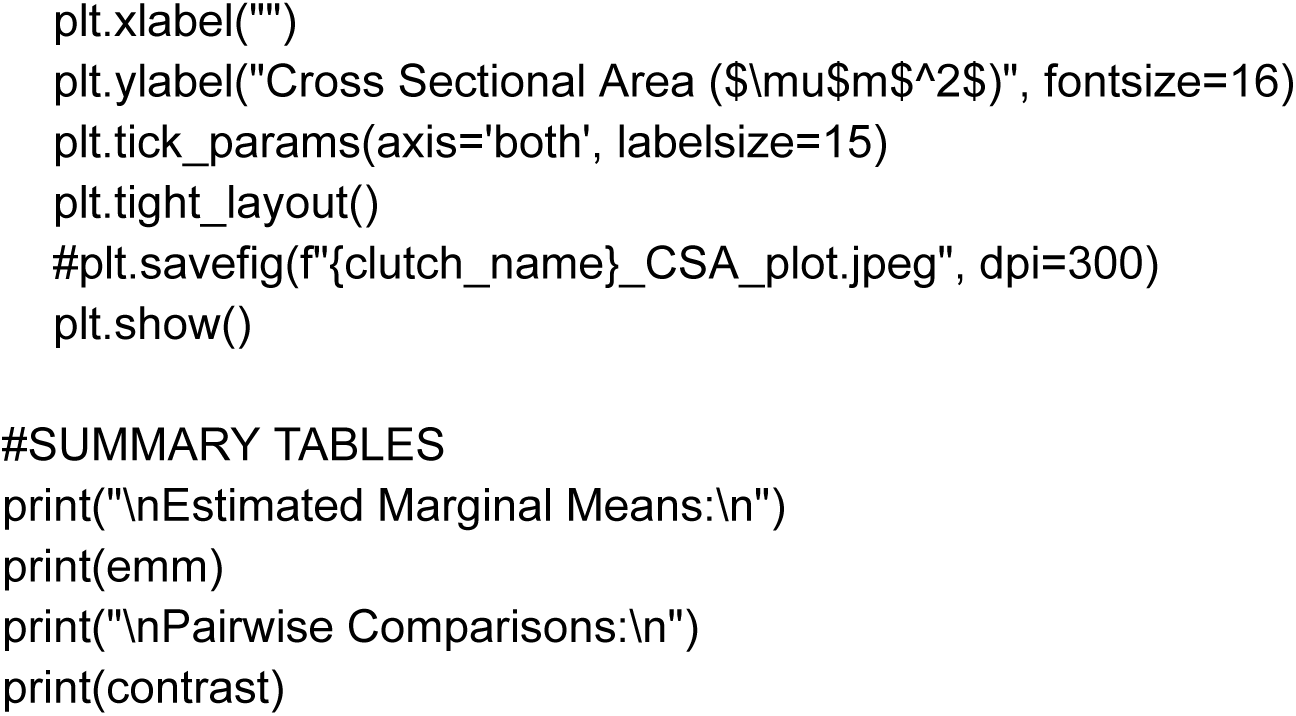

### EMM DIFFERENCE AND FOREST PLOT

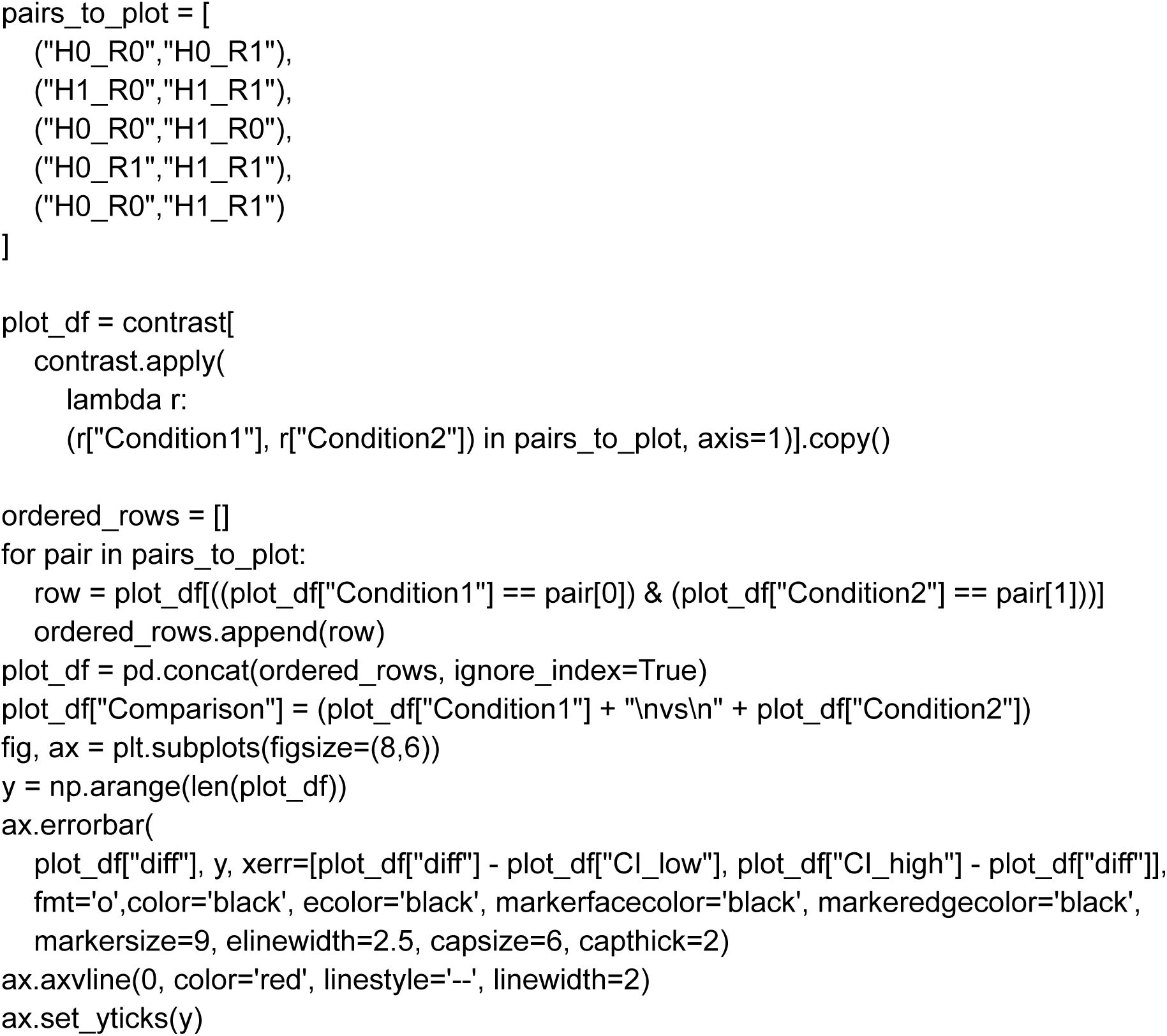
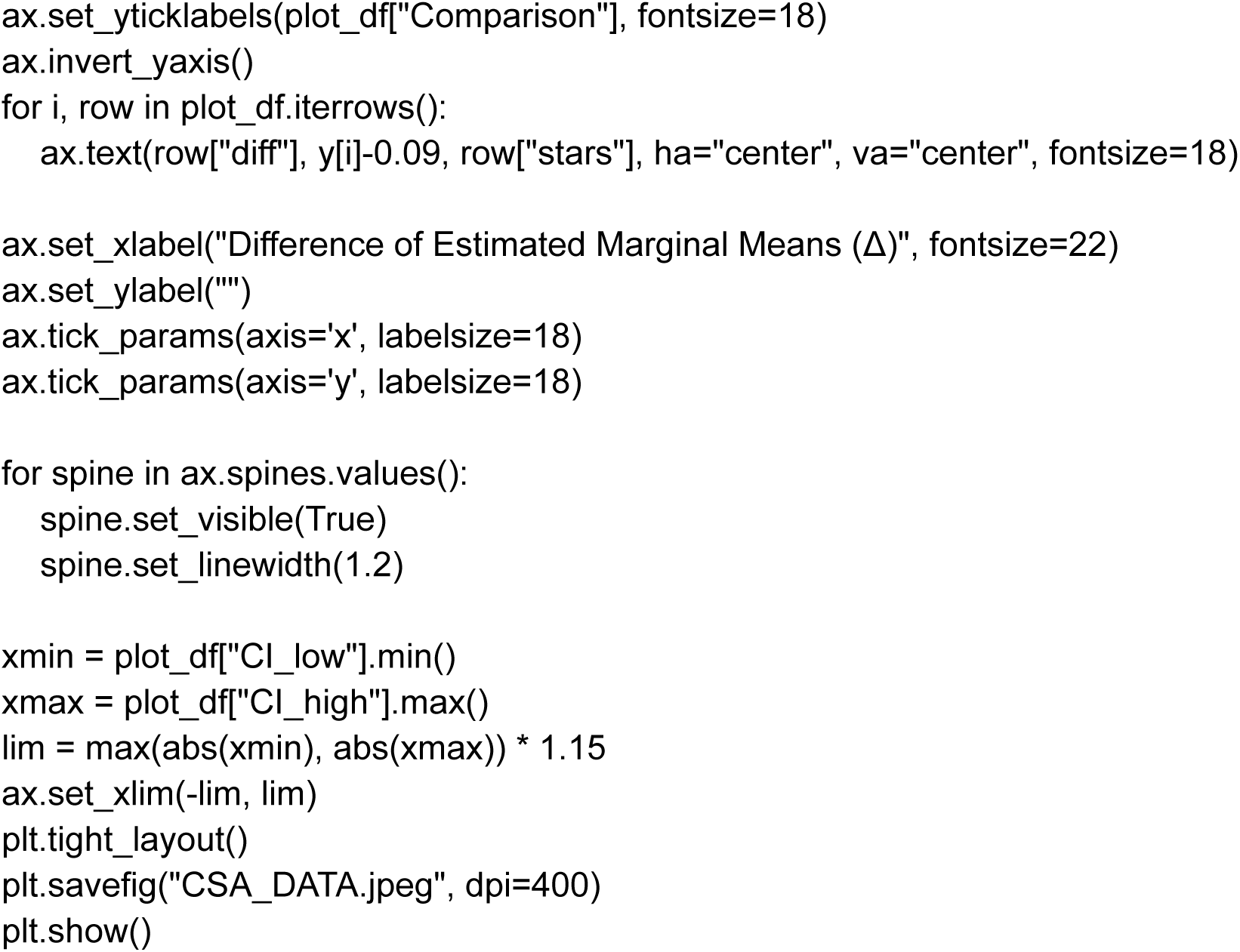

